# Structural robustness and temporal vulnerability of the starvation-responsive metabolic network in liver of healthy and obese mice

**DOI:** 10.1101/2024.06.17.599249

**Authors:** Keigo Morita, Atsushi Hatano, Toshiya Kokaji, Hikaru Sugimoto, Takaho Tsuchiya, Haruka Ozaki, Riku Egami, Dongzi Li, Akira Terakawa, Satoshi Ohno, Hiroshi Inoue, Yuka Inaba, Yutaka Suzuki, Masaki Matsumoto, Masatomo Takahashi, Yoshihiro Izumi, Takeshi Bamba, Akiyoshi Hirayama, Tomoyoshi Soga, Shinya Kuroda

**Author notes:** Correspondence to (S.K.).

## Abstract

Adaptation to starvation is a multi-molecular and temporally ordered process, that could be impaired in obesity. To elucidate how the healthy liver regulates various molecules in a temporally ordered manner during starvation and how obesity disrupts this process, we measured time course multiomic data in the liver of wild-type (WT) and leptin-deficient obese (*ob*/*ob*) mice during starvation. Using the measured data, we constructed a starvation-responsive metabolic network, that is a transomic network including responsive molecules and their regulatory relationships during starvation, and analyzed the structure of the network. In WT mice, ATP and AMP, the energy indicators, regulated various metabolic reactions in the network as the hub molecules, both of which were not responsive in *ob*/*ob* mice. However, the structural properties of the network were maintained in *ob*/*ob* mice. In WT mice, the molecules in the network were temporally ordered through metabolic process coordinated by the hub molecules including ATP and AMP and were positively or negatively co-regulated. By contrast, both temporal order and co-regulation were disrupted in *ob*/*ob* mice. Taken together, the starvation-responsive metabolic network is structurally robust, but temporally vulnerable by the loss of responsiveness of the hub molecules in obesity. In addition, we proposed a potential therapeutic target to treat the negative effects of obesity on intermittent fasting to extend lifespan.

**One Sentence Summary:** Hub molecules activate or inhibit various molecules in a temporally ordered manner in healthy liver, and the regulatory network is structurally robust but temporally vulnerable to obesity.

## Introduction

Adaptation to starvation is important for life to maintain systemic homeostasis, that is a multi-molecular and temporally ordered process to optimize fuel usage (*1–6*). This process involves various molecules, including metabolites, lipids, metabolic enzymes, transcription factors (TFs), and protein kinases. These molecules are temporally ordered across metabolic processes, such as glycogen degradation, gluconeogenesis, and fatty acid degradation. However, how the liver, a central organ for metabolism (*7*, *8*), coordinates various molecules in a temporally ordered manner during starvation at a multiomic scale has been elusive. This is because of the lack of a multiomic measurement at precise time intervals during starvation in mammals, although many researches have conducted omic measurements during starvation at a limited number of time intervals in mammals (*9–19*) or time course omic measurements in microorganisms (*20–23*) or worms (*24*).

Obesity is a pathological status that affects various metabolic processes (*25*– *29*) including adaptation to starvation (*30*), but the global landscape of the effects at a multiomic scale has been elusive. In addition, obesity cancels the beneficial effects on lifespan of intermittent fasting, that is a permissive form of dietary restriction because of an accessible compliance (*31*). The effects of intermittent fasting, which is a long-term and repetitive starvation, result from the cumulative effects of short-term starvation. Thus, clarifying the differences of adaptation to short-term starvation between healthy and obese mice would be informative for investigating an effective therapeutic target of intermittent fasting for obese patients.

Analysis of properties of biological network is a powerful approach to understand such a multi-molecular process like adaptation to starvation (*32–35*). We have developed transomic analysis, an analytical workflow for addressing multi-molecular processes (*36–38*). With this approach, we can integrate multiomic data into a transomic network, offering a global landscape of biochemical reactions. Recently, we showed that analysis of the structural properties of the transomic network aids in their interpretation during oral glucose ingestion (*39*). A network contains of nodes and edges: nodes are entities such as molecules; edges are relationships between nodes such as regulatory events (activation or inhibition). Network structure is how the edges connect nodes. Biological networks, such as metabolic networks (*40–42*), are often scale-free, which means that a small number of “hub” nodes interact with many other nodes, and large number of “leaf” nodes interact with a few other nodes (*43*). We showed that the transomic network is likely scale-free and structural properties of the networks were different between healthy and obese mice during oral glucose ingestion (*39*). However, we have not investigated the relationships between the structural properties and temporal orders of the transomic network, as well as temporal co-regulations, the correlated temporal patterns of nodes in a biological network. Because adaptation to starvation is a temporally ordered multi-molecular process, integrating the structural and temporal analysis of the transomic network would offer mechanistic insights of adaptation to starvation.

In this study, we measured multiomic data, constructed a transomic network, and conducted structural and temporal analyses of the network in the liver of wild-type (WT) and leptin-deficient obese (*ob*/*ob*) mice, a commonly used model of obesity caused by overeating due to loss of leptin (*44*). Through structural analyses, we showed that the starvation-responsive metabolic network is a scale-free-like network and that the structural properties of the network were maintained in *ob*/*ob* mice. We integrated structural and temporal analyses of the transomic network. We found that the network was temporally ordered by hub nodes, such as ATP and AMP, in WT mice, but the temporal order was disrupted in *ob*/*ob* mice, as well as temporal co-regulations. This global feature of the starvation-responsive metabolic network was conserved in glycolysis/gluconeogenesis. However, the structural property of the starvation-responsive metabolic network in urea cycle was disrupted as well as the temporal properties, suggesting the profound disruption of amino acid metabolism in *ob*/*ob* mice during starvation. Our omic data was also informative for the study of dietary restriction. Taken together, the starvation-responsive metabolic network was well-structured, temporally ordered, and co-regulated in healthy liver. The network was structurally robust but temporally vulnerable to obesity.

The focus here is adaptive responses to starvation, rather than direct comparisons between WT and *ob*/*ob* mice in a steady state. We have conducted such comparative studies previously for fasting and fed states (*45*, *46*). Note that identical data of 0 h starvation have been already published in one of our studies (*46*).

## Results

### Workflow for the construction and analysis of a global starvation-responsive transomic network in liver of WT and *ob*/*ob* mice

To describe the temporal dynamics of various molecules during adaptation to starvation in healthy liver and pathological alterations caused by obesity, we constructed and analyzed a global starvation-responsive transomic network in liver of WT and *ob*/*ob* mice (Fig. 1). We collected multiomic data from mice starved for 0, 2, 4, 6, 8, 12, 16, and 24 hours (Fig. 1**A**). We measured the metabolome, lipidome, FFA & acyls (free fatty acids, acyl-carnitines, and acyl-CoAs), transcriptome, proteome, and phosphoproteome in liver. We also measured the metabolome in plasma.

**Fig. 1:**
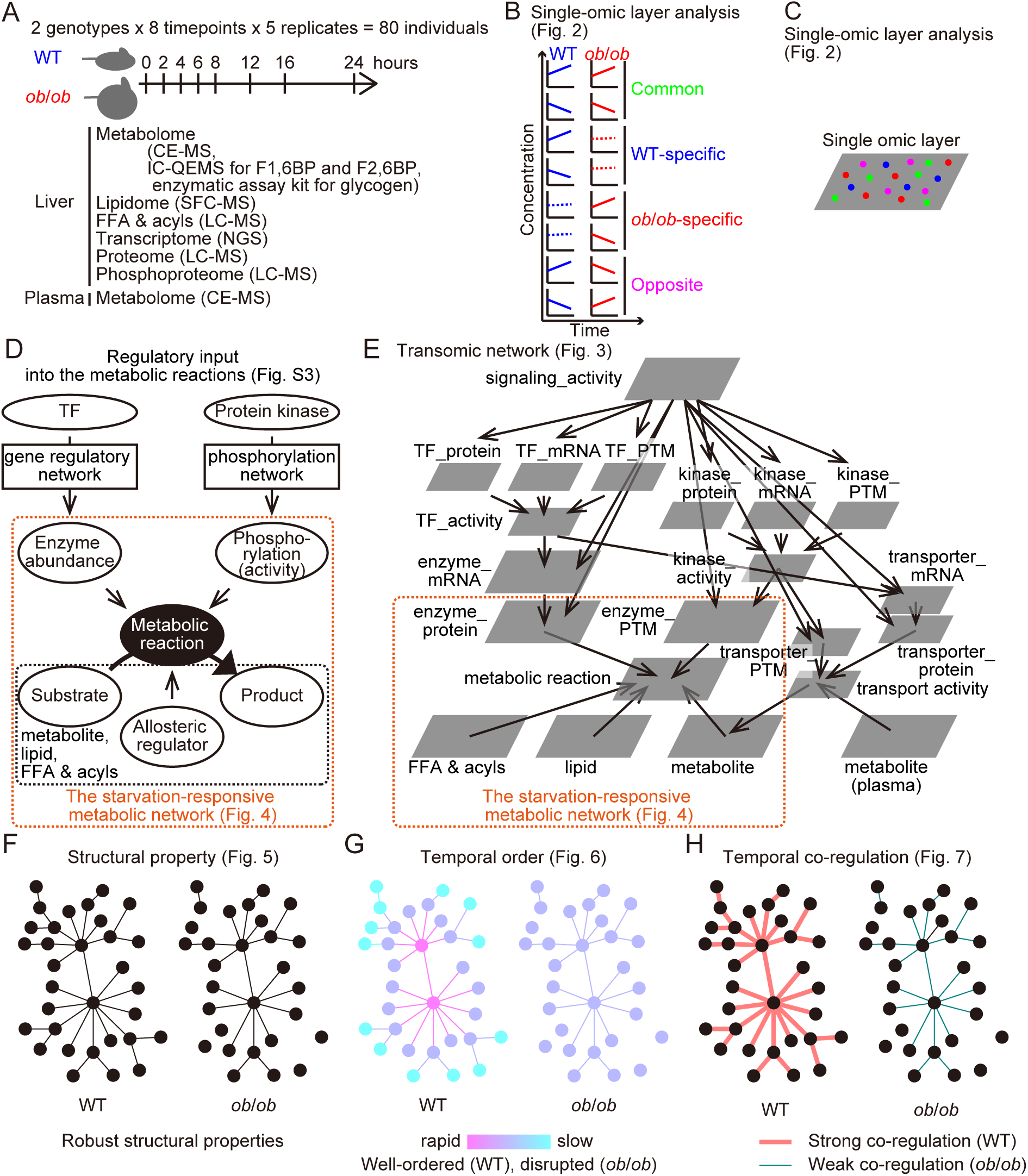
Workflow for the construction and analysis of a global starvation-responsive transomic network in liver of WT and *ob*/*ob* mice (**A**) Experimental design. WT and *ob*/*ob* mice were starved for the indicated hours, followed by the collection of liver and plasma samples. These samples were subjected to multiomic measurements. It should be noted that metabolome was basically measured by CE-MS, but IC-QEMS was used for quantification of F1,6BP and F2,6BP. Enzymatic assay kit was used for glycogen. (**B**) Classification of molecules into 4 groups: commonly responsive in both WT and *ob*/*ob* mice (common, green); WT-specifically (WT-specific, blue); *ob*/*ob*-specifically (*ob/ob*-specific, red); oppositely (opposite, magenta) responsive molecules between WT mice and *ob*/*ob* mice. (**C**) Schema for a single-omic layer composed of responsive molecules classified as in (**B**). (**D**) Schema of regulatory input into the metabolic reactions. TFs increase or decrease metabolic enzyme abundance through gene regulation. Protein kinases increase or decrease enzyme phosphorylation to activate or inhibit enzyme activity. The metabolites, FFA & acyls, and lipids act as substrates, products, and allosteric regulators in metabolic reactions. The subset of the network that is defined as the starvation-responsive metabolic network is outlined in orange. (**E**) Schematic representation of the global starvation-responsive transomic network consisting of the indicated layers. The subset of the network that is defined as the starvation-responsive metabolic network is outlined in orange. (**F**) Structural properties of the starvation-responsive metabolic network in WT and *ob/ob* mice. (**G**) Temporal order of the starvation-responsive metabolic network in WT and *ob*/*ob* mice. (**H**) Temporal co-regulations among the time courses molecules in the starvation-responsive metabolic network in WT and *ob*/*ob* mice.

Using the collected data, we conducted a single-omic layer analysis (Fig. 1**B**, and **C**). We identified starvation-responsive molecules and classified them into 4 groups in each layer (Fig. 1**B**): “common,” those that exhibited the same response in both WT and *ob*/*ob* mice; “WT-specific,” those that occurred only in WT mice; “*ob*/*ob*-specific,” those that occurred only in *ob*/*ob* mice; and “opposite,” those that exhibited opposing responses between WT and *ob*/*ob* mice. These results formed a single omic layer (Fig. 1**C**). Note that phosphorylation is a process occurred on a phosphosite, not a molecule, but we refer phosphorylation events as molecules as well as metabolites, lipids, FFA & acyls, proteins, and transcripts for simplicity.

The starvation-responsive molecules cooperatively activate or inhibit metabolic reactions to adapt to starvation across the multiomic layers (Fig. 1**D**). Metabolites, lipids, and FFA & acyls serve as substrates, products, and allosteric regulators for metabolic reactions. Enzyme abundance and phosphorylation activate or inhibit the metabolic reactions. TFs increase or decrease enzyme abundance through the gene regulatory network, although the control is partial because of the effects of protein translation and degradation. Protein kinases increase or decrease enzyme phosphorylation through kinase-substrate network to regulate enzyme activity. Using bioinformatic platforms, we integrated single omic layers along the regulatory network of metabolic reactions and constructed a global starvation-responsive transomic network (Fig. 1**E**). In this manuscript, we refer to “regulation” as encompassing both activation and inhibition when no specific regulatory relationship is mentioned, for simplicity.

Metabolic enzymes, metabolites, lipids, and FFA & acyls, are direct, major and fine regulators of metabolism (*37*, *47*, *48*). Therefore, we focused on the starvation-responsive metabolic network, that is the sub-network of the global starvation-responsive transomic network (Fig. 1**D** and **E**, dotted orange outline) to investigate structural and temporal properties of the network. The starvation-responsive metabolic network contains the direct regulators of metabolic reactions: metabolite, lipid, FFA & acyls, enzyme_protein (protein abundances of metabolic enzymes), and enzyme_PTM (phosphorylation of metabolic enzymes). We investigated structural properties of this starvation-responsive metabolic network (Fig. 1**F**), the temporal order of changes in the network (Fig. 1**G**), and temporal co-regulation among molecules (Fig. 1**H**).

### Single-omic layer analysis: identification of starvation-responsive molecules and their regulators

We measured seven omic layers: metabolome, lipidome, FFA & acyls, transcriptome, proteome, and phosphoproteome in the liver and metabolome in plasma (see fig. S1 for representative examples, and fig. S2 and Data File S1 for complete data) and performed a single-omic layer analysis in WT and *ob*/*ob* mice (Fig. 2). We conducted one-way ANOVA tests (ANOVA-like testing of edgeR (*49–51*) for transcriptome) in WT and *ob*/*ob* mice respectively, and defined molecules that showed a *P* value less than 0.025 as starvation-responsive molecules (Fig. 2**A**; Data File S2), whose false discovery rates were less than 0.1 except for phosphoproteome data (Table 1). Based on the area under the curve (AUC) of the log2-transformed time courses (Fig. 2**B**), starvation-responsive molecules with positive AUCs were classified as increased molecules (orange), and those with negative AUCs were classified as decreased molecules (purple). Using these classifications, we defined four classes of molecules: common (changing similarly in both WT and *ob*/*ob* mice, green), WT-specific (blue), *ob*/*ob*-specific (red), and opposite (changing in opposite directions in WT and *ob*/*ob* mice, magenta) (Fig. 2**C**).

**Fig. 2:**
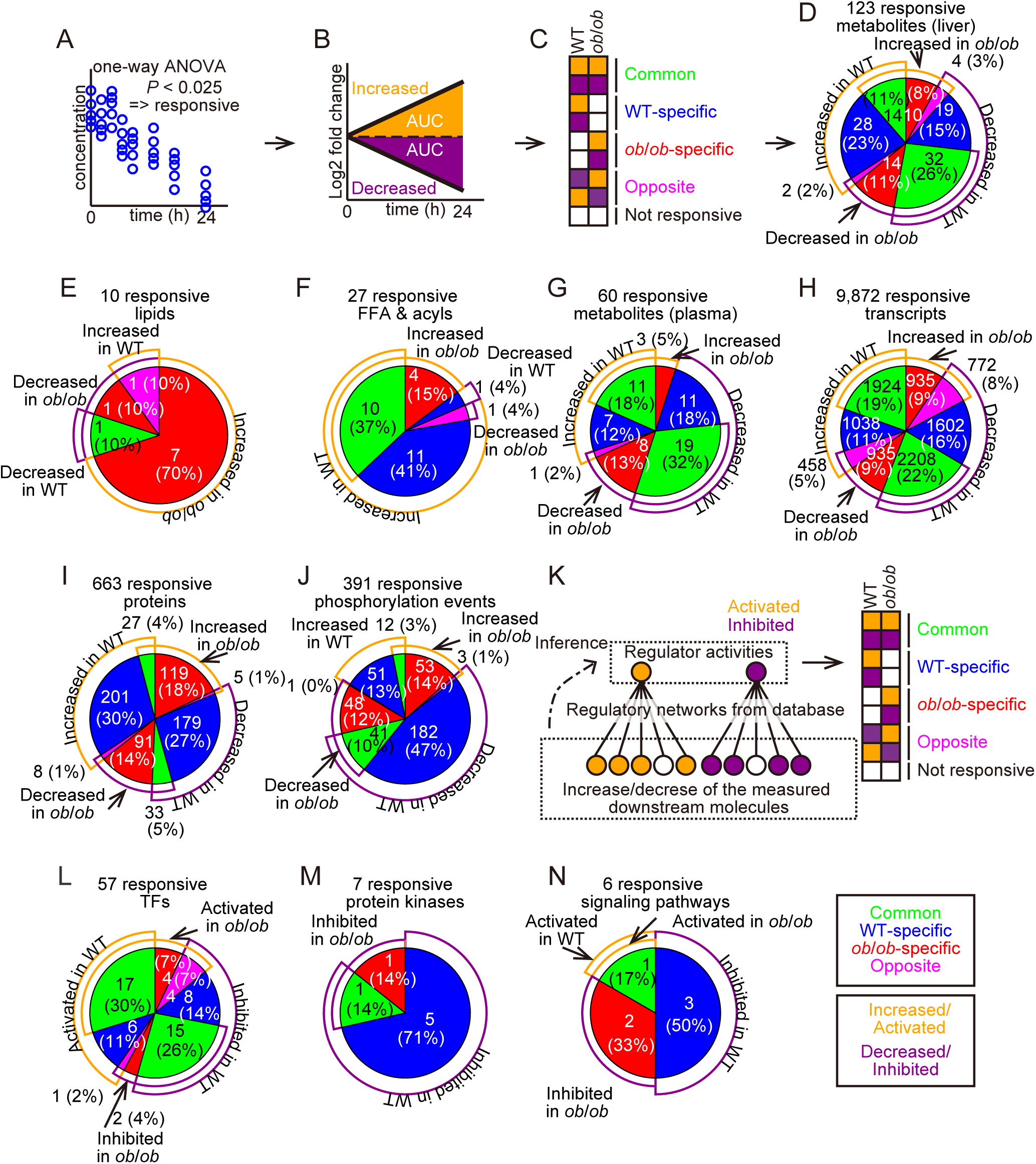
Single-omic layer analysis during starvation (**A**-**C**) Identification and classification of starvation-responsive molecules. (**A**) One-way ANOVA tests (*P* < 0.025) defined starvation-responsive molecules. (**B**) Area under the curve (AUC) of log2-transformed mean time courses classified molecules into increased (AUC > 0, orange) and decreased (AUC < 0, purple) classes. (**C**) Classification of starvation-responsive molecules: common in both WT and *ob*/*ob* (either increased or decreased in both, green), WT-specific (blue), *ob*/*ob*-specific (red), or opposite (increased in WT and decreased in *ob*/*ob*, or vice versa, magenta). (**D**-**J**) The number of molecules and distribution in each group shown as pie charts colored as defined in (**C**). Increased or decreased molecules are indicated by the orange or purple arcs, respectively. The number and the ratio of significantly changed molecules of the molecules in each category are shown. Responsive metabolites (**D**), lipids (**E**), and FFA & acyls (**F**) in liver. Responsive plasma metabolites (**G**). Responsive transcripts (**H**), proteins (**I**), and phosphorylation events (**J**) in liver. (**K**) Strategy for inferring activities of upstream regulators and signaling pathways. Briefly, we constructed regulatory networks using database and identified increased and decreased downstream molecules from the measured data. In the case of positive regulators, we assumed that a regulator is activated if most of the downstream molecules increased and vice versa (see Methods). We screened regulators whose inferred activities were consistent with the measured data (see Methods). Activated or inhibited regulators were classified into common (green), WT-specific (blue), *ob*/*ob*-specific (red), or opposite (magenta) groups. (**L**-**N**) The number and distribution of these inferred TFs (**L**), protein kinases (**M**), or signaling pathways (**N**) are shown in pie charts. *n* = 5 biological replicates per group.

**Table 1.** Maximum *Q*-values for the omic layers of the starvation-responsive molecules. Maximum *Q*-values for each omic layer for molecules with *P*-values less than 0.025. All the maximum *Q*-values were less than 0.1 except for phosphoproteome, suggesting that false discovery rates of the starvation-responsive molecules were less than 0.1 in all the omic layers except for phosphoproteome.

| Omics type | Maximum $Q$-value |
| --- | --- |
| Metabolome (Liver) | 0.0029 |
| Transcriptome | 0.0489 |
| Proteome | 0.0661 |
| Phosphoproteome | 0.1645 |
| Lipidome | 0.0502 |
| FFA & acyls | 0.053 |
| Metabolome (Plasma) | 0.0016 |

We identified starvation-responsive molecules in the single-omic layer analyses of metabolites (Fig. 2**D**), lipids (Fig. 2**E**; Table 2), FFA & acyls (Fig. 2**F**), plasma metabolites (Fig. 2**G**), transcripts (Fig. 2**H**), proteins (Fig. 2**I**), and phosphorylation events (Fig. 2**J**). The common group comprised more than one-third of the total responsive metabolites in both liver (Fig. 2**D**) and plasma (Fig. 2**G**), FFA & acyls (Fig. 2**F**), and transcripts (Fig. 2**H**). Most starvation-responsive lipids were in the *ob*/*ob*-specific group (Fig. 2**E**). More than half of responsive proteins (Fig. 2**I**) and phosphorylation events (Fig. 2**J**) were WT-specifically responsive.

**Table 2.** Measured lipid classes Measured lipid classes and their abbreviations.

| Common name | Abbreviation |
| --- | --- |
| Lysophosphatidylcholine | LPC |
| Lysophosphatidylethanolamine | LPE |
| Cholesterol | Chol |
| Phosphatidylethanolamine | PE |
| Triacylglycerol | TG |
| Ceramide | Cer |
| Diacylglycerol | DG |
| Monoacylglycerol | MG |
| Phosphatidic acid | PA |
| Phosphatidylcholine | PC |
| Phosphatidylglycerol | PG |
| Phosphatidylinositol | PI |
| Phosphatidylserine | PS |
| Sphingomyelin | SM |
| Cholesterol ester | CE |
| Hexose ceramide | HexCer |

KEGG pathway enrichment analysis (Data File S3) revealed pathways that were associated with the starvation-responsive changes in both the transcriptome (Table 3) and proteome (Table 4). For example, KEGG pathways such as “fatty acid degradation” and “PPAR signaling pathway” were enriched in both transcriptome (Table 3) and proteome (Table 4) in WT mice. However, this analysis did not find any pathways that were enriched in the phosphoproteome data (Data File S3).

**Table 3.** KEGG pathway enrichment analysis of the starvation-responsive transcripts KEGG pathways with *Q*-value < 0.1 are shown.

|  | WT | *ob/ob* |
| --- | --- | --- |
| Increased | Fatty acid degradation, Lysine degradation, Glycerolipid metabolism, Biosynthesis of unsaturated fatty acids, PPAR signaling pathway, Peroxisome, Fat digestion and absorption | PPAR signaling pathway, Fat digestion and absorption |
| Decreased | Pentose and glucuronate interconversions, Ascorbate and aldarate metabolism, Steroid biosynthesis, Steroid hormone biosynthesis, Tyrosine metabolism, Selenocompound metabolism, Glutathione metabolism, Thiamine metabolism, Retinol metabolism, Porphyrin metabolism, Terpenoid backbone biosynthesis, Sulfur metabolism, Metabolism of xenobiotics by cytochrome P450, Drug metabolism - cytochrome P450, Drug metabolism - other enzymes, ABC transporters, Protein export, Mismatch repair, Protein processing in endoplasmic reticulum, Necroptosis, Cellular senescence, Signaling pathways regulating pluripotency of stem cells, RIG-I-like receptor signaling pathway, Serotonergic synapse, Prolactin signaling pathway, Bile secretion | Glycolysis / Gluconeogenesis, Pentose phosphate pathway, Fructose and mannose metabolism, Fatty acid biosynthesis, Steroid biosynthesis, Steroid hormone biosynthesis, Glycine, serine and threonine metabolism, Tryptophan metabolism, Selenocompound metabolism, Glutathione metabolism, Starch and sucrose metabolism, Amino sugar and nucleotide sugar metabolism, Pyruvate metabolism, Glyoxylate and dicarboxylate metabolism, One carbon pool by folate, Nicotinate and nicotinamide metabolism, Retinol metabolism, Porphyrin metabolism, Terpenoid backbone biosynthesis, Sulfur metabolism, Aminoacyl-tRNA biosynthesis, Metabolism of xenobiotics by cytochrome P450, Drug metabolism - cytochrome P450, Drug metabolism - other enzymes, Biosynthesis of unsaturated fatty acids, ABC transporters, DNA replication, Protein export, Base excision repair, Mismatch repair, Non-homologous end-joining, Protein processing in endoplasmic reticulum, Peroxisome, AMPK signaling pathway, Longevity regulating pathway, Thermogenesis, Phototransduction, Insulin signaling pathway |

**Table 4.** KEGG pathway enrichment analysis of the starvation-responsive proteins KEGG pathways with *Q*-value < 0.1 are shown.

|  | WT | *ob/ob* |
| --- | --- | --- |
| Increased | Citrate cycle (TCA cycle), Fatty acid degradation, Arginine biosynthesis, Valine, leucine and isoleucine degradation, Phenylalanine metabolism, Phenylalanine, tyrosine and tryptophan biosynthesis, beta-Alanine metabolism, Propanoate metabolism, PPAR signaling pathway | Ribosome, Lysosome |
| Decreased | Pentose and glucuronate interconversions, Ascorbate and aldarate metabolism, Steroid hormone biosynthesis, Glutathione metabolism, One carbon pool by folate, Retinol metabolism, Porphyrin metabolism, Metabolism of xenobiotics by cytochrome P450, Drug metabolism - cytochrome P450, Drug metabolism - other enzymes, Protein processing in endoplasmic reticulum, Antigen processing and presentation, Serotonergic synapse | No enrichment |

We inferred changes in the activities of following regulators of the starvation-responsive molecules (Fig. 2**K**): TFs (Fig, 2**L**; Table 5; Data File S4), protein kinases (Fig. 2**M**; Table 6; Data File S4), and signaling pathways (Fig. 2**N**; Table 7) based on the regulatory network obtained from bioinformatic platforms and measured omic data. Note that signaling pathway layer (Fig. 2**N**; Table 7) contains significantly enriched signaling pathways in any of the transcriptome (Table 3) or proteome data (Table 4). We classified the activities of these regulators as common, WT-specific, *ob*/*ob*-specific, or opposite (Fig. 2**K**). The common group comprised more than half of the total responsive TFs (Fig. 2**L**; Table 5), including Pparα and Hnf4α, well-established for promoting lipid degradation (*5*). Note that some circadian TFs are included beacause of sampling time and some results were not consistent with the previous studies such as failure of identification of Foxo1 and unexpected identification of Creb1 as an inhibited TF. These problems were caused by insufficient accuracy and comprehensiveness of the database and omic data, particularly the limited number of quantified proteins and phosphorylation sites. “PKC_group” kinases including Prkce, Prkcd and Prkcz, master regulators of liver metabolism (*52–54*), were commonly inhibited (Fig. 2**M**; Table 6). The “PPAR signaling pathway” was inferred as an activated signaling pathway (Fig. 2**N**; Table 7) commonly in WT and *ob*/*ob* mice.

**Table 5.** Activated or inhibited TFs during starvation Activated or inhibited TFs during starvation. ‘Up’, ‘Down’, and ‘NE’ represent activation, inhibition, and no enrichment, respectively.

| WT | *ob/ob* | TFs |
| --- | --- | --- |
| Up | Up | Arntl, Clock, Cry1, Egr1, Ep300, Foxa3, Hnf4a, Myc, Nelfa, Nfil3, Ppara, Prox1, Rorc, Stat3, Stat5a, Tead1, Yap1 |
| Up | NE | Cebpb, Crebbp, Hdac3, Med1, Rad21, Rora |
| Up | Down | Klf15 |
| NE | Up | Hsf1, Nr1h4, Pbrm1, Rela |
| NE | Down | Gps2, Gtf2e2 |
| Down | Up | Bcl6, Foxa1, Gata4, Ncoa3 |
| Down | NE | Ctbp2, Cux2, Foxa2, Nr1h3, Nr3c1, Onecut1, Rxra, Tead4 |
| Down | Down | Ahr, Cebpa, Creb1, Dbp, E2f1, Esr1, Mlxipl, Ncor1, Nfe2l2, Nr1d1, Nr1d2, Pparg, Srebf1, Ssu72, Xbp1 |

**Table 6:** Activated or inhibited protein kinase groups during starvation Activated or inhibited protein kinase groups during starvation. ‘Up’, ‘Down’, and ‘NE’ represent activation, inhibition, and no enrichment, respectively.

|  |  |  |
| --- | --- | --- |
| WT | *ob/ob* | Protein kinases |
| Up | Up | NE |
| Up | NE | NE |
| Up | Down | NE |
| NE | Up | NE |
| NE | Down | NEK1_NEK5_NEK3_NEK4_NEK11_NEK2_group |
| Down | Up | NE |
| Down | NE | CaMKII_group, PAK_group, PDHK_group, PKD_group, SLK_group |
| Down | Down | PKC_group |

**Table 7:** Activated or inhibited signaling pathways during starvation Activated or inhibited signaling pathways during starvation. ‘Up’, ‘Down’, and ‘NE’ represent activation, inhibition, and no enrichment, respectively.

| WT | *ob/ob* | Signaling pathways |
| --- | --- | --- |
| Up | Up | PPAR signaling pathway |
| Up | NE | NE |
| Up | Down | NE |
| NE | Up | NE |
| NE | Down | AMPK signaling pathway,<br>Insulin signaling pathway |
| Down | Up | NE |
| Down | NE | Prolactin signaling<br>pathway, RIG-I-like<br>receptor signaling pathway,<br>Signaling pathways<br>regulating pluripotency of<br>stem cells |
| Down | Down | NE |

### Construction of the global starvation-responsive transomic network by integrating multiomic data

Using the starvation-responsive molecules and their regulators (Fig. 2), we constructed a global starvation-responsive transomic network (Fig. 3**A**; Data File S5). The network consists of nodes, layers, and edges. Nodes represent measured molecules (Figs 2**D**-**J**), inferred activities of the TFs, protein kinases, and signaling pathways (Figs. 2**L**, **M**, and **N**), metabolic reactions defined using KEGG (*55–57*), and transport activities from TCDB (*58*). Based on chemical properties, biological functions, or both, nodes were clustered into layers (see Table 8 for the precise definition). For example, the enzyme_protein layer includes proteins (chemical property) that are metabolic enzymes (biological functions). Enzyme_PTM layer represents phosphorylation (chemical property) of metabolic enzymes (biological functions). Metabolite, lipid, and FFA & acyls layers are assigned based on the chemical properties of the molecules. Edges are the regulatory connections between nodes, connected based on the regulatory relationships among omic layers (fig. S3**A**, **B**, **C**, and **D**), using bioinformatic platforms. For example, edges from metabolite, lipid, and FFA & acyls layers to metabolic reaction layer represent the regulatory relationships among metabolites, lipids, and FFA & acyls— as substrates, products, and allosteric regulators— to metabolic reactions. Edges from enzyme_protein and enzyme_PTM to the metabolic reaction layer represent the regulatory relationships between metabolic enzymes and metabolic reactions mediated by enzyme abundance (enzyme_protein) and phosphorylation (enzyme_PTM). Colors of nodes and edges represent the type of response: green, common in both WT and *ob*/*ob* mice; blue, specific to WT mice; red, specific to *ob*/*ob* mice; magenta, opposite responses between WT and *ob*/*ob* mice (Fig. 3**A** and **B**).

**Fig. 3:**
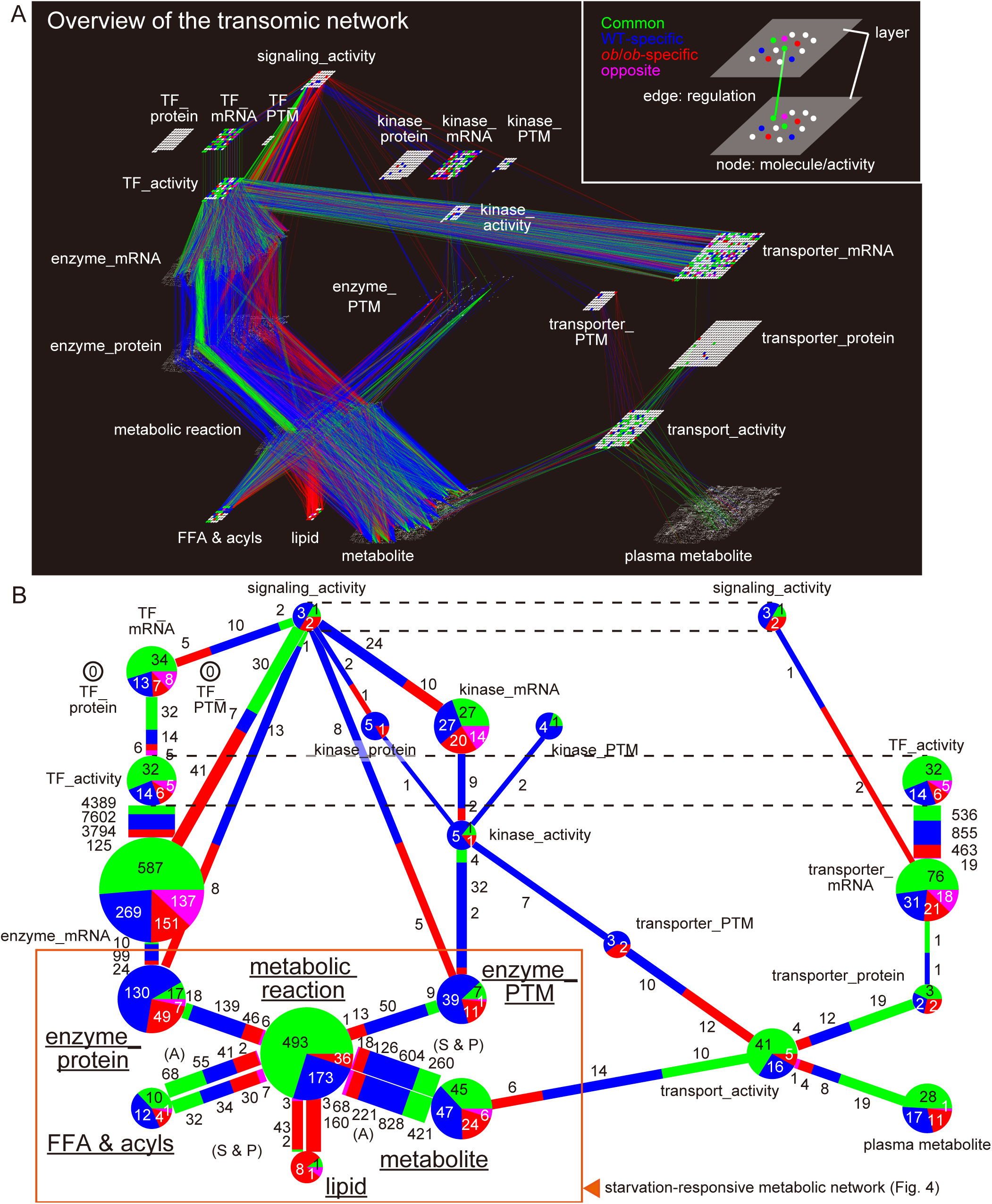
Global starvation-responsive transomic network (**A**) An overview of the global starvation-responsive transomic network. Nodes in the layers represent metabolic reactions, transport activities, measured molecules, measured phosphorylation events, and the inferred activities of TF, protein kinase, and signaling pathway. Edges between the nodes represent regulatory relationships identified using bioinformatic platforms. Colors represent the responsiveness: green indicates common responses both in WT and *ob*/*ob* mice; blue, specific to WT mice; red, specific to *ob*/*ob* mice; magenta, opposite responses between the WT and *ob*/*ob* mice. For metabolic reaction and transport_activity layers, the classes were defined by the existences of regulations regardless of the regulatory directions (activation or inhibition): green indicates the node is regulated in both WT and *ob*/*ob* mice; blue, specific to WT mice; red, specific to *ob*/*ob* mice. (**B**) The proportions of nodes and edges classified as common (green), WT-specific (blue), *ob*/*ob*-specific (red), and opposite (magenta) in the global starvation-responsive transomic network. Pie charts represent the distribution of nodes in the omic layers, and bar plots between pie charts represent the distribution of edges between omic layers. Type of regulation into the metabolic reactions for metabolite, FFA& acyls, and lipid are indicated as allosteric regulation (denoted as “A”) and substrate and product (denoted as “S & P”). The starvation-responsive metabolic network is outlined in orange, and layer names included in the starvation-responsive metabolic networks are underlined.

**Table 8:** Omic layers in the global starvation-responsive metabolic network Classification of molecules or inferred regulators into omic layers, biological function, or chemical property.

| Layer name | Chemical property | Biological function | Omic |
| --- | --- | --- | --- |
| metabolite | metabolite | metabolite | metabolome |
| metabolic reaction | - | metabolic reaction | inference |
| enzyme_mRNA | mRNA | enzyme | transcriptome |
| enzyme_protein | protein | enzyme | proteome |
| enzyme_PTMs | PTM | enzyme | phosphoproteome |
| TF_mRNA | mRNA | TF | transcriptome |
| TF_protein | protein | TF | proteome |
| TF_PTMs | PTM | TF | phosphoproteome |
| TF_activity | - | TF | inference |
| kinase_mRNA | mRNA | kinase | transcriptome |
| kinase_protein | protein | kinase | proteome |
| kinase_PTMs | PTM | kinase | phosphoproteome |
| kinase_activity | - | kinase | inference |
| signaling_activity | - | signaling pathway | inference |
| transporter_mRNA | mRNA | transporter | transcriptome |
| transporter_protein | protein | transporter | proteome |
| transporter_PTMs | PTM | transporter | phosphoproteome |
| transporter_activity | - | transporter | inference |
| lipid | lipid | lipid | lipidome |
| bloodmetabolite | metabolite | metabolite in blood | metabolome |
| FFA & acyls | fatty acid, acyl-CoA, acyl-carnitine | FFA & Acyls | fatty acid, acyl-CoA, acyl-carnitine |

We showed the proportions of the commonly, WT-specifically, *ob*/*ob*-specifically, and oppositely responsive nodes and edges in the global starvation-responsive transomic network (Fig 3**B**). In this network, pie charts represent the proportions of the classes of responses of nodes in the omic layers, and bar plots between pie charts represent proportions of the classes of edges between omic layers. For clear visualization, pie charts for signaling_activity and TF_activity are displayed twice.

We assessed the proportions of the global starvation-responsive transomic network (Fig. 3**B**). Although less than half (47 out of 122 nodes) of the starvation-responsive metabolites (“metabolite” in Fig. 3**B**) were specific to WT mice, more than half regulatory connections between the metabolites (“metabolite” in Fig. 3**B**) and the metabolic reactions (“metabolic reaction” in Fig. 3**B**) were specific to WT mice (604 out of 1,008 substrate and product relationships and 828 out of 1,538 allosteric regulatory relationships). Among the starvation-responsive lipids (“lipid” in Fig. 3**B**), but not the FFA & acyls (“FFA & acyls” in Fig. 3**B**), most changes (8 out of 10) were specific to *ob*/*ob* mice. This enrichment in the *ob*/*ob*-specific response also occurred among the substrate, product, and allosteric regulatory connections between the lipids (“lipid” in Fig. 3**B**) and the metabolic reactions (“metabolic reaction” in Fig. 3**B**) with 43 out of 48 substrate and product relationships *ob/ob*-specific and 160 out of 163 allosteric regulator relationships *ob/ob*-specific. This enrichment of *ob*/*ob*-specifically responsive regulatory connections was due to a larger number of *ob*/*ob*-specifically responsive lipids. In addition, the number of regulations from commonly responsive lipids and oppositely responsive lipids was very small (fig. S3**E**).

In the enzyme_protein layer (“enzyme_protein” in Fig. 3**B**), most nodes were in the WT-specific class (130 of the 203, 64%) and only 17 of the 203 (8.4%) starvation-responsive nodes responded in common between WT and *ob/ob* mice.

However, more than half of all the starvation-responsive nodes (587 out of 1144; 51.3%) exhibited a common response in the enzyme_mRNA layer (“enzyme_mRNA” in Fig. 3**B**). This disconnection between the responses at the mRNA and protein levels indicated that the protein abundance of the metabolic enzymes is determined processes occurring after transcription, such as the rate of translation and protein degradation, that are known to be downregulated in *ob*/*ob* mice (*59*, *60*). Among most of the nodes and edges related to phosphorylation (“kinase_protein”, “kinase_PTM”, “kinase_activity”, and “enzyme_PTM” in Fig. 3**B**), WT-specific groups occurred most often compared to the common, *ob*/*ob*-specific, and opposite groups. These results suggested a decrease in regulation by phosphorylation in response to starvation in *ob*/*ob* mice.

Although regulatory connections from metabolites, lipids, enzyme_protein, and enzyme_PTM contain a small portion of commonly responsive regulatory connections, most of the starvation-responsive metabolic reactions (“metabolic reaction” in Fig. 3**B**) were commonly responsive in both WT and to *ob*/*ob* mice (493 out of 702).

Compared with nodes in the other classes of responses, nodes with common responses were most abundant in several layers related to transport_activities (“plasma metabolite”, “transporter_mRNA”, “transporter_protein”, and “transport_activity” in Fig. 3**B**).

Taken together, the regulatory connections from metabolite, lipid, enzyme_protein, and enzyme_PTM layers to metabolic reactions were different between WT and *ob/ob* mice. Disconnections between the responses of enzyme_mRNA and enzyme_protein were observed in *ob/ob* mice, probably due to the disrupted protein translation and degradation, while the transport activities were maintained.

### Structural properties of the starvation-responsive metabolic network

We focused on the sub-network of the global starvation-responsive transomic network that directly regulate metabolic reactions, defined as the starvation-responsive metabolic network (Fig. 4). To visualize the differences between WT and *ob/ob* mice, we show the starvation-responsive metabolic network separately for WT (Fig. 4**A**) and *ob*/*ob* (Fig. 4**B**) mice. For each genotype, the networks include the layers for metabolite, lipid, FFA & acyls, enzyme_protein, and enzyme_PTM (Data File S6) and their regulatory relationship to the metabolic reaction layer (fig. S3**A**). The edges connecting metabolite, lipid, and FFA & acyls layers to the metabolic reaction layer represent their roles as substrates, products, and allosteric regulators of the metabolic reactions.

**Fig. 4:**
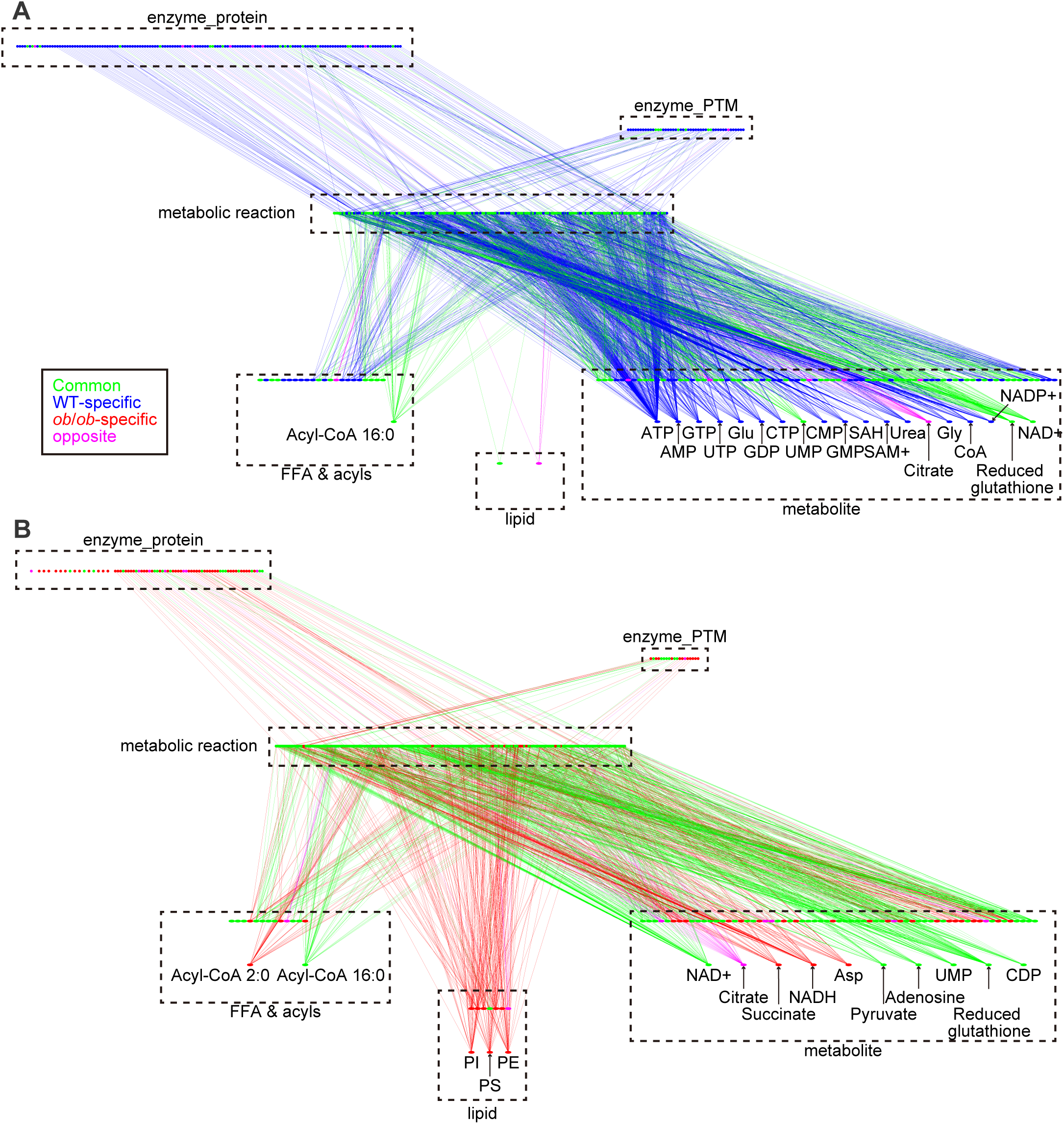
Starvation-responsive metabolic network Projected views of the starvation-responsive metabolic networks in WT (**A**) and *ob*/*ob* mice (**B**). The network is composed of direct regulators of metabolic reactions: metabolites, lipids, FFA & acyls, enzyme_protein, and enzyme_PTM. Colors represent the responsiveness: green indicates common responses both in WT and *ob*/*ob* mice; blue, specific to WT mice; red, specific to *ob*/*ob* mice; magenta, opposite responses between the two genotypes. Hub nodes, which will be defined in Fig. 5, are labeled.

We analyzed the structural properties of the starvation-responsive metabolic networks in WT and *ob/ob* mice (Fig. 5). We calculated the number of edges connected to each node (called “degree”) and then assessed the degree distribution of the network, by a log-log scatter plot (Fig. 5**A**). We found that nodes with large degree were rare, while nodes with small degree were abundant. This type of network is called a scale-free network, and nodes with large degree are called “hub”, while nodes with small degree are called “leaf” (Fig. 5**B**). Degree distribution of a scale-free network can be approximated by a power-law (*32*, *43*, *61*) defined as:

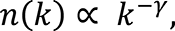

where *n*(*k*) denotes the number of nodes with a degree of *k*, and γ is a constant called the scaling parameter. The scaling parameter represents the heterogeneity of the degree distribution (*62*) with a small scaling parameter representing a large heterogeneity of degree distribution of the network. From the calculated scaling parameter for each genotype’s network, we found that the starvation-responsive network for WT and *ob/ob* mice had a similar amount of heterogeneity in the degree distribution (Fig. 5**A**). Because the calculated scaling parameters do not meet the strict definition of scale-free network that should be 2 < γ < 3 (*61*, *63*), we call the starvation-responsive metabolic network as a scale-free-like network.

**Fig. 5:**
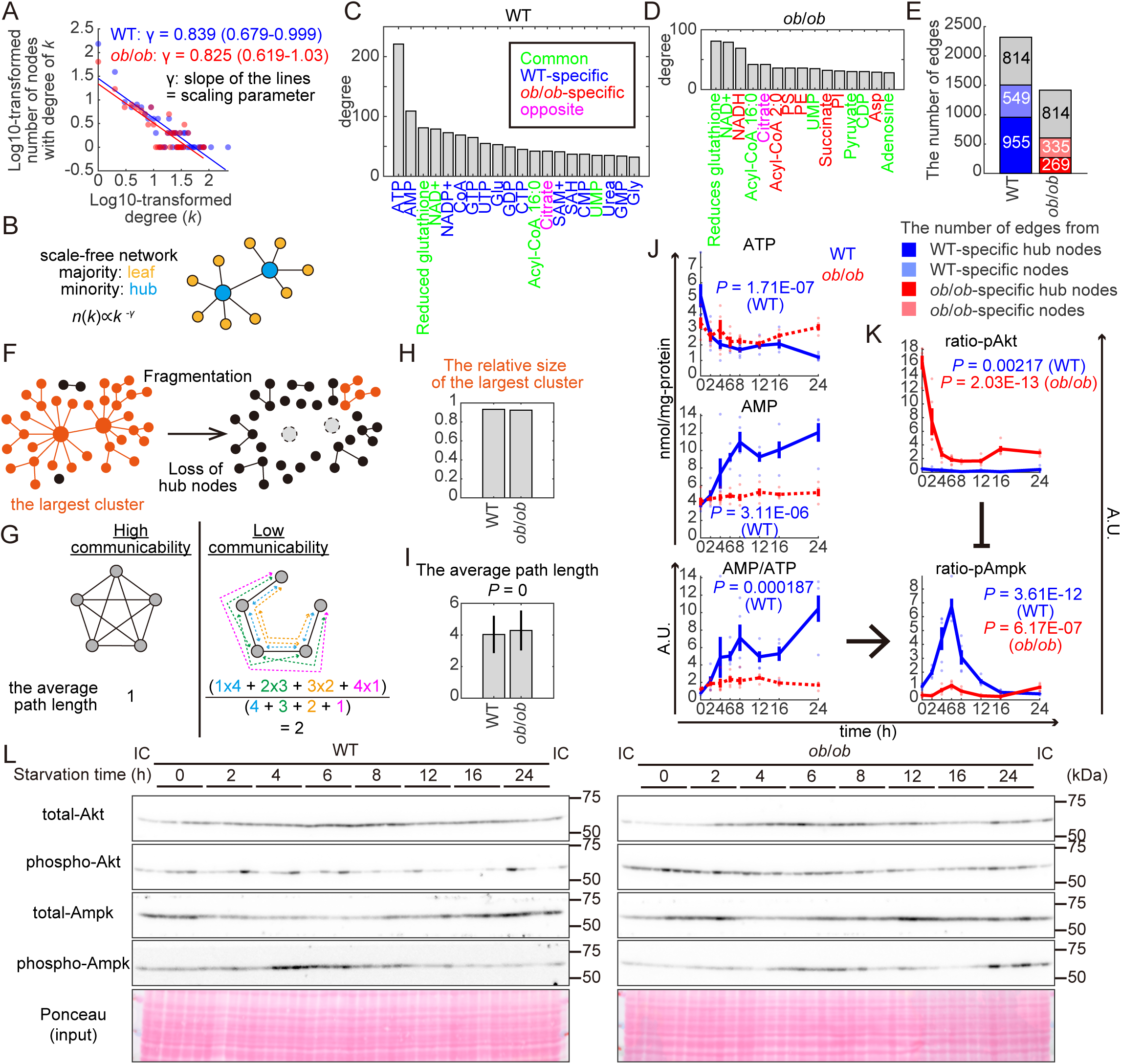
Structural properties of the starvation-responsive metabolic network (**A**) Degree distributions of the starvation-responsive metabolic network in WT (blue) and *ob*/*ob* (red) mice. The x-axis represents the log10-transformed degree (*k*); the y-axis represents the log10-transformed number of nodes with degree of *k*. Lines show the linear regression of the degree distributions. The slope of the lines (γ), which is defined as a scaling parameter, and the 95% credible interval are shown. *n* = 990 in WT and *n* = 721 in *ob*/*ob* mice for the number of nodes. (**B**) Diagram of a scale-free network, where *n*(*k*) denotes the number of nodes with degree of *k*, and γ is a constant called a scaling parameter. Circles represent nodes; lines represent edges. Light blue represents the hub nodes; yellow represents leaf nodes. (**C** and **D**) Hub nodes with the top 2% highest degree in WT (**C**) and *ob*/*ob* mice (**D**). The x-axis represents hub nodes. The y-axis represents degrees. Colors represent the responsiveness: green indicates common responses both in WT and *ob*/*ob* mice; blue, specific to WT mice; red, specific to *ob*/*ob* mice; magenta, opposite responses between WT and *ob*/*ob* mice. (**E**) The number of edges (y-axis) from WT-specifically responsive hub nodes (blue), WT-specific nodes except for the hub nodes (light blue), *ob*/*ob*-specific hub nodes (red), *ob*/*ob*-specific nodes except for the hub nodes (light red), other nodes (commonly or oppositely responsive, gray). (**F**) Illustration of fragmentation of a network through loss of hub nodes. The largest cluster is orange. (**G**) Illustration of how communicability of a network influences average path length (average length of the shortest paths between all the pairs of nodes). (**H**) The relative size of the largest cluster in the starvation-responsive metabolic network in WT and *ob*/*ob* mice, calculated as the number of nodes in the largest cluster of the network divided by the total number of responsive nodes in the network for each genotype. (**I**) The average path length of the starvation-responsive metabolic network in WT and *ob*/*ob* mice. *n* = 853,884 paths in WT and *n* = 443,643 paths in *ob*/*ob* mice. The error bars show standard deviation. *P* value of *t*-test is shown. (**J**) Time courses of ATP, AMP, and AMP/ATP ratio during starvation in WT and *ob*/*ob* mice. The x-axis represents the starvation time (hours) and the y-axis represents the concentration (nmol/mg-protein) for ATP and AMP, and ratio for AMP/ATP. Blue lines are the responses to starvation in WT mice, and red lines are those in *ob*/*ob* mice. Solid lines represent that the responses were significant; dotted lines represent no significance. Data are shown as the mean and SEM. Dots represent the data from individual mice. (**K**) Time courses of phosphorylation ratio of Akt and Ampk during starvation in WT and *ob*/*ob* mice. The x-axis represents the starvation time (hours) and the y-axis represents the phosphorylation ratio (the amount of phosphorylated protein divided by the total amount of protein) of Akt and Ampk. (**L**) Images of the western blotting analysis for (**K**) at the indicated starvation time (h) of the indicated genotype. IC represents internal control to normalize the signals across membranes. *n* = 5 biological replicates per group. *P* values of one-way ANOVA of the molecules or ratio with significant changes are shown.

We defined the nodes that had a degree in the top 2% among the responsive nodes in each genotype as hub nodes (Fig. 5**C** and **D**), that regulated multiple metabolic pathways (fig. S4). We identified 20 hub nodes in the network of WT mice and 15 hub nodes in the network of *ob*/*ob* mice (Fig. 5**C**, **D** and labeled on Fig. 4**A**, **B**). In WT mice, 15 of the 20 were WT-specific (Fig. 4**A**; Fig. 5**C**). In *ob*/*ob* mice, 7 hub nodes out of 15 were *ob*/*ob*-specific and 3 of the 7 were lipids (Fig. 4**B**; Fig. 5**D**). The hub nodes with the highest degrees were ATP (degree >200) and AMP (degree >100) in the network of WT mice (Fig. 5**C**); whereas all hub nodes in the network of *ob*/*ob* mice had degrees less than 100 with reduced glutathione (degree 81) and NADH (degree 69) (Fig. 5**D**). Thus, the starvation-responsive metabolic network in *ob/ob* mice lacked the most highly connected hub nodes such as ATP and AMP. The number of edges from the 15 WT-specific hub nodes was 955 out of 2,318 edges (41%) in the starvation-responsive metabolic network in WT mice (Fig. 5**E**), suggesting a large contribution of WT-specific hub nodes to the starvation-responsive metabolic network of WT mice, which was lost in *ob*/*ob* mice.

To evaluate the effect of the loss of responsiveness of the highly connected hub nodes in *ob/ob* mice, we analyzed the structural properties of the starvation-responsive metabolic network: the relative size of the largest cluster (Fig. 5**F**) and average path length (Fig. 5**G**). Cluster is a set of nodes connected through edges regardless of the distances. The largest cluster is the cluster that includes the largest number of nodes in a network. Fragmentation breaks the network into isolated clusters and reduce the relative size of the largest cluster (Fig. 5**F**), that is the number of nodes in the largest cluster divided by the total number of the responsive nodes in each genotype. Average path length (Fig. 5**G**) is the average number of edges connecting two edges, which increases by loss of communicability among nodes (*64*). In the starvation-responsive metabolic networks of WT and *ob*/*ob* mice, the relative sizes of the largest cluster were similar (Fig. 5**H**), indicating that the network was not fragmentated in *ob*/*ob* mice. Additionally, the average path lengths were significantly different but almost similar in the networks of the WT and *ob*/*ob* mice (Fig. 5**I**), indicating that communicability was almost maintained in *ob*/*ob* mice. Collectively, all structural properties of the starvation-responsive metabolic networks, the scaling parameter (Fig. 5**A**), the relative size of the largest cluster (Fig. 5**H**), and the average path length (Fig. 5**I**) were similar between WT and *ob*/*ob* mice. These results indicated that the starvation-responsive metabolic network was structurally maintained in *ob*/*ob* mice, despite the loss of responsiveness of highly connected hub nodes such as ATP and AMP.

The maintenance of the structural properties of the starvation-responsive metabolic network in *ob*/*ob* mice seems counter-intuitive because it has been known that the loss of hub nodes in a scale-free network drastically alter the structural properties (*64*). Taking a close look at the degree distribution (Fig. 5**A** and fig. S5), the number of nodes (y-axis of Fig. 5**A**) with small degree (log10-transformed degree < 0.5 in the x-axis of Fig. 5**A**) were smaller in *ob*/*ob* mice than in WT mice, but the number of nodes (y-axis of Fig. 5**A**) with log10-transformed degree ranging from 0.5 to 2 (x-axis of Fig. 5**A**) were comparable between WT and *ob*/*ob* mice (Fig. 5**A**, see fig. S5 in detail). Specifically, the percentage of nodes among the responsive nodes was higher in *ob*/*ob* mice than WT mice for nodes with a log10-transformed degree between 0.5 and 1 (fig. S5**G**) and between 1 and 1.5 (fig. S5**H**). Totally, the percentages of nodes with relatively large degrees (log10-transformed degree > 1) were similar between WT and *ob*/*ob* mice (fig. S5**K**). The maintenance of nodes with relatively large degrees could maintain the structural properties of the starvation-responsive metabolic network in *ob*/*ob* mice.

To experimentally evaluate the effects of the loss of the responsiveness of ATP and AMP in *ob*/*ob* mice, we measured the phosphorylation of Ampk, which reflect its kinase activity, by western blotting. Ampk activity is known to depend on the AMP/ATP ratio (*65*), that was WT-specifically responsive in our dataset (Fig. 5**J**). Starvation-responsive phosphorylation of Ampk in *ob*/*ob* mice was weaker than in WT mice (Fig. 5**K**, and **L**), although significant responses were observed in both WT and *ob*/*ob* mice. This weak response of Ampk in *ob*/*ob* mice could be due to an aberrantly large decrease of phosphorylation ratio of Akt (Fig. 5**K**, and **L**), a negative regulator of Ampk (*66*). Taken together, the loss of AMP/ATP responsiveness reduced the response of Ampk activity, although it was partially rescued by the larger of Akt activity.

### Temporal order of the starvation-responsive metabolic network in WT mice and its disruption in *ob*/*ob* mice

Adaptation to starvation is a temporally well-ordered metabolic process (*1–6*). To explore the relationships between the network structure and the temporal order of the starvation-induced changes in the starvation-responsive metabolic network, we defined the time when the response reached half of the maximum amplitude of the responses as *t*-half (*37*). A small *t*-half represents rapid responsiveness, and a large *t*-half represents slow responsiveness. We analyzed molecules [metabolites, lipids, FFA & acyls, proteins (enzyme_protein), and phosphorylation events (enzyme_PTM)] in the starvation-responsive metabolic networks, because *t*-half can be defined only for measured molecules not for metabolic reactions. We refer to hub nodes (Fig. 5**C** and **D**) as hub molecules because all the hub nodes were molecules.

We investigated the relationships between network structure and *t*-half in WT and *ob*/*ob* mice. We found a negative correlation between *t*-half and degree of molecules in WT mice (Fig. 6**A**), indicating that the molecules connected with many edges responded rapidly to starvation. In addition, 9 out of 20 hub molecules, including ATP, were essential for the negative correlation with *t*-half in WT mice (fig. S6), suggesting a dominant role of hub molecules in coordinating the temporal order of the starvation-responsive metabolic network. By contrast, no significant correlation between *t*-half and degree of molecules was observed in *ob*/*ob* mice (Fig. 6**B**). Indeed, molecules with many edges, either increased or decreased, responded to starvation more rapidly than those with few edges in WT mice (Fig. 6**A**), but not in *ob*/*ob* mice (Fig. 6**B**). The *t*-half values of *ob*/*ob*-specific hub molecules were larger than those of WT-specific hub molecules (Fig. 6**C**), indicating a slower response of the *ob*/*ob*-specific hub molecules.

**Fig. 6:**
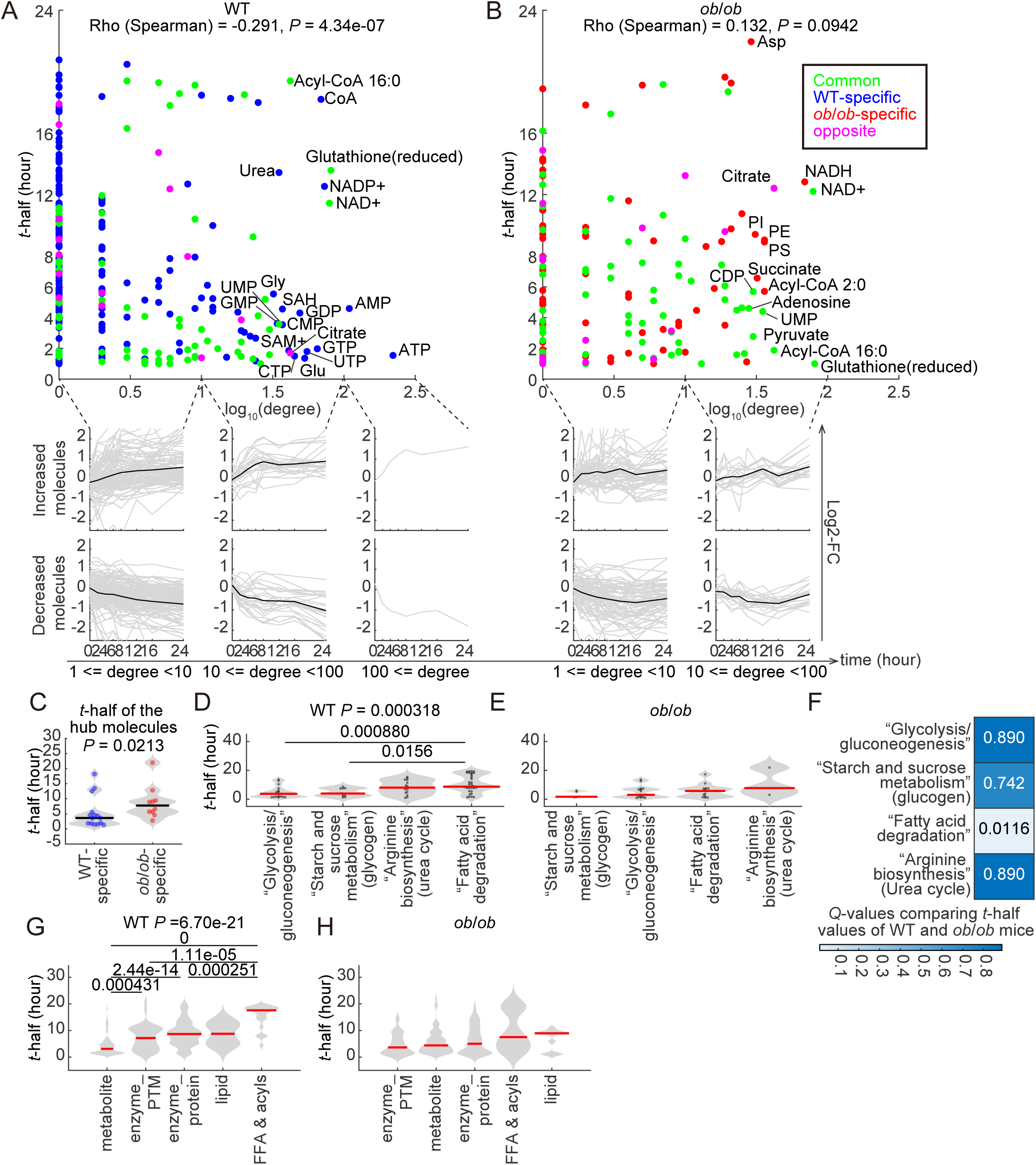
Temporal order of the starvation-responsive metabolic network in WT mice and its disruption in *ob*/*ob* mice (**A** and **B**) Scatter plots for the log10-transformed degree (x-axis) and *t*-half (y-axis) in WT (**A**) and *ob*/*ob* (**B**) mice. Dots represent the molecules (including phosphorylation events) in the starvation-responsive metabolic network and colors denote the responsiveness: green indicates common responses both in WT and *ob*/*ob* mice; blue, specific to WT mice; red, specific to *ob*/*ob* mice; magenta, opposite responses between WT and *ob*/*ob* mice. Spearman’s correlation coefficient (Rho) and *P* values are shown. Hub molecules for each genotype are labeled (see Fig. 5C and **D**). Time courses of increased and decreased molecules connected with few (1 ≤ degree < 10), relatively many (10 ≤ degree < 100), and many (100 ≤ degree) edges in WT (**A**) and *ob*/*ob* (**B**) mice are also shown. The x-axis represents starvation time (hours) and the y-axis represents the log2-transformed concentration to the concentrations before starvation of the molecules in each genotype. *n* = 291 molecules for WT and *n* = 163 molecules for *ob*/*ob* mice. (**C**) Values of *t*-half for the WT-specific and *ob*/*ob*-specific hub molecules (see Fig. 5C and **D**). The *P* value of Wilcoxon’s rank sum test is shown. *n* = 15 hub molecules for WT and *n* = 10 hub molecules for *ob*/*ob* mice. (**D** and **E**) Distributions of *t*-half values of molecules (metabolites, lipids, FFA & acyls, proteins, and phosphorylation events) in the indicated metabolic pathways in WT (**D**) and *ob*/*ob* mice (**E**). The data are ordered along the x-axis by the median *t*-half (red line) with each molecule represented by a dot. *n* = 19, 10, 14, and 36 molecules in mice WT for “Glycolysis/gluconeogenesis”, “Starch and sucrose metabolism”, “Arginine biosynthesis”, and “Fatty acid degradation”. *n* = 5, 15, 13, and 3 molecules in *ob*/*ob* mice for “Starch and sucrose metabolism”, “Glycolysis/gluconeogenesis”, “Fatty acid degradation”, and “Arginine biosynthesis”. *P* values of the Kruskal-Wallis test are shown above the panels and those of the Dunn-Sidak test are shown above the lines indicating the pairs of metabolic pathways with significant differences (*P* < 0.05). (**F**) *Q*-values comparing *t*-half values of WT and *ob*/*ob* mice for each metabolic pathway. (**G** and **H**) Distributions of *t*-half values in each omic layer in the starvation-responsive metabolic network in WT (**G**) and in *ob*/*ob* (**H**) mice. The data are ordered along the x-axis by the median *t*-half (red line). *n* = 98, 47, 154, 2, and 23 molecules in mice WT for metabolite, enzyme_PTM, enzyme_protein, lipid, and FFA & acyls. *n* = 19, 75, 73, 15, and 10 molecules in *ob*/*ob* mice for enzyme_PTM, metabolite, enzyme_protein, FFA & acyls, and lipid. *P* values of the Kruskal-Wallis test are shown above the panels and those of the Dunn-Sidak test are shown above the lines indicating the pairs of omic layers with significant differences (*P* < 0.05).

We conducted *t*-half analysis to evaluate how the temporal order of use of the metabolic fuels such as glycogen, carbohydrates, amino acids and lipids (*1–6*), differs during starvation in WT and *ob/ob* mice. We classified molecules into the following metabolic pathways: “Starch and sucrose metabolism” (containing glycogen and glycogen metabolic enzymes), “Glycolysis/gluconeogenesis”, “Arginine biosynthesis” (containing the urea cycle that could indicate amino acid degradation), and “Fatty acid degradation” of the KEGG database (*55–57*). We compared *t*-half values for the molecules in each of the metabolic pathways for the starvation-responsive metabolic network in WT (Fig. 6**D**) and *ob*/*ob* mice (Fig. 6**E**). In WT mice, we observed significant differences in *t*-half values of molecules among the investigated metabolic pathways (Fig. 6**D**), indicating that the metabolic processes are temporally ordered across the metabolic pathways in WT mice. Among them, “Fatty acid degradation” responded to starvation the slowest, which is consistent with fatty acids as fuel for the late stage of starvation (*1–6*). By contrast, no significant differences in *t*-half values were observed in *ob*/*ob* mice (Fig. 6**E**), indicating that the temporal order among the metabolic pathways was disrupted in *ob*/*ob* mice.

To examine which metabolic pathways largely contribute to the disturbed temporal order among metabolic pathways in *ob*/*ob* mice, we compared the *t*-half values between WT and *ob*/*ob* mice for each metabolic pathway (Fig. 6**F**).

We found that *t*-half values were significantly smaller in “Fatty acid degradation” of *ob*/*ob* mice than WT mice. “Fatty acid degradation” degrades fatty acids to ketone body (3-Hydroxybutyrate) through acyl-CoAs and acyl-carnitines during starvation (fig. S7**A**).We manually reconstructed major metabolic reactions for fatty acid degradation based on “Fatty acid degradation” (mmu00071), “Propanoate metabolism” (mmu00640), and “Butanoate metabolism” (mmu00650) in KEGG database (*55–57*), and found that many metabolic enzymes (enzyme_protein and enzyme_PTM) and allosteric regulators regulate the fatty acid degradation (fig. S7**B**). Among substrates (fatty acids), intermediates (acyl-CoAs and acyl-carnitines), products (3-Hydroxybutyrate, a ketone body), and regulators (metabolites, enzyme_protein, and enzyme_PTM), only the *t*-half values of acyl-CoAs showed statistically significant differences between WT and *ob*/*ob* mice (fig. S7**C**), suggesting that the significant difference in *t*-half values between WT and *ob*/*ob* mice (Fig. 6**F**) was mainly due to acyl-CoAs. Acyl-CoAs increased smoothly and continuously in WT mice, but showed a transient increase in the early phase of starvation in *ob*/*ob* mice (figs. S7**A** and **B**).

Although the difference in *t*-half values between WT and *ob*/*ob* mice was not statistically significant, acyl-carnitines showed time courses similar to those of acyl-CoAs (figs. S7**A** and **B**). By contrast, fatty acids, the source of acyl-CoAs and acyl-carnitines, showed similar time courses between WT and *ob*/*ob* mice. This disconnection in the time courses between substrates (fatty acids) and intermediate (acyl-CoAs and acyl-carnitines) in fatty acid degradation could be a hallmark of disrupted flux in fatty acid degradation.

This disruption could be due to the enzyme_protein because the number of responsive enzyme_protein is the largest among substrates, intermediates, products, and regulators in WT mice, and this number largely decreased in *ob*/*ob* mice (fig. S7**D**).

However, most enzyme_mRNA corresponding to these enzyme_protein were commonly responsive in both WT and *ob*/*ob* mice (11 out of 13, fig. S7**B**). This disconnection between responses at the mRNA and protein levels could be due to the decreased rate of protein translation and degradation in *ob*/*ob* mice (Fig. 3**B**) (*59*, *60*). Consistently, the correlations between the time courses of enzyme_mRNA and enzyme_protein for each metabolic enzyme in fatty acid degradation were significantly decreased in *ob*/*ob* mice (fig. S7**E**). To further validate this hypothesis, we evaluated the protein phosphorylation related to protein translational (fig. S8**A** and **B**) and autophagic (fig. S8**C** and **D**) activities. Phosphorylation of eIF2α represents inhibition of translation (*67*). Phosphorylation ratio of eIF2α increased in *ob*/*ob* mice before starvation (fig. S8**A** and **B**), suggesting translational deficiency in *ob*/*ob* mice, consistent with the previous report (*60*). Although measuring autophagic activity in vivo without an exogenous probe remains challenging (*68*), phosphorylation of p62 at Ser^351^ is an autophagic substrate (*69*) and can serve as a marker for autophagic degradation. Phosphorylated p62 significantly decreased in WT mice during starvation, but not in *ob*/*ob* mice (fig. S8**C** and **D**), suggesting autophagy deficiency in *ob*/*ob* mice, consistent with the previous report (*59*). Taken together, deficiency in protein translation and degradation in *ob*/*ob* mice (fig. S8) reduced the number of responsive enzyme_protein in *ob*/*ob* mice (fig. S7) and disrupted the flux of fatty acid degradation, reslting in a transient increase in acyl-CoAs and acyl-carnitines during starvation in *ob*/*ob* mice (figs. S7**A** and **B**).

In addition, we compared *t*-half values of molecules in each omic layer in the starvation-responsive metabolic network in WT (Fig. 6**G**) and *ob*/*ob* mice (Fig. 6**H**). In WT mice, we observed significant differences in *t*-half values of molecules among the omics layers (Fig. 6**G**), indicating that adaptation to starvation is temporally ordered across omic layers in WT mice. Metabolites responded to starvation fastest, indicating that metabolic reactions are initially regulated by metabolites through the effect of starvation-induced changes in substrates and products and metabolite-mediated allosteric regulators. Phosphorylation (enzyme_PTM), protein abundance (enzyme_protein), and lipids responded next. The FFA & acyls layer responded the slowest, consistent with fatty acid degradation important for the late stages of adaptation to starvation (*3*). By contrast, no significant differences in *t*-half values were observed in *ob*/*ob* mice (Fig. 6**H**), indicating that the temporal order of the starvation response by the omic layers was disrupted.

Taken together, these results indicated that hub molecules coordinate the temporal order of the responses of molecules, metabolic pathways, and omic layers in the starvation-responsive metabolic network in WT mice. In *ob*/*ob* mice, the temporal order of the starvation response throughout the network was disrupted. Deficiency in protein translation and degradation in *ob*/*ob* mice (Fig. 3**B**, and fig. S7**E** and S8) (*59*, *60*) also disturbed the fatty acid degradation (fig. S7).

### Co-regulation among molecules in the starvation-responsive metabolic network and disruption in *ob*/*ob* mice

Molecules positively or negatively regulated by common regulatory mechanisms show positively or negatively correlated time courses, defined as co-regulation (Fig. 7**A**). Thus, positively or negatively correlated time courses represent co-regulations, which could be disrupted by the disrupted temporal order of metabolites, lipids, FFA & acyls, protein abundance, and phosphorylation events in *ob*/*ob* mice (Fig. 6). As an index of temporal co-regulation (Fig. 7**A**), we calculated the pairwise Pearson’s correlation coefficients for time courses of each pair of molecules (including phosphorylation events) in the starvation-responsive metabolic network in WT and *ob*/*ob* mice, respectively (Fig. 7**B**). The distribution of correlation coefficients in WT mice was bimodal with peaks at around 0.75 and −0.75, indicating both positive or negative co-regulation occurred among molecules. By contrast, the distribution of correlation coefficients was unimodal with a broad peak around 0 in *ob*/*ob* mice, indicating that co-regulation was disrupted in *ob*/*ob* mice.

**Fig. 7:**
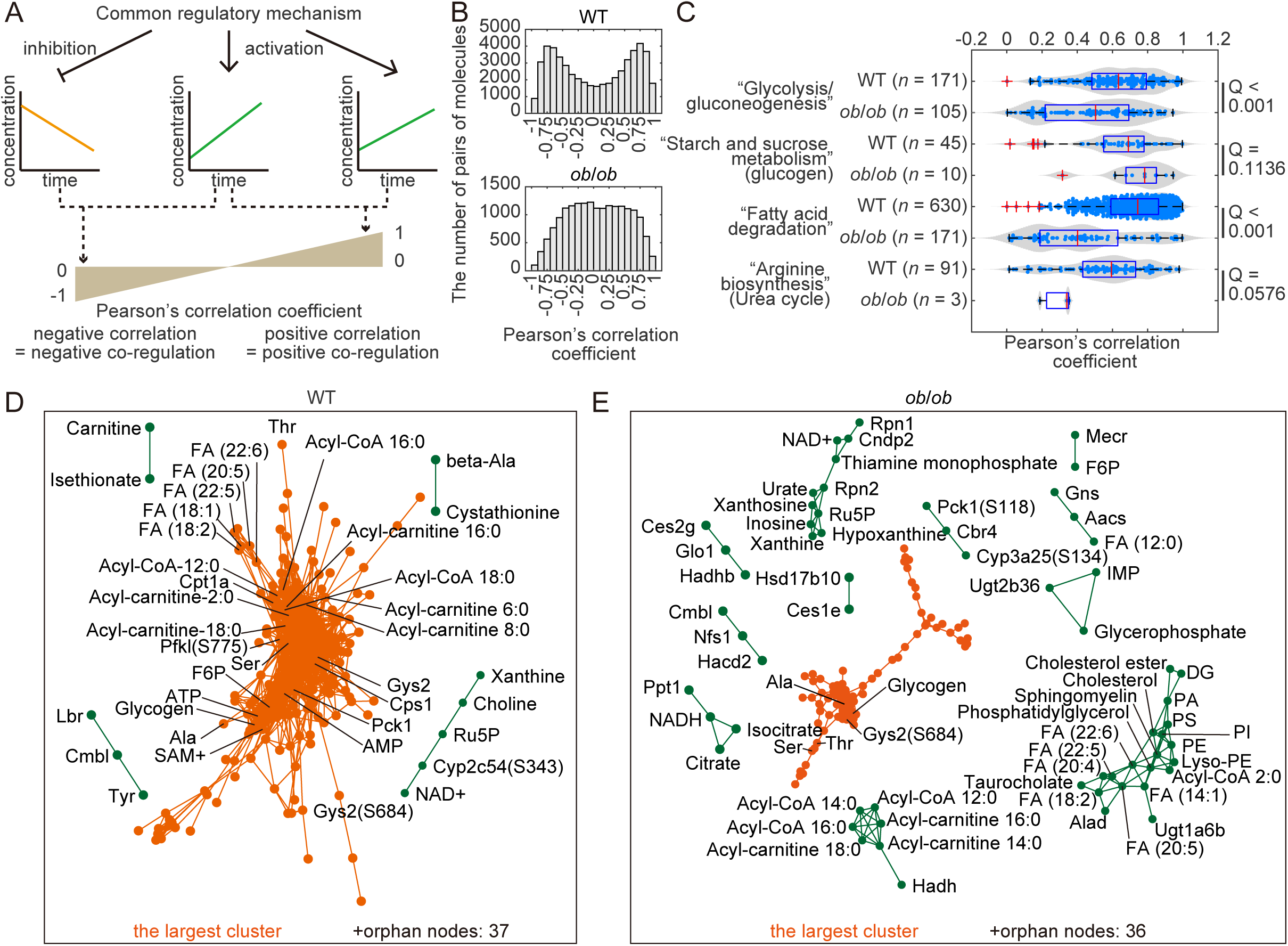
Co-regulation among molecules in the starvation-responsive metabolic network and disruption in *ob*/*ob* mice (**A**) Schematic representation of the analyses for co-regulation among the molecules in the starvation-responsive metabolic network. Time courses of molecules sharing a common regulatory mechanism could be positively or negatively correlated, defined as co-regulation. Pairwise Pearson’s correlation coefficients were calculated for the time courses of molecules that showed significant responses during starvation in the starvation-responsive metabolic network in WT and *ob*/*ob* mice. Positive correlation represents positive co-regulation and negative correlation represents negative co-regulation. (**B**) Distribution of the pairwise Pearson’s correlation coefficients for time courses of responsive molecules in the starvation-responsive metabolic network in WT and *ob*/*ob* mice. The x-axis represents correlation coefficients; the y-axis represents the number of pairs of molecules examined for correlation. Pairs with an absolute value of correlation coefficient > 0.9 were connected by edges in the correlation metabolic network (see fig. S12). *n* = 324 molecules for WT mice, *n* = 192 molecules for *ob*/*ob* mice. (**C**) The absolute values of pairwise Pearson’s correlation coefficients among time courses of responsive molecules in the indicated metabolic pathways in WT and *ob*/*ob* mice. The x-axis represents the absolute values of Pearson’s correlation coefficient. The y-axis represents metabolic pathways and genotypes. Light blue dots represent the absolute values of pairwise Pearson’s correlation coefficients among time courses of responsive molecules in each metabolic pathway of the starvation-responsive metabolic network. Red vertical lines indicate median, boxes the interquartile range (25th to 75th percentile), and whiskers 1.5 times the interquartile range. Red crosses represent outliers. *Q*-values comparing the correlation coefficients of WT and *ob*/*ob* mice by Wilcoxon rank sum test followed by FDR through Benjamini-Hochberg procedure are shown. The numbers of pairs of molecules are shown in the parenthesis. (**D** and **E**) The correlation metabolic network during starvation in WT (**D**) and *ob*/*ob* mice (**E**). Orange indicates the largest cluster; all other clusters are dark green. Individual orphan nodes with no connections are shown only by numbers.

We evaluated the extent of disruption of co-regulations among molecules in *ob*/*ob* mice in each metabolic pathway (Fig. 7**C**). We compared the absolute values of pairwise Pearson’s correlation coefficients for the time courses of each pair of molecules (including phosphorylation events) in each metabolic pathway in the starvation-responsive metabolic network between WT and *ob*/*ob* mice (Fig. 7**C**). We found statistically significant (*Q* < 0.1) decrease in the absolute values of pairwise Pearson’s correlation coefficients in *ob*/*ob* mice compared to WT mice, in all the investigated metabolic pathways except for “Starch and sucrose metabolism”. This suggests that the extent of disruption in co-regulations in *ob*/*ob* mice was so substantial, leading to disrupted co-regulation even among the molecules in the same metabolic pathways.

To examine the detailed disruptions in each metabolic pathway, we manually reconstructed major metabolic reactions for glycolysis/gluconeogenesis (fig. S9) and the urea cycle (fig. S10) based on “Glycolysis/gluconeogenesis” and “Arginine biosynthesis” in KEGG database (*55–57*), respectively. We focused on these pathways because the pairwise Pearson’s correlation coefficients among molecules in “Glycolysis/gluconeogenesis” and “Arginine biosynthesis” (including urea cycle) significantly decreased in *ob*/*ob* mice (Fig. 7**C**). Consistent with the disrupted *t*-half values in the manually reconstructed map of fatty acid degradation (fig. S7C), the pairwise Pearson’s correlation coefficients in “Fatty acid degradation” also significantly decreased in *ob*/*ob* mice (Fig. 7**C**).

We manually reconstructed major metabolic reactions for glycolysis/gluconeogenesis based on “Glycolysis/gluconeogenesis” (mmu00010) in KEGG database (*55–57*) for WT (fig. S9**A**) and *ob*/*ob* mice (fig. S9**B**). We counted the number of edges from each regulator and found that ATP and AMP regulate various metabolic reactions in glycolysis/gluconeogenesis in WT mice (fig. S9**A**, and **C**), while molecules with fewer edges than ATP and AMP regulate the metabolic reactions in glycolysis/gluconeogenesis in *ob*/*ob* mice (fig. S9**B**, and **D**). We assessed the degree distribution of the network by a log-log scatter plot and found that degree distribution was similar between WT and *ob*/*ob* mice (fig. S9**E**) in glycolysis/gluconeogenesis, consistent with the case of the whole starvation-responsive metabolic network (Fig. 5**A**). We assessed the temporal order of the starvation-responsive metabolic network in glycolysis/gluconeogenesis by comparing the *t*-half and degree of responsive molecules (figs. S9**F** and **G**). Although it was not statistically significant, we found a weak negative correlation between *t*-half and the degree of responsive molecules in glycolysis/gluconeogenesis of WT mice (figs. S9**F**), indicating that the molecules connected with many edges responded rapidly to starvation (*t*-half values of nodes with log10-transformed degree more than 0.6 were small). By contrast, the correlation coefficient between *t*-half and the degree of molecules was slightly positive in *ob*/*ob* mice (fig. S9**G**), suggesting that the temporal order of the molecules was disrupted in *ob*/*ob* mice. We also calculated the pairwise Pearson’s correlation coefficients for time courses of each pair of molecules (including phosphorylation events) in glycolysis/gluconeogenesis in WT and *ob*/*ob* mice (fig. S9**H**). Consistent with the case of the whole starvation-responsive metabolic network (Fig. 7**B**), the distribution of correlation coefficients in WT mice was bimodal with peaks at around 1 and −0.75, indicating that both positive or negative co-regulation occurred among molecules. By contrast, the distribution of correlation coefficients was unimodal with a broad peak around 0 in *ob*/*ob* mice, suggesting that co-regulation was disrupted in *ob*/*ob* mice.

Taken together, the starvation-responsive metabolic network in glycolysis/gluconeogenesis is regulated by highly connected hub molecules such as ATP and AMP in WT mice, but by molecules connected with fewer edges than ATP or AMP in *ob*/*ob* mice (figs. S9**C** and **D**). The network was structurally robust (fig. S9**E**) but temporally vulnerable (figs. S9**F**, **G**, and **H**) in liver of WT and *ob*/*ob* mice.

We manually reconstructed major metabolic reactions for urea cycle based on “Arginine biosynthesis” (mmu00220) in KEGG database (*55–57*) (fig. S10**A**). We counted the number of edges from each regulator in WT (figs. S10**B)** and *ob*/*ob* (fig. S10**C**) mice. We found that Glu and ATP, both of which were WT-specific hub nodes in the starvation-responsive metabolic network (Fig. 5**C**), regulated various metabolic reactions in WT mice (fig. S10**B**). In *ob*/*ob* mice, succinate, an *ob*/*ob*-specific hub node in the starvation-responsive metabolic network (Fig. 5**D**), and citrulline regulated various metabolic reactions in *ob*/*ob* mice (fig. S10**C**). We assessed the degree distribution of the network by a log-log scatter plot and calculated the scaling parameters in WT and *ob*/*ob* mice (fig. S10**D**). In contrast to the case of the whole starvation-responsive metabolic network (Fig. 5**A**) and glycolysis/gluconeogenesis (fig. S9**E**), we found that the scaling parameter in *ob*/*ob* mice was smaller than that in WT mice (fig. S10**D**), suggesting that the degree distribution was altered and the proportion of nodes connected with many edges relatively increased in *ob*/*ob* mice. This result suggests that the structure of the starvation-responsive metabolic network was altered in urea cycle of *ob*/*ob* mice. We assessed the temporal order of the starvation-responsive metabolic network in urea cycle by comparing the *t*-half and the degree of responsive molecules (figs. S10**E** and **F**). We found a significant negative correlation between *t*-half and the degree of molecules in WT mice (fig. S10**E**), indicating that the molecules connected with many edges responded rapidly to starvation in urea cycle. By contrast, this significant correlation between *t*-half and the degree of molecules was lost in *ob*/*ob* mice (fig. S10**F**), suggesting that the temporal order of the molecules was disrupted in *ob*/*ob* mice. We also calculated the pairwise Pearson’s correlation coefficients for time courses of each pair of molecules (including phosphorylation events) in urea cycle in WT and *ob*/*ob* mice, respectively (fig. S10**G**). The distribution of correlation coefficients in WT mice was bimodal with peaks at around 1 and −0.6, indicating both positive and negative co-regulation occurred among molecules in urea cycle. The distribution of correlation coefficients in *ob*/*ob* mice was also bimodal but the peaks of the distribution were at around 0.5 and −0.3, suggesting that co-regulation among molecules was disrupted in *ob*/*ob* mice. Taken together, the starvation-responsive metabolic network in urea cycle was both structurally and temporally altered in *ob*/*ob* mice, suggesting the severe disruption of amino acid metabolism in *ob*/*ob* mice.

It should be noted that the distribution of correlation coefficients (Fig. 7**B**) could also be affected by the threshold for significance of the responses to starvation, because the distribution of correlation coefficient among time courses of all measured molecules was unimodal both in WT and *ob*/*ob* mice (fig. S11).

To obtain a qualitative landscape of co-regulations at a molecular level, we constructed networks of molecules that exhibited strong co-regulations (the absolute value of Pearson’s correlation coefficient > 0.9, see fig. S12) among metabolites, lipids, FFA & acyls, enzyme_protein, and enzyme_PTM in WT mice (Fig. 7**D**) and *ob/ob* mice (Fig. 7**E**), named as the correlation metabolic network (Data File S7). Note that the correlation metabolic network is data-dependent and the network structure highly depends on the threshold for the absolute value of correlation coefficient (fig. S12).

The correlation metabolic network in WT mice included a single large cluster comprised of 275 molecules, representing molecules with various biological functions such as carbohydrate metabolism and fatty acid degradation (Fig. 7**D**), indicating that most of the responsive molecules in WT mice are co-regulated. ATP and AMP, the rapid hub molecules in the starvation-responsive metabolic network (Fig. 5 and 6) were included in the large cluster, possibly suggesting that ATP and AMP formed this large cluster by regulating the large number of metabolic reactions. By contrast, the network in *ob*/*ob* mice was fragmentated with some clusters containing functionally related molecules (Fig. 7**E**). Taken together, temporal co-regulation of metabolism in response to starvation was disrupted in the *ob/ob* mice.

### Functional analysis of transcriptomic and proteomic data for potential clinical application

Dietary restriction, including caloric restriction (reducing total calorie intake) and intermittent fasting (setting fasting periods in daily life with or without reducing the total calorie intake), extends lifespan and improves health in multiple species (*6*, *31*, *70*, *71*). However, obesity cancels the effects of intermittent fasting, that is a permissive form of dietary restriction considering the compliance, to extend lifespan (*31*).

Investigating a therapeutic target to treat the negative effects of obesity on intermittent fasting is clinically important. To achieve this, comprehensive and unbiased measurements are necessary, because manually measured phenotypic traits failed to explain 61% of the diet to lifespan covariance (*31*). Intermittent fasting, a long-term and repetitive form of starvation, is expected to include the cumulative effects of short-term starvation up to 24 hours. Thus, analyzing our omics data on short-term starvation is valuable, particularly to reveal the differences of starvation-induced changes between WT and *ob*/*ob* mice. Since metabolic traits associated with lean tissue mass explained only 3.8% of the lifespan covariance (*31*), we focused on general functions of liver using our transcriptomic and proteomic data, rather than on metabolism. In addition, we used MSigDB (*72*), a more generalized database than KEGG (*55–57*).

To functionally characterize the starvation-responsive transcripts, we conducted an enrichment analysis of significantly changed (either increased or decreased) transcripts in WT and *ob*/*ob* mice (Fig. 8**A** and table Data File S8). Hallmarks, which are categories of molecules in MSigDB (*72*), related to immune responses, such as TNFA_SIGNALING_VIA_NFKB, INTERFERON_ALPHA_RESPONSE, and INTERFERON_GAMMA_RESPONSE; metabolism such as CHOLESTEROL_HOMEOSTASIS, FATTY_ACID_METABOLISM, and PEROXISOME; stress responses such as HYPOXIA, APOPTOSIS, and UNFOLDED_PROTEIN_RESPONSE; and cellular signaling such as MTORC1_SIGNALING, P53_PATHWAY, and KRAS_SIGNALING_DN were identified as commonly enriched hallmarks in both WT and *ob*/*ob* mice (Fig. 8**A** and Data File S8). Hallmarks such as ESTROGEN_RESPONSES_EARLY, ESTROGEN_RESPONSES_LATE, and GLYCOLYSIS were identified as WT-specifically enriched hallmarks (Fig. 8**A** and Data File S8). MYC_TARGET_V1 and MYC_target_V2 were identified as *ob*/*ob*-specifically enriched hallmarks (Fig. 8**A** and Data File S8). The common enrichment of the hallmarks related to immune responses in both WT and *ob*/*ob* mice suggest that the changes of the immune repertoire observed with dietary restriction (*31*) also occurred with short-term fasting within 24 hours.

**Fig. 8:**
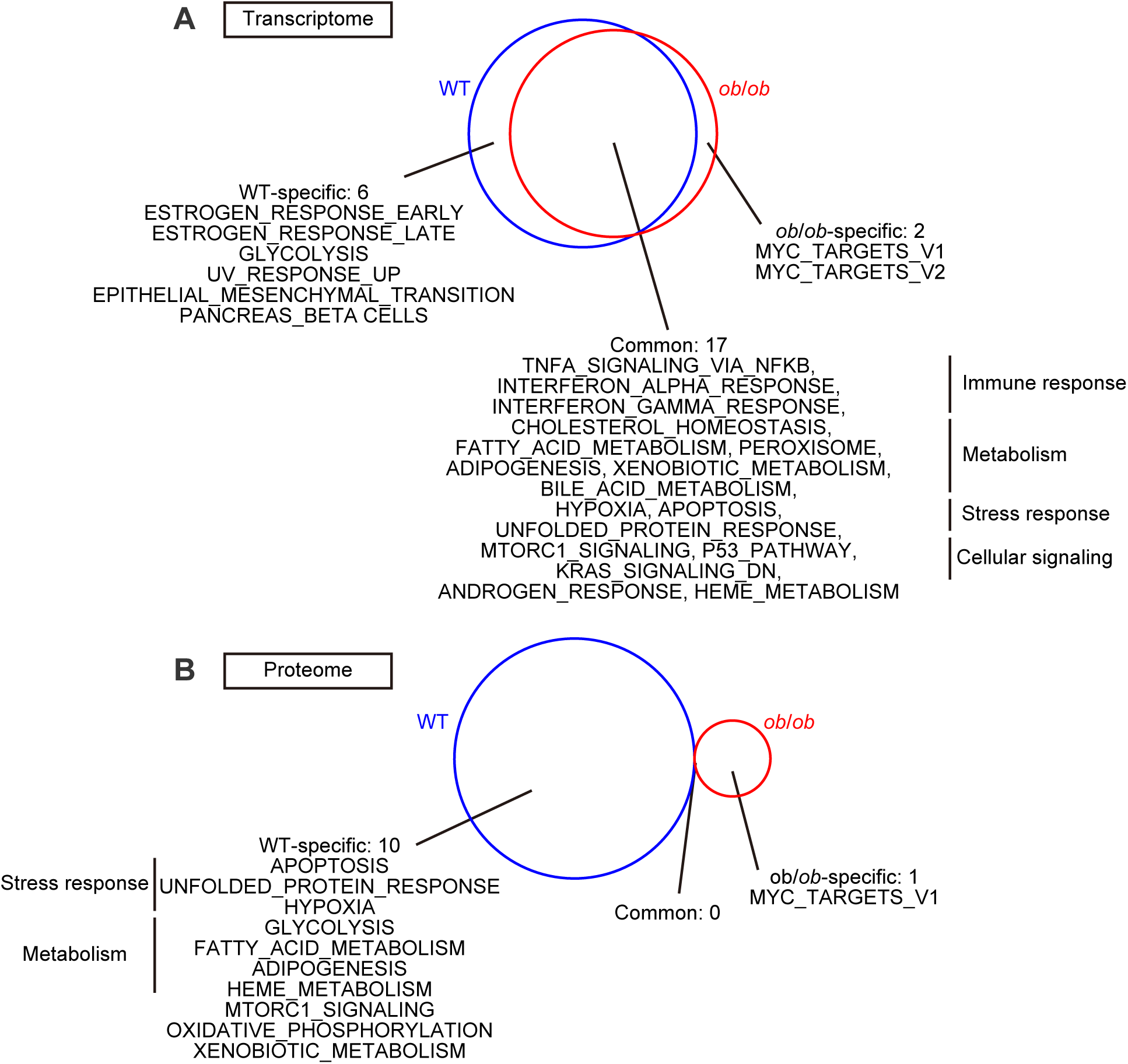
Functional analysis of transcriptomic and proteomic data for potential clinical application (**A**) Enrichment analysis of significantly changed transcripts in WT (blue circle) and *ob*/*ob* (red circle) mice using MSigDB. (**B**) Enrichment analysis of significantly changed proteins in WT (blue circle) and *ob*/*ob* (red circle) mice using MSigDB.

To functionally characterize the starvation-responsive proteins, we conducted an enrichment analysis of significantly changed proteins in WT and *ob*/*ob* mice (Fig. 8**B** and Data File S8). Hallmarks related to stress responses such as APOPTOSIS and UNFOLDED_PROTEIN_RESPONSE; metabolism such as GLYCOLYSIS and FATTY_ACID_METABOLISM; MTORC1_SIGNALING; and OXIDATIVE_PHOSPHORYLATION were identified as WT-specifically enriched hallmarks. MYC_TARGETS_V1 was identified as an *ob*/*ob*-specifically enriched hallmark. No hallmark was commonly enriched in both WT and *ob*/*ob* mice, suggesting that the responses to starvation are functionally distinct between both genotypes in proteomic data.

Taken together, commonly enriched hallmarks were most abundant in transcriptomic data, while WT-specifically responsive hallmarks were most abundant in proteomic data, suggesting that the responses to starvation in WT and *ob*/*ob* mice were functionally similar in the transcriptome, but totally different in the proteome. This disconnection between responses at the mRNA and protein levels in *ob*/*ob* mice could be due to deficiencies in protein translation and degradation in *ob*/*ob* mice (fig. S8). Therefore, targeting deficiencies in protein translation and degradation could be a potential therapeutic target to treat the insensitivity of intermittent fasting in obese mice (*31*).

## Discussion

We measured multiomic data during starvation in WT and *ob*/*ob* mice (Fig. 2) and constructed a global starvation-responsive transomic network by integrating these data (Fig. 3). We also showed that the starvation-responsive metabolic network (Fig. 4), a sub network of the global starvation-responsive transomic network, is a scale-free-like network. The hub molecules of the networks, regulating many metabolic reactions, were energy indicators, such as ATP and AMP in WT mice, and less highly connected hub molecules, such as lipids in *ob*/*ob* mice (Fig. 5). Despite the loss of the responsiveness of the hub energy indicators in *ob*/*ob* mice, structural properties of the network in *ob*/*ob* mice were maintained (Fig. 5), likely through the maintenance of nodes with relatively large degrees (Fig. S5). By contrast, the temporal properties including the temporal order (Fig. 6) and the temporal co-regulation of molecules were disrupted in *ob*/*ob* mice (Fig. 7). Similar to the whole starvation-responsive metabolic network, the starvation responsive metabolic network of glycolysis/gluconeogenesis in *ob*/*ob* mice was structurally maintained but temporally disrupted compared to the network in WT mice (fig. S9). However, the starvation-responsive metabolic network of urea cycle in *ob*/*ob* mice was both structurally and temporally disrupted comparing to the network in WT mice (fig. S10). In addition, we proposed a potential clinical target to treat the negative effects of obesity on the intermittent fasting (Fig. 8).

The maintenance of the structural properties of the starvation-responsive metabolic network in *ob*/*ob* mice (Fig. 5) suggests the robustness of the metabolic networks against pathological perturbations. Robustness of the structural properties of metabolic networks have also been investigated in cancers (*41*, *42*). Structural properties of metabolic networks based on the tissue-specific gene expression were similar between 15 cancers and corresponding normal tissues (*41*). Metabolic network based on flux correlations showed similar structural properties between normal and cancer tissues (*42*). These two studies suggest that structural properties of metabolic networks in cancer are similar to those of normal tissues, despite profound differences in gene expression patterns of cancer and non-cancer cells (*73*). These studies, together with our results, indicate that structural properties of metabolic networks are robust among various metabolic perturbations including starvation in healthy or obese conditions and cancers.

By contrast to the robust structural properties (Fig. 5), temporal properties, which were maintained depending on the hub molecules including energy indicators such as ATP and AMP in WT mice, were disrupted in *ob*/*ob* mice (Figs. 6 and 7). These results highlight the significance of the highly connected hub molecules for temporal properties of the starvation-responsive metabolic networks. If a small number of hub molecules regulate the entire network, nodes in the network would be uniformly regulated, resulting in a temporally ordered and co-regulated network. Thus, the starvation-responsive metabolic network is temporally disrupted in *ob*/*ob* mice, that lost the responsiveness of the highly connected hub molecules such as ATP and AMP.

Structural and temporal properties of the starvation-responsive metabolic network were evaluated for specific metabolic pathways, such as glycolysis/gluconeogenesis (fig. S9) and urea cycle (fig. S10). The starvation-responsive metabolic network of glycolysis/gluconeogenesis in *ob*/*ob* mice was structurally maintained but temporally disrupted, consistent with the whole starvation-responsive metabolic network (fig. S9). However, the starvation-responsive metabolic network of urea cycle in *ob*/*ob* mice was disrupted both structurally and temporally (fig. S10), suggesting a strong disruption of amino acid metabolism in *ob*/*ob* mice during starvation. Amino acids during starvation are supplied by autophagy (*74*), a protein degradation process that is disrupted in *ob*/*ob* mice (fig. S8**C** and **D**). Therefore, a disrupted amino acid supply caused by autophagy deficiency could result in severely disrupted amino acid metabolism during starvation in *ob*/*ob* mice.

If the metabolic network is temporally vulnerable, the structural robustness observed in starvation (several hours) and cancers (∼years) could be disrupted in a rapid metabolic shift such as glucose ingestion (minutes) because of the lack of time for rescuing the structural properties. Previously, we conducted a transomic analysis of liver in WT and *ob*/*ob* mice during oral glucose ingestion, in which we measured metabolome, lipidome, and transcriptome (*37*, *39*). Small fraction of the responsive metabolites (15 out of 88, 17.0%) and transcripts (732 out of 5,091, 14.4%) was commonly responsive in WT and *ob*/*ob* mice during oral glucose ingestion, while more than one-third of the responsive molecules were commonly responsive during starvation (Fig. 2) (*37*). Scaling parameters of the transomic networks, representing the heterogeneity of the network, were different during oral glucose ingestion but similar during starvation between WT and *ob*/*ob* mice (*39*). Taken together, glucose ingestion requires distinct responsive molecules and heterogeneities of transomic network between WT and *ob*/*ob* mice, suggesting metabolic network is more vulnerable to temporally rapid perturbations (oral glucose ingestion) than slow (starvation) or chronic (cancer) perturbations.

By evaluating the temporal order among omic layers during starvation (Fig. 6), we found that metabolite layer was the fastest even compared to the enzyme_PTM layer, which is composed of protein phosphorylation events (Fig. 6**G**). Our previous study showed that rapid metabolic regulation by metabolites also occurs in the liver of WT mice during oral glucose ingestion (*37*). However, we did not measure the phosphoproteome in that study (*37*), thus we cannot ascertain if metabolite regulation precedes phosphorylation-mediated regulation. Because here we measured both the metabolome and phosphoproteome, our data indicated that metabolites take the initiating role in controlling the transomic layers during starvation in liver.

One of the limitations of this study is the accuracy of the inference of regulator activities, including TFs and protein kinases, and regulatory relationships from metabolites, lipids, and FFA & acyls to metabolic reactions. For inference of TF activity, we failed to identify Foxo1, which is master regulator of liver metabolism and is necessary for upregulation of glucose production during starvation (*75*). We can explain this discrepancy because we removed data for Foxo1 in liver from the data obtained from the ChIP-Atlas database (*76*, *77*) as not meeting our quality check process for inference input (see Methods). Another example, Creb1, which has been known to be upregulated during starvation (*5*), was classified as a down-regulated TF (Table 5), probably due to the failure of detection in phosphorylation, that is important for Creb1 activity (*5*). For kinase inference, we failed both mTorc1 and Ampk as responsive protein kinases, both of which are known to be important for adaptation to starvation (*78*, *79*). Indeed, phosphorylation of Ampk was confirmed in our dataset by western blotting (Fig. 5**K** and **L**), but not our phosphoproteome data, suggesting insufficient comprehensiveness and quantitative capacity of our phosphoproteome data. In addition, protein kinases were identified for less than 5% of phosphorylation events at present (*80*), thus the prediction algorithm could be inaccurate. For regulatory relationships including those from metabolites, lipids, and FFA & acyls to metabolic reactions, we identified regulatory relationships that could be extracted from existing databases. Thus, unfamiliar or unknown regulatory relationships not registered in the databases we used were not included in our study. These limitations originate from incomprehensiveness of omics measurement (for example, lack of phosphorylated Creb1 and Ampk) and database (for example, lack of Foxo1). These limitations could be partially solved by recently developed techniques of measurements such as LiP-SMAP (*81*), PROMIS (*82*), and MIDAS (*83*) for protein-metabolite (including lipids and FFA & acyls) interaction, LiP-MS (*84*) for post-transcriptional regulations including phosphorylation, and integrating recently developed bioinformatic tools such as CollectTRI-based database (*85*), The Kinase Library (*86*), and OmniPath (*87*) to transomic analyses for algorithm, but it is still challenging to approach these limitations especially in vivo at present. Therefore, we need some breakthrough for accurate inferences of regulator activities.

Another limitation of our study is that we only used one model of obesity. By only using *ob*/*ob* mice, we cannot exclude the possibility that the results are caused by the effect of leptin deficiency (*88*). We compared our results with the previous study using high fat diet (HFD) as a model for obesity (*30*). We observed similar changes in metabolic enzymes and in glycogen during starvation in WT and *ob/ob* mice (fig. S1) as were reported for both lean and HFD mice (*30*). Thus, our results are related to obesity not leptin-deficiency itself.

In addition, our experimental condition is not optimized for dietary restriction, because we focused on adaptation to starvation. Although we proposed that disturbed protein translation and degradation are potential therapeutic targets to treat the negative effects of obesity on intermittent fasting to extend lifespan (*31*), validation is needed in studies focusing on dietary restriction under well-controlled experimental conditions.

Despite several limitations, our present study successfully described the global landscape of adaptation to starvation in WT and *ob*/*ob* mice and suggested that the highly connected hub energy indicators temporally coordinate the starvation-responsive metabolic network in the healthy condition and that these critical hubs are no longer starvation-responsive in obesity, resulting in temporal disruption of the starvation-responsive metabolic network. Robustness of the structures against pathological perturbations could be a general nature of metabolic networks, combining with the previous studies in cancer (*41*, *42*). Vulnerability of temporal properties of the network in *ob*/*ob* mice could be helpful to understand the pathology of adaptive responses to starvation in obesity.

## Materials and Methods

### Mouse experiments

All the animal experiments were conducted after approval by the animal ethics committee of the University of Tokyo. Ten-week-old C57BL/6 WT and *ob*/*ob* male mice were purchased from Japan SLC Inc. Mice were starved for 0, 2, 4, 6, 8, 12, 16, 24 hours, starting at approximately 18:00. For omic measurement, mice were euthanized by cervical dislocation, blood was collected from retro-orbital sinus into tubes containing 0.5 mg of EDTA, and liver was dissected and immediately frozen in liquid nitrogen. Blood was centrifuged at 2,300 × *g* for 15 min at 4°C to separate plasma.

Plasma was frozen in liquid nitrogen and stored at −80°C. The frozen liver was crushed with dry ice to a fine powder with a Wonder Blender (WB-1, OSAKA CHEMICAL CO., LTD). Blood glucose was determined using ACCU-CHECK (Roche).

### Metabolomic analysis

Metabolomic analysis was conducted as previously described (*37*). For liver, total metabolites were extracted with methanol:chroloform:water (2.5:2.5:1) along with total proteins. Approximately 40 mg of liver was homogenized in 500 µl of ice-cold methanol containing internal standards [20 μM l-methionine sulfone (Wako), 2-morpholinoethanesulfonic acid, monohydrate (Dojindo), and d-camphor-10-sulfonic acid (Wako)] for normalization of peak intensities of mass spectrometry (MS) among runs. Next, 500 μL of chloroform and then 200 μL of water were added and mixed.

After centrifugation at 4,600 × *g* for 15 min at 4°C, the aqueous layer was filtered through a 5 kDa cutoff filter (UFC3LCCNB-HMT, Human Metabolome Technologies) to remove protein contamination. Simultaneously, proteins were precipitated from the interphase and organic layers by addition of 800 μL of ice-cold methanol followed by centrifugation at 12,000 × *g* for 15 min at 4°C. The pellet was washed with 1 mL of ice-cold 80% (v/v) methanol and resuspended in 500 μL water, followed by sonication with Bioruptor (UCW-310, COSMO BIO). By addition of 500 μL of 2% SDS and 100 mM Tris-HCl pH8.8, the pellet was lysed for 1 hour at 4℃ with gentle mixing followed by sonication using Handy Sonic (UR-20P, TOMY). The concentration of total protein was determined with the bicinchoninic acid (BCA) assay (23227, Thermo Fisher Scientific) and used for normalization of concentration among samples.

For plasma, 40 μL of the samples were used to extract metabolites with methanol:chloroform:water (2.5:2.5:1). The samples were dissolved in 400 μL of ice-cold methanol containing the internal standards [20 μM l-methionine sulfone (Wako), 2-morpholinoethanesulfonic acid, monohydrate (Dojindo), and d-camphor-10-sulfonic acid (Wako)], followed by addition of 400 μL of chloroform, and then 120 μL of water. After centrifugation at 10,000 × *g* for 3 min at 4 °C, the aqueous layer was filtered through a 5 kDa cutoff filter (UFC3LCCNB-HMT, Human Metabolome Technologies) to remove protein contamination.

The filtrate was lyophilized and suspended in 50 μL of water containing the reference compounds [200 μM each of trimesate (206-03641, Wako) and 3-aminopyrrolidine (404624, Sigma-Aldrich)]. Metabolites were quantified using an Agilent 1600 Capillary Electrophoresis system (Agilent technologies), a G1603A Agilent CE-MS adapter kit, and a G1607A Agilent CE electrospray ionization (ESI)– MS sprayer kit. For cationic compounds, a fused silica capillary [50 μm internal diameter (i.d.) × 100 cm] was used with 1 M formic acid as the electrolyte (*89*).

Methanol/water (50%, v/v) containing 0.01 μM hexakis (2,2-difluoroethoxy) phosphazene was used as sheath liquid and delivered at 10 μl/min. ESI–time-of-flight (TOF) MS was conducted in positive ion mode, and the capillary voltage was set to 4 kV. Each acquired spectrum was recalibrated automatically using the masses of the reference standards [^13^C isotopic ion of a protonated methanol dimer (2 MeOH + H)]^+^, mass/charge ratio (*m*/*z*) 66.0631 and [hexakis(2,2-difluoroethoxy)phosphazene + H]^+^, *m*/*z* 622.0290. For identification of metabolites, relative migration times of all peaks to the reference compound 3-aminopyrrolidine were calculated, followed by comparison of their *m*/*z* values and relative migration times to the metabolite standards. For quantification of the metabolites, peak areas were compared to calibration curves generated using internal standardization techniques with methionine sulfone. The other parameters and conditions were the same as previously described (*90*). For anionic metabolites, COSMO (+) (chemically coated with cationic polymer) capillary (50 μm i.d. by 105 cm) (Nacalai Tesque, Kyoto, Japan) was used with a 50 mM ammonium acetate solution (pH 8.5) as the electrolyte. Methanol/5 mM ammonium acetate (50%, v/v) containing 0.01 μM hexakis (2,2-difluoroethoxy) phosphazene was used as sheath liquid and delivered at 10 μl/min. ESI-TOFMS was conducted in negative ion mode, and the capillary voltage was set to 3.5 kV. Trimesate and d-camphor-10-sulfonic acid were used as the reference and the internal standards, respectively. The other parameters and conditions were same as described previously (*91*). The Agilent MassHunter software (Agilent technologies) was used for data analysis (*89–91*).

F1,6P and F2,6P were measured separately using IC-QEMS (*92*). A Dionex IonPac AS11-HC-4 µm column (250 × 0.4 mm, 4 µm; Thermo Fisher Scientific) was used at 35°C to separate metabolite. We used KOH as an eluent at the speed of 0.02 mL/min. The gradient was as follows: 1 mM from 0 to 2 min, 20 mM at 16 min, 100 mM at 35 min. As a sheath solution, isopropanol containing 0.1% acetic acid was used at the speed of 5 µl/min. MS was performed in the ESI negative-ion mode. The parameters were as follows: sheath gas, 20 (arbitrary units); auxiliary gas, 10 (arbitrary units); spray voltage, 4.0 kV; capillary temperature, 300°C; S-lens, 50 (arbitrary units). Data acquisition was conducted in full MS scan mode. Parameters of the scanning were as follows: resolution, 70,000; auto-gain control target, 3×10^6^; maximum ion injection time, 100 ms; scan range, 70–1,000 *m*/*z*.

### Lipidomic analysis

Liver samples were prepared for lipid extraction using the Bligh and Dyer’s method (*93*) with minor modifications. Briefly, the lipids and their related metabolites were extracted from frozen and crushed liver tissues (∼50 mg) with 1 ml of cold methanol. The samples were vigorously mixed for 1 min and sonicated for 5 min. The extracts were then centrifuged at 16,000 ×*g* for 5 min at 4°C, and the resultant supernatant was collected. Protein concentrations in the pellet were determined using the bicinchoninic acid (BCA) assay (23227, Thermo Fisher Scientific). The collected supernatant (180 µL) was mixed with 200 µL of chloroform, 150 µL of water, 10 µL of internal standard (IS) A (Mouse SPLASH Lipidomix Mass Spec Standard, Avanti Polar Lipids Inc., Alabaster, AL, USA) containing 1.0 nmol phosphatidylcholine (PC) 15:0–18:1 (d7), 0.070 nmol phosphatidylethanolamine (PE) 15:0–18:1 (d7), 0.20 nmol phosphatidylserine (PS) 15:0–18:1 (d7), 0.050 nmol phosphatidylglycerol (PG) 15:0–18:1 (d7), 0.20 nmol phosphatidylinositol (PI) 15:0–18:1 (d7), 0.10 nmol phosphatidic acid (PA) 15:0–18:1 (d7), 0.45 nmol lysophosphatidylcholine (LPC) 18:1 (d7), 0.020 nmol lysophosphatidylethanolamine (LPE) 18:1 (d7), 2.5 nmol cholesteryl ester (CE) 18:1 (d7), 0.15 nmol diacylglycerol (DG) 15:0–18:1 (d7), 0.35 nmol triacylglycerol (TG) 15:0–18:1 (d7)–15:0, and 0.20 nmol sphingomyelin (SM) d18:1–18:1 (d9), 10 µL of IS B (Avanti Polar Lipids Inc.) containing 1.0 nmol monoacylglycerol (MG) 18:1 (d7), 0.10 nmol ceramide (Cer) d18:1 (d7)–15:0, 0.10 nmol hexosylceramide (HexCer) d18:1 (d7)–18:1, 1.1 nmol free fatty acid (FA) 16:0 (^13^C16), and 3.0 nmol cholesterol (d7), and 10 µL of IS C (Merck, Darmstadt, Germany) containing 0.40 nmol acylcoenzyme A (acyl-CoA) 2:0 (^13^C^2^). The aqueous and organic layers were separated by vortexing and subsequent centrifugation at 16,000 ×*g* and 4°C for 5 min.

The aqueous (upper) layer (250 µL) was transferred to a clean tube for analysis by a liquid chromatography (Nexera X2 UHPLC system, Shimadzu Co., Kyoto, Japan) with a metal-free peek-coated InertSustain C18 column (2.1 mm i.d. × 150 mm, 3 µm particle size, GL Sciences Inc., Tokyo, Japan) coupled with a Q Exactive, high-performance benchtop quadrupole Orbitrap high-resolution tandem mass spectrometer (Thermo Fisher Scientific) (C18-LC/MS) for acyl-CoAs and acyl-carnitines (*94*). After the aqueous layer extracts were evaporated under vacuum, dried extracts were stored at −80°C until use for C18-LC/MS analysis. Prior to analysis, the dried aqueous layer was reconstituted in 50 µL of water.

The organic (lower) layer (140 µL) was transferred to a clean tube for analysis by supercritical fluid chromatography (SFC, Nexera UC system, Shimadzu Co) with an ACQUITY UPC^2^ HSS C18 column (3.0 mm i.d. × 100 mm, 1.8 µm particle size, Waters, Milford, MA, USA) coupled with a triple quadrupole mass spectrometry (TQMS, LCMS-8060, Shimadzu Co.) (C18-SFC/MS/MS) for FAs, TGs, and CEs (*94*) and an SFC (Shimadzu Co.) with a ACQUITY UPC2 Torus diethylamine (DEA) (3.0 mm i.d. × 100 mm, 1.7 µm particle size, Waters) coupled with a TQMS (DEA-SFC/MS/MS) for PCs, PEs, PSs, PGs, PIs, PAs, LPCs, LPEs, MGs, DGs, SMs, cholesterol, Cers, and HexCers (*94*, *95*). The liver lipid extract (140 µL) was diluted to a final volume of 200 µL with a solution of methanol for C18-SFC/MS/MS and DEA-SFC/MS/MS analysis.

### Transcriptomic analysis

Total RNAs were extracted from approximately 10 mg of liver using RNeasy Mini Kit (74106, QIAGEN) and QIAshredder (79656, QIAGEN). The amount and quality of RNAs were assessed using Nanodrop (Thermo Fisher Scientific) and the 2100 Bioanalyzer (Agilent Technologies). The cDNA libraries were prepared using the TruSeq Stranded mRNA Kit (Illumina, San Diego, CA, USA). The resulting cDNAs were subjected to 150 base pair (bp) paired-end sequencing on an Illumina NovaSeq6000 Platform (Illumina). Sequences were aligned to the reference genome of mice derived from the Ensembl database (*96*, *97*) (GRCm38/mm10, Ensembl release 97) using the STAR software package (v.2.5.3a) (*98*). The RSEM tool (*99*) (v.1.3.0) was used to estimate gene expression levels from the aligned sequences. Resultant read counts were used as inputs for edgeR software (*49*). For other analyses, transcripts per kilobase million (TPM) were used.

### Proteomic analysis

Total proteins extracted during sample preparation of metabolomic analysis were subjected to the following proteomic analysis. For standardization among runs of measurement, proteins from frozen liver powder of Lys(6)-SILAC mice (Silantes) were also extracted with methanol/chloroform extraction. Protein concentration of the lysates was determined using the bicinchoninic acid (BCA) assay (23227, Thermo Fisher Scientific) and calibrated to 1 mg/mL. Then, 50 µg each of non-labeled and labeled lysates were mixed for peptide preparation. Cysteine residues were blocked by 2 mM Tris (2-carboxyethyl) phosphine hydrochloride (TCEP) (Thermo Fisher Scientific) at 37°C for 30 min followed by alkylation with 10 mM 2-iodoacetamide (IAA) at room temperature for 30 min. The proteins were precipitated with acetone for 3 hours at −30°C and the resulting pellet was dispersed in 50 mM triethylammonium bicarbonate by ultrasonic treatment (three times for 30 s with intervals of 30 s) with a Bioruptor (Diagenode). The protein suspension was subjected to digestion with lysyl endopeptidase (Wako) for 16 h at 37°C. Resulting peptides were centrifuged at 15,000 × *g* for 15 min at 4°C and subjected to C18-StageTip purification (*100*). Peptides were resolved in 50 µL 3% acetonitrile (can)-0.1% foromic acid prior to MS analysis.

All samples were analyzed with Q Exactive HF-X (Thermo Fisher Scientific) instrument equipped with an UltiMate 3000 RSLCnano LC System (Thermo Fisher Scientific) via a nano-electrospray source with a column oven set at 50°C (AMR Inc.). Peptides were directly injected onto a 75 μm × 25 cm PicoFrit emitter (New Objective, Woburn, MA, USA) packed in-house with C18 core-shell particles (CAPCELL CORE MP 2.7 μm, 160 Å material; Osaka Soda Co., Ltd., Osaka, Japan) with a linear gradient of 7–42% B for 76 min, 42-80% B for 5 min, and 80% B for 9 min at a flow rate of 100 nL/min, where A is 0.1% formic acid and B is 0.1% formic acid and 80% acetonitrile. MS data acquisition was performed in data-independent acquisition (DIA) mode. For quantification in the DIA mode, MS1 spectra were collected in the range of *m*/*z* 423-865 at 60,000 resolution to set an AGC target of 3 × 10^6^. For MS2 spectra, collections were conducted in the range of > *m*/*z* 200 at 30,000 resolution to set an AGC target of 3 × 10^6^. The isolation width was configured to 8 Th with stepped normalized collision energies of 20, and 23%. Through the optimization by Skyline 4.1 (*101*), isolation window patterns in *m*/*z* 428-860 were used as window placements. In the DIA mode for a spectral library, the pool of all samples was analyzed by using the above measurements and the gas-phase fractionation method. In the gas-phase fraction method, six MS ranges (*m*/*z* 426-502, 498-574, 570-646, 642-718, 714-790, and 786-862) were used, and each was measured by overlapping DIA (*102*, *103*). MS1 spectra were collected at 120,000 resolution to set an AGC target of 3 × 10^6^. MS2 spectra were collected in the range of > *m*/*z* 200 at 60,000 resolution to set an AGC target of 3 × 10^6^. The isolation width was set to 2 Th with stepped normalized collision energies of 20 and 23%. Isolation window patterns in the ranges of *m*/*z* 428-500, 500-572, 572-644, 644-716, 716-788, and 788-860 were used as window placements optimized by Skyline 4.1.

The spectra library was generated by searching MS data in the library against the mouse Ensembl release 97 database using Scaffold DIA (Proteome Software Inc., Portland OR, USA). The parameters were as follows: experimental data search enzyme, LysC/P; maximum missed cleavage sites, 1; precursor mass tolerance, 6 ppm; fragment mass tolerance, 8 ppm; and static modification, cysteine carbamidomethylation. The threshold for identification of peptides was a peptide false discovery rate (FDR) < 1%. To identify peptides derived from Lys(6)-SILAC mice, a library of identified peptides with a mass increase 6.020129 Da on Lys was built and added to the library obtained above. All raw data was processed with DIA-NN (ver. 1.8) using library search mode using the following parameters: reannotation, Mus_musculus.GRCm38.97.pep.all.fa; modification, UniMod188 (6.020129), and peak-translation. Using the output from DIA-NN software, the SILAC ratios of all assigned fragment ion pairs were calculated, and the median of the SILAC ratios of each protein was used for protein quantification. Proteins assigned to a single ensemble gene ID were used in the following analyses. The SILAC ratios of all the proteins were normalized by the median of the SILAC ratio in each biological sample.

### Phosphoproteomic analysis

Total proteins extracted during sample preparation of metabolomic analysis were subjected to the following phosphoproteomic analysis. As described above, 2 mg of protein solution was subjected to reduction and alkylation by TCEP and IAA, respectively. Then, proteins were precipitated in three volumes of ice-cold acetone and centrifuged at 16,000 × *g* for 2 h. The pellet was rinsed with 90% acetone followed by digestion with 30 µg of trypsin in 600 µl of 100 mM ammonium bicarbonate for 16 h. Thirty-five µl of 10% trifluoroacetic acid (TFA) and 450 µl of ACN were added to the protein digests (1 mg/300 µl). The mixture was centrifuged at 16,000 × *g* and the supernatant was collected into a new tube. Phosphopeptides were enriched using Fe^3+^-IMAC (*104*). NTA-agarose resin (1 mL) (Qiagen) was washed twice with 3 ml of distilled water and 2 ml of 100 mM FeCl3 in 0.1% acetic acid was added. Following a wash with 2 ml of 0.1% TFA, 2% ACN and 0.1% TFA, 60% ACN, the resin was stored in 1 ml of 0.1% TFA, 60% ACN (Fe^3+^-IMAC agarose resin). C18-StageTip (3M) packed with 75 µl of Fe^3+^-IMAC agarose resin was equilibrated with 200 µl of 0.1% TFA, 60% ACN, then the protein digests were loaded onto the IMAC/C18-StageTip and the loaded system was washed twice with 200 µL of 0.1% TFA, 60% ACN. Following equilibration with 200 µl of 0.1% TFA, 2% ACN, 200 µl of 1% inorganic phosphate was added to elute phosphopeptides from the Fe^3+^-IMAC agarose resin and bind them to the C18-StageTip. Following a wash with 200 µL of 0.1% TFA, 2% ACN, the peptides were eluted with 0.1% TFA, 60%ACN from the IMAC/C18-StageTip. The resulting peptides were dried, reconstituted in 3% ACN with 0.1% formic acid, and subjected to MS analysis.

All samples were analyzed with Orbitrap Exploris 480 MS (Thermo Fisher Scientific) instrument equipped with an UltiMate 3000 RSLCnano LC System (Thermo Fisher Scientific) via a nano-electrospray source with a column oven set at 60°C (AMR Inc.). Peptides were directly applied onto a 75 μm × 120 mm column (Nikkyo Technos Co.,Ltd) with a linear gradient of 5–32% B for 70 min and 32-65% B for 10 min at a speed of 200 nL/min, where A is 0.1% formic acid and B is 0.1% formic acid and 80% acetonitrile.

MS data were acquired in overlapping data-independent acquisition (DIA) mode (*102*, *103*). For quantification in the DIA mode, MS1 spectra were collected in the range of *m*/*z* 390-1010 at 30,000 resolution to set an AGC target of 3 × 10^6^. MS2 spectra were collected in the range of > *m*/*z* 200 at 30,000 resolution to set an AGC target of 3 × 10^6^, and stepped normalized collision energies of 22, 26, and 30%. The isolation width for MS2 was set to 10 Th and overlapping window patterns in *m*/*z* 400-1000 were used window placements optimized by Skyline v4.1 (*101*). In the DDA-MS mode for a spectral library, the pool of all samples was analyzed by using the gas-phase fractionation method with MS ranges of *m*/*z* 395-555, 545-705, 695-1005, and 390-1010. MS1 spectra were collected at 120,000 resolution to set an AGC target of 3 × 10^6^. Within the MS ranges of *m*/*z* 395-555, 545-705, 695-1005, the 20 most intense ions with charge states of 2^+^ to 5^+^ more than 5.0 × 10^3^ were fragmented by HCD with normalized collision energies of 22, 26, and 30%, and MS2 spectra were collected in the range of > *m*/*z* 200 at 60,000 resolution to set an AGC target of 5 × 10^5^. Within the MS ranges of *m*/*z* 390-1010, the 40 most intense ions with charge states of 2^+^ to 5^+^ that exceeded 5.0 × 10^3^ were fragmented by HCD with normalized collision energies of 22, 26, and 30%, and MS2 spectra were collected in the range of > *m*/*z* 200 at 30,000 resolution to set an AGC target of 2 × 10^5^.

The spectra library was built by searching MS data in the library against the mouse Ensembl database (GRCm38/mm10, Ensembl release 97) using Proteome Discoverer v2.3 (Thermo Fisher Scientific). The setting parameters were as follows: Fragmentation, HCD; Precursor Tolerance, 8 ppm; Fragment Tolerance, 0.02 Da; Digestion Enzyme, Trypsin; Max Missed Cleavages, 1; Variable Modification, Phospho [S, T, Y]; Fixed Modification, Carbamidomethylation [C]; Site Probability Threshold, 75; Peptide FDR, < 1%. Unique phosphopeptides were subjected to the following analyses. To determine phosphorylation at each phosphosite, the intensities of all signals of quantified phosphopeptides that include the phosphosite of interest were summed.

### Glycogen content assay

Glycogen content in liver was measured using Amplex Red Glucose/Glucose Oxidase Assay Kit glucose assay (Thermo Fisher Scientific) after digestion of glycogen to glucose (*37*, *105*). Approximately 20 mg of crushed tissue samples was incubated for 1 hour at 95°C in 1 mL of 30% (w/v) KOH solution. The lysate (50 μL) was collected into another 1.5 mL tube and neutralized by 15.3 μL of glacial acetic acid. Total protein concentration of the liver digest was determined using the BCA assay (23227, Thermo Fisher Scientific) and the concentration of protein was adjusted to 1 μg/μL. Lipids were removed by the Bligh and Dyer method and glycogen was extracted. Ice-cold methanol (120 μL), 50 μL of chloroform, 10 μL 1% (w/v) of linear polyacrylamide, and 70 μL of water were added to the liver digest (50 μL) and mixed. The mixture was incubated for 30 min on ice and centrifuged at 12,000 × *g* to remove the aqueous layer. The glycogen was precipitated by adding 200 μL of methanol, followed by centrifugation at 12,000 × *g* for 30 min at 4°C. The pellet was washed with ice-cold 80% (v/v) methanol and dried completely. Glycogen pellets were resuspended in 20 μL of amyloglucosidase (0.1 mg/mL; Sigma-Aldrich) in 50 mM sodium acetate buffer and incubated for 2 hours at 55°C. The glycogen content was determined using the Amplex Red Glucose/Glucose Oxidase Assay Kit glucose assay (Thermo Fisher Scientific) following the manufacture’s protocol.

### Western blotting analysis

Total proteins extracted during sample preparation of metabolomic analysis were boiled in sample buffer (58.28 mM Tirs-HCl, pH 6.8, 4.7% glycerol, 2.82% SDS, 6% β-Mercaptoethanol, and 0.0094% bromophenol blue). Samples were subsequently separated by SDS-PAGE followed by transfer to nitrocellulose membrane, and then immunoblotting was conducted using total Akt (#9272, Cell Signaling), phospho-Akt (Ser473) (#9271, Cell Signaling), total eIF2α (#9722, Cell Signaling), phospho-eIF2α (Ser51) (#9721, Cell Signaling), total Ampk (#2532, Cell Signaling), phospho-Ampk (Thr172) (#2531, Cell Signaling), total p62 (PM045, MBL), and phospho-p62 (PM074, MBL). A Peroxidase (HRP)-conjugated anti-rabbit antibody (NA9340V) was used as a secondary antibody. Immunodetection was performed using Immobilon Western Chemiluminescent HRP Substrate (Millipore) and signals were detected by a luminoimage analyzer (Fusion System Solo 7S; M&S Instruments Inc). Quantification and brightness adjustment were conducted with the Fiji software (ImageJ; National Institutes of Health). For the images with high background, we used the subtract background function in the Fiji software (*106*). Conversion of the images to 8-bit image was conducted by Photoshop CS6 (Adobe).

### Identification of starvation-responsive molecules

We analyzed molecules or phosphorylation events that were quantified in more than half of replicates in both WT and *ob*/*ob* mice at all the time points after food removal (Fig. 2 and Fig. S1). For every molecule, one-way-ANOVA tests were conducted in data for WT or *ob*/*ob* mice. Molecules with *P* < 0.025 were defined as starvation-responsive molecules. False discovery rate (FDR) of the responsive molecules was less than 0.1 except for phosphoproteome data (Table 1). FDR was calculated using the Storey’s procedure (*107*) except for the lipidomic and FFA & acyls layers, including molecules in the lipid and FFAs & acyls layers. For lipidomic and FFA & acyls analyses, Benjamini-Hochberg procedure was conducted (*108*), because the number of measured molecules were small in these omic layers (16 for lipidome and 41 for FFA & acyls).

For transcriptomic analysis, the likelihood ratio test in the edgeR package (version 3.34.1) (*49*) of the R language (version 4.1.1) was used to identify significantly changed molecules during starvation. Mean time courses for each molecule in WT and *ob*/*ob* mice were log2-transformed relative to the geometric mean of the values of WT and *ob*/*ob* mice before food deprivation (0 h). Using these log2-transformed time courses, the area under the curve (AUC) was calculated as the area between the time course curves during starvation and the baseline value at 0 h (before food deprivation).

Molecules with positive AUC and *P* value < 0.025 were defined as increased molecules, and molecules with negative AUC and *P* value < 0.025 were defined as decreased molecules.

### Pathway enrichment analysis

We conducted pathway enrichment analysis for increased or decreased transcripts (Table 3; Data File S3), proteins (Table 4; Data File S3), and phosphoproteins (Data File S3). From KEGG database (*55–57*), we extracted gene sets of pathways in “Metabolism”, “Genetic Information Processing”, “Environmental Information Processing”, “Cellular Processes”, and “Organismal Systems”. Note that “Global/overview pathways” were removed. The enrichment of increased or decreased molecules in each pathway was determined using one-tailed Fisher’s exact test followed by FDR correction by the Benjamini-Hochberg procedure (*108*). Transcripts, proteins, and phosphoproteins quantified in more than half of the replicates in WT and *ob*/*ob* mice were used as background. Pathways with *Q*-values < 0.1 were identified significantly enriched.

### Inference for activities of TFs

We inferred the TF activities using transcriptomic data and ChIP-Atlas (*76*, *77*) (Table 5; Data File S4). ChIP-Atlas includes a database curating publicly available ChIP-seq experiments, that can be used for inference of the activities of TFs (*109*). The list for all the ChIP-seq experiments was downloaded (https://github.com/inutano/chip-atlas/wiki#downloads_doc) and all the experimental IDs of mm10 were extracted. We focused on the ChIP-seq experiments conducted using “liver” or “hepatocytes” in mice. Using this list of experimental IDs of mm10, we scraped “Target Genes” information that includes genes whose transcription start sites (TSSs) were identified as binding partners with the bait proteins (TFs) (https://chip-atlas.dbcls.jp/data/mm10/target/Protein.Distance.tsv). The “Distance” was set as 1 kb from TSS. We set the threshold for statistical significance of values calculated by peak-caller MACS2 as 50 and identified target genes for each ChIP-seq experiment. To keep quality of the inference, we removed ChIP-seq experiments that detected less than 50 target genes. Through these processes, we obtained a network between bait proteins (TFs) for each ChIP-seq experiment and “Target Genes” with TSSs that were bound to the bait proteins (TFs). Note that this network is a network of ChIP-seq experiments and transcripts, not TFs and transcripts. In other words, ChIP-seq experiments for the identical TFs were not combined at this step.

Using this network, we conducted TF enrichment analyses for increased or decreased transcripts during starvation in WT and *ob*/*ob* mice. The enrichment of increased or decreased transcripts in each ChIP-seq experiment was determined using one-tailed Fisher’s exact test followed by FDR correction conducted by the Benjamini-Hochberg procedure (*108*). Transcripts quantified in more than half of the replicates in both WT and *ob*/*ob* mice were used as background. ChIP-seq experiments with *Q*-values less than 0.1 were identified as significantly enriched ChIP-seq experiments. TFs for which significant enrichment was observed in at least one ChIP-seq experiment were identified as candidate TFs that changed their activities during starvation.

We further screened the candidate TFs to define the up-regulated or down-regulated TFs using transcriptomic, proteomic, and phosphoproteomic data. It should be noted that TFs that are registered as negative regulators in Transcriptional Repressor Complex (GO_0017053) were defined as negative regulators, and other TFs were regarded as activators in this study. If a transcriptional activator was enriched in increased transcripts, and any of the transcriptomic, proteomic, or phosphoproteomic data of the TF increased and none of the transcriptomic nor proteomic data decreased, the TF was identified as an up-regulated TF. If a transcriptional activator was enriched in decreased transcripts, and any of the transcriptomic, proteomic, or phospho-proteomic data of the TF decreased and none of the transcriptomic nor proteomic data increased, the TF was identified as a down-regulated TF. If a transcriptional inhibitor was enriched in increased transcripts, and any of the transcriptomic, proteomic, or phosphoproteomic data of the TF decreased and none of the transcriptomic nor proteomic data increased, the TF was identified as an up-regulated TF. If a transcriptional inhibitor was enriched in decreased transcripts, and any of the transcriptomic, proteomic, or phosphoproteomic data of the TF increased and none of the transcriptomic nor proteomic data decreased, the TF was identified as a down-regulated TF.

Downstream genes of the inferred TFs were defined based on the gene regulatory network from the ChIP-Atlas database. For each up-regulated TF, increased transcripts whose TSS bind to the TF in at least one ChIP-experiment were defined as downstream transcripts of the TF. For each down-regulated TF, decreased transcripts whose TSS bind to the TF in at least one ChIP-experiment were defined as downstream transcripts of the TF.

### Inference for activities of protein kinases

We inferred the protein kinase activities using phosphoproteomic data and NetPhorest (*110*) (Table 6; Data File S4). We constructed a kinase regulatory network using NetPhorest (*110*), a protein sequence-based algorithm to infer kinase-substrate relationships. The inputs for NetPhorest were mouse protein sequences that were associated with ENSMUSP in FASTA format obtained from Mus_musculus.GRCm38.97.pep.all.fa in Ensembl database (*96*, *97*), and a list for phosphorylated residues measured in our phosphoproteomic data. The sequences in FASTA format were handled using Seqkit tool (*111*). With the NetPhorest algorithm, we obtained posterior probabilities that a residue was phosphorylated by protein kinase groups. We selected protein kinase groups with posterior probability value more than 0.1 for each residue. Protein kinases contained in protein kinase groups were obtained from NetworKIN (*112*). We converted the NetworKIN kinase names to Ensembl IDs using Ensembl databases, STRING data in NetWorKIN_release3.0, and manual curation (Data File S10).

We obtained a regulatory network among transcripts, proteins, phosphorylation, and activities of protein kinases and phosphorylation of substrate proteins. Using this network, we conducted kinase enrichment analyses for increased or decreased phosphorylation events during starvation in WT and *ob*/*ob* mice. The enrichment of increased or decreased phosphorylation events in each kinase group was determined using one-tailed Fisher’s exact test followed by FDR correction conducted by the Benjamini-Hochberg procedure (*108*). Phosphorylation events quantified in more than half of the replicates in WT and *ob*/*ob* mice were used as background. Protein kinase groups with *Q*-values less than 0.1 were identified significantly enriched protein kinases.

We screened the candidate protein kinases to define the upregulated or downregulated protein kinases using transcriptomic, proteomic, and phosphoproteomic data. If a protein kinase was enriched in increased phosphorylation events, and any of the transcriptomic, proteomic, or phosphoproteomic data of the protein kinase increased and none of the transcriptomic nor proteomic data decreased, the protein kinase was identified as an up-regulated protein kinase. If a protein kinase was enriched in decreased phosphorylation events, and any of the transcriptomic, proteomic, or phospho-proteomic data of the protein kinase decreased and none of the transcriptomic, proteomic, nor phosphoproteomic data increased, the protein kinase was identified as a downregulated protein kinase.

### Inference for activities of signaling pathways

Pathways in KEGG (*55–57*) database that contain “signaling” in their pathway names were defined as signaling pathways. We identified up-regulated or down-regulated signaling pathways that were enriched in increased or decreased transcripts, proteins, or phosphorylation events at least one omics layer of transcriptomic, proteomic, or phosphoproteomic data (Table 7; Data File S3).

### Construction of allosteric regulatory network

Allosteric regulation of metabolic reactions by metabolites was determined using BRENDA database (https://www.brenda-enzymes.org/download.php) (*113*). Activating and inhibiting compounds in mammals were extracted from brenda_2023_1.txt. Note that the taxonomy list of mammals was obtained from NCBI (https://ftp.ncbi.nlm.nih.gov/pub/taxonomy/new_taxdump/) (*114*). Compound names in brenda_2023_1.txt were transformed to KEGG ID via InChI key or using compound names in KEGG (ftp.kegg.net/kegg/ligand/compound.tar.gz) (*55–57*) or HMDB (https://hmdb.ca/downloads, ver. 5) (*115*). InChI keys of BRENDA ligands were obtained from the search results of “Search Ligands” function in the classic view of BRENDA without any query terms. Through these processes, we obtained the table of the pairs of EC numbers and KEGG compound IDs.

### Construction of transport network

We used the Transporter Classification Database (TCDB, https://tcdb.org/) (*58*) to identify transporters and substrates. Transport_activity was defined by the substrates of transports. To define substrates of transports, TC2ChEBI.txt in TCDB was processed as follows. TC2ChEBI.txt includes ChEBI IDs, that is a hierarchical database and linked to each other with “is_a” function (for example, anion (CHEBI: 22563) **is a** ion (CHEBI: 24870)) (*116*). To focus on specific substrates, we selected ChEBI IDs that include less than 50 ChEBI IDs that can be traced by “is_a” function. The selected ChEBI IDs were transformed to KEGG compound IDs using KEGG (ftp.kegg.net/kegg/ligand/compound.tar.gz, downloaded on 13rd/12/2022) (*55–57*) or HMDB (https://hmdb.ca/downloads, ver. 5) (*115*), directly or via compound names.

For transporter protein, we selected transporters that are expressed in human or mice using Uniprot database (ftp.uniprot.org/pub/databases/uniprot/knowledgebase/complete/) (*117*), using taxonomy list of mammals in NCBI (ftp.ncbi.nlm.nih.gov/pub/taxonomy/new_taxdump/) (*114*). Among them, we further screened the transporters localizing on cell membrane, by selecting transporters that include “Apical cell membrane”, “Apicolateral cell membrane”, “Basal cell membrane”, “Basolateral cell membrane”, “Cell membrane”, or “Lateral cell membrane” in SUBCELLULARLOCATION in Uniprot database with at least one evidence in ECO:0000250, ECO:0000269, ECO:0000303, or ECO:0000305. Uniprot accessions were transformed to Entrez IDs by Uniprot database. Entrez IDs for human were translated to Entrez IDs for mice using gene ortholog table in NCBI database (gene_orthologs, ftp.ncbi.nlm.nih.gov/gene/DATA/) (*114*).

Combining information of substrates and transporter proteins, we obtained a table of combinations of TCDB IDs, transporter Entrez IDs and KEGG compound IDs. Note that the transport_activities were defined by the name of substrates of the transport.

### Construction of the global starvation-responsive transomics network

To construct the global starvation-responsive transomics network, nodes were grouped into layers based on the chemical properties, biological functions, or both (Table 8).

Layers were defined as follows:

- The signaling_activity layer includes signaling pathways of KEGG database (*55–57*), that contains “signaling” in their pathway names.
- The TF_activity, TF_mRNA, TF_protein, and TF_PTM layers include activities, transcripts, proteins, and phosphorylations of all the TFs investigated, those were included in “liver” or “hepatocytes” data in ChIP-Atlas database (*76*, *77*).
- The kinase_activity layer contains all the activities of the protein kinase groups in the NetPhorest database (*110*).
- The kinase_mRNA, kinase_protein, and kinase_PTM layers include transcripts, proteins, and phosphorylations of the protein kinases that belong to the kinase groups in the NetPhorest database, respectively.
- Metabolic enzymes were defined using KEGG Pathway information (ftp.kegg.net\kegg\pathway\organisms\mmu.tar.gz and ftp.kegg.net\kegg\pathway\pathway.list). The enzymes that belong to the pathways that are classified in “Metabolism” were defined as metabolic enzymes, except for Slc33a1 (mmu:11416), a transporter.
- The enzyme_mRNA, enzyme_protein, and enzyme_PTM layers include transcripts, proteins, and phosphorylations of metabolic enzymes, respectively.
- The metabolite (in the liver), lipid, FFA & acyls, and plasma metabolite layers include all the compounds measured in our metabolomic, lipidomic, and the FFA & acyls measurements.
- The transport_activity layer includes all the possible transport activities, that is defined using substrates of the transports in TCDB.
- The transporter_mRNA, transporter_protein, and transporter_PTM layers include all the measured transcripts, proteins, and phosphorylations of transporters regulating transport_activities, respectively.
- The metabolic reaction layer includes EC numbers in the KEGG database (*55*– *57*) (ftp.kegg.net\kegg\genes\organisms\mmu\mmu_link.tar.gz) that metabolic enzymes belong to.

Regulatory connections between nodes were determined using databases. The following databases or assumptions were used to infer the edges (regulatory connections):

- KEGG database (ftp.kegg.net\kegg\ligand\enzyme.tar.gz) (*55–57*) was used to infer the substrate and product regulation from metabolites, lipids, and FFA & acyls to metabolic reactions.
- The BRENDA (*113*) database was used to infer the allosteric regulatory connections from metabolites, lipids, and FFA & acyls to metabolic reactions.
- Regulation from enzyme_protein and enzyme_PTM to metabolic reactions were defined using KEGG database (*55–57*).
- Regulation of transport_activities was assumed to be mediated by the plasma metabolites, transporter_proteins, and transporter_PTMs.
- TCDB was used to determine regulation of metabolites in the liver by transport_activity (*58*).
- To determine regulatory connections between the TF_mRNA, TF_protein, TF_PTM and TF_activity layers, gene symbols of the bait in ChIP-Atlas database (TFs) were translated to ENSMUSG IDs using Ensembl database (ver. 97) (*96*, *97*).
- ChIP-Atlas (*76*, *77*) was used to determine regulatory connections between the TF_activity, and enzyme_mRNA or transporter_mRNA layers.
- Regulation between the enzyme_mRNA and enzyme_protein, and transporter_mRNA and transporter_protein, were established by aligning Entrez IDs (*114*) of the metabolic enzymes and transporters.
- For protein kinase and phosphorylations, the kinase_mRNA, kinase_protein, and kinase_PTM layers were assumed to regulate the kinase_activity layer.
- NetPhorest (*110*) was used to determine regulation between the kinase_activity layer and the enzyme_PTM layer.
- Signaling pathways were assumed to regulate TF_mRNA, TF_protein, TF_PTM, kinase_mRNA, kinase_protein, kinase_PTM, enzyme_mRNA, enzyme_protein, enzyme_PTM, transporter_mRNA, transporter_protein, and transporter_PTM. The regulatory relationships were defined using KEGG database. Note that signaling pathways and downstream molecules were connected regardless of increase or decrease of the molecules.

For phosphorylation-related regulation, phosphorylation events curated as “activity, inhibited”, “enzymatic activity, inhibited”, “protein degradation”, “receptor desensitization, induced”, “receptor inactivation, induced”, “receptor internalization, induced”, or “receptor recycling, inhibited” in PhosphositePlus (https://www.phosphosite.org/staticDownloads) (*118*) were assumed to negatively regulates their downstream nodes.

### Construction and structural analysis of the starvation-responsive metabolic network

The starvation-responsive metabolic network is a sub-network of the global starvation-responsive metabolic network containing metabolic reactions and their direct regulators, including metabolite, lipid, FFA & acyls, enzyme_protein and enzyme_PTM layers (Data File S6). It should be noted that if a metabolite regulates a metabolic reaction both as a substrate or product, and as an allosteric regulator, the edges were counted as a single edge in the starvation-responsive metabolic network. The network was regarded as directed graph for the calculation of degree and degree distribution. The degree of each node (*k*) was counted by the centrality function in MATLAB, with the ‘outdegree’ option specified, and degree distribution (*n*(*k*)) was determined. Scaling parameter was inferred by fitlm function in MATLAB. Hub nodes were defined as responsive nodes within the top 2% based on highest degrees.

The network was regarded as an undirected graph for the calculation of the following structural properties. Because the starvation-responsive metabolic network is composed of metabolic reactions and their direct regulators, all the edges are directed to metabolic reactions and no edge starts from metabolic reactions. In this case, for the calculation of the cluster sizes, only weakly connected components are generated. For simplicity, we regarded the starvation-responsive metabolic network as an undirected graph. In addition, the average path length would be 1 if we regard the starvation-responsive metabolic network as a directed graph, because the regulators of the metabolic reactions (metabolites, lipids, FFA & acyls, proteins and phosphorylation events) regulate metabolic reactions with a single edge, and no edge starts from the metabolic reactions.

The conncomp function in MATLAB was used to determine the maximum cluster size. Distances between nodes were calculated by the distances function in MATLAB and the average path length of the graphs were determined. Note that distances among nodes without any connection were removed for calculation of the average path length.

To connect hub nodes to metabolic pathways (fig. S4), metabolic reactions were classified into metabolic pathways in carbohydrate metabolism, lipid metabolism, nucleotide metabolism, and amino acid metabolism using KEGG database (*55–57*).

Then the number of metabolic reactions in each pathway that was regulated by the hub nodes was counted. The number of metabolic reactions were log2-transformed to display in the heatmap. Note that some metabolic reactions overlap among pathways because KEGG database (*55–57*) allow overlaps of metabolic reactions among pathways.

### Calculation of *t*-half and temporal order analysis

The index of the temporal rate of response (*t*-half) was defined as the time when the responses to starvation reached the half of the maximum absolute values of the gain of the responses. We defined *t*-half for the responsive nodes not for metabolic reactions.

Calculated *t*-half values were analyzed by the following procedures. The correlation between *t*-half and log10-transformed degree was determined by Spearman’s correlation coefficient. Comparison between *t*-half values of WT-specific and *ob*/*ob*-specific hub molecules were conducted by Wilcoxon rank sum test. The classification of starvation-responsive metabolites, FFA & acyls, and lipids into metabolic pathways was conducted using KEGG database (*55–57*) for comparison of *t*-half values among metabolic pathways. Note that some molecules are included in multiple metabolic pathways. Comparisons of *t*-half values among metabolic pathways and omic layers were conducted using Kruskal-Wallis test followed by Dunn-Sidak test. Comparisons of *t*-half values between WT and *ob*/*ob* mice were conducted by Wilcoxon rank sum test followed by FDR correction conducted by the Benjamini-Hochberg procedure (*108*).

To identify the essential molecules for negative correlation between log10-transformed degree and *t*-half in the starvation-responsive metabolic network in WT mice, leave-one-out correlation and cumulative leave-one-out correlation were calculated for each molecule. The leave-one-out correlation of molecule X was defined as correlation coefficient between the log10-transformed degree and *t*-half for all the responsive molecules in the starvation-responsive metabolic network except for X. The cumulative leave-one-out correlation of molecule X was defined as correlation coefficient between the log10-transformed degree and *t*-half of the responsive molecules in the starvation-responsive metabolic network with leave-one-out correlations that were smaller (the contribution to the negative correlation between log10-transformed degree and *t*-half is smaller) than X.

### Analysis of co-regulation among molecules in the starvation-responsive metabolic network and construction of correlation metabolic network

Pairwise Pearson’s correlation coefficients were calculated among time courses of significantly changed or all the molecules [metabolites, lipids, FFA & acyls, protein (enzyme_protein), and phosphorylation events (enzyme_PTM)] in the starvation-responsive metabolic network in WT and *ob*/*ob* mice. The same procedure was conducted for significantly changed molecules in each metabolic pathway and the correlation coefficients between WT and *ob*/*ob* mice were compared by Wilcoxon rank sum test followed by FDR correction conducted by the Benjamini-Hochberg procedure (*108*).

Based on these correlations, pairs of molecules whose Pearson’s correlation coefficients are more than the threshold were connected by edges, resulting in correlation networks of the responsive molecules in the starvation-responsive metabolic network in WT and *ob*/*ob* mice, named as correlation metabolic networks. We tested several thresholds for the absolute value of the Pearson’s correlation coefficients (from 0.3 to 0.9). For each tested threshold, degree of each node was calculated using centrality function in MATLAB with the ‘degree’ option specified, and then degree distribution (*n*(*k*)) was determined. The scaling parameter was inferred by fitlm function in MATLAB for each threshold. In addition, the number of the clusters (fig. S12**B**), the relative size of the large cluster (fig. S12**C**) and the average path length (fig. S12**D**) were calculated using MATLAB (conncomp function for the number of the clusters and the relative size of the largest cluster and distances function for the average path length).

### Functional analysis of transcriptomic and proteomic data for potential clinical application

Hallmark gene sets (h.all.v7.4.entrez.gmt) were downloaded from the download page of MSigDB (*72*) (http://www.gsea-msigdb.org/gsea/login.jsp). Gene IDs were converted using gene ortholog table in NCBI database (gene_orthologs, https://ftp.ncbi.nlm.nih.gov/gene/DATA/) (*114*). We conducted functional enrichment analysis for significantly changed transcripts and proteins during starvation. The enrichment of significantly changed molecules in each hallmark was determined using one-tailed Fisher’s exact test followed by FDR correction by the Benjamini-Hochberg procedure (*108*). Transcripts and proteins quantified in more than half of the replicates in WT and *ob*/*ob* mice were used as background. Hallmarks with *Q*-values < 0.1 were identified significantly enriched.

### Construction and analysis of each metabolic pathway

For fatty acid degradation (fig. S7), map of “Fatty acid degradation” (mmu00071) was extracted from KEGG database (*55–57*) and measured fatty acids, acyl-carnitines, and acyl-CoAs were basically annotated to palmitic acid (C00249), palmitoyl-CoA (C00154), and L-palmitoylcarnitine (C02990), respectively. Long-chain fatty acids and acyl-carnitines, whose number of carbons were more than 20, were annotated to long-chain fatty acid (C00638) and long-chain acyl-[acyl-carrier protein] (C20683). In addition, “Propanoate metabolism” (mmu00640) and “Butanoate metabolism” (mmu00650) were merged for acyl-CoA (3:0) and ketone body metabolism, respectively. Major metabolic reactions, substrates and products were manually extracted for further analyses.

For glycolysis/gluconeogenesis (fig. S9) and urea cycle (fig. S10), major metabolic reactions, substrates and products were extracted from the map of “Glycolysis/gluconeogenesis” (mmu00010) and “Arginine biosynthesis” (mmu00220), respectively. Allosteric regulators (metabolites, FFA & acyls, and lipids), enzyme_protein, and enzyme_PTM of the extracted metabolic reactions were obtained from the global starvation-responsive transomic network (Fig. 3A; Data File S5) and visualized. For fatty acid degradation, significantly responsive enzyme_mRNA in WT or *ob*/*ob* mice corresponding to the responsive enzyme_protein in WT or *ob*/*ob* mice were also extracted and visualized. The starvation-responsive metabolic network of the extracted molecules was obtained from the starvation-responsive metabolic network (Data File S6). For glycolysis/gluconeogenesis (fig. S9) and urea cycle (fig. S10), the degree of each node (*k*) was counted by the centrality function in MATLAB, with the ‘outdegree’ option specified. The degree distribution (*n*(*k*)) was also determined. Note that the networks were regarded as direct graphs for the calculation of degree and degree distribution. The fitlm function in MATLAB was used to calculate the scaling parameters. Correlations between *t*-half values and log10-transformed degree were determined by Spearman’s correlation coefficient. Pairwise Pearson’s correlation coefficients among time courses of the extracted molecules in each metabolic pathway were calculated to examine the co-regulations among the molecules in each metabolic pathway.

For fatty acid degradation, *t*-half values of WT and *ob*/*ob* mice in each molecular type (FFA, acyl-carnitine, acyl-CoA, ketone body, metabolite, enzyme_protein, and enzyme_PTM) were compared by Wilcoxon rank sum test followed by FDR correction conducted by the Benjamini-Hochberg procedure (*108*).

For fatty acid degradation, correlation between the time courses of enzyme_mRNA and enzyme_protein in each molecule in each genotype, that is included the extracted molecules visualized (fig. S7), were calculated using Pearson’s correlation, followed by a comparison between WT and *ob*/*ob* mice by *t*-test.

### Implementation

Statistical test, enrichment analysis, construction of transomic network, and network analysis were conducted using MATLAB 2021a (The MathWorks Inc.). For edgeR, R language (version 4.1.1) was used. For handling of database, MATLAB, Python 3.8, R language, and Cygwin (version 3.2.0-1) were used. Transcriptomic analysis obtaining count and TPM data from sequences and running of NetPhorest software were conducted on the supercomputer system from the National Institute of Genetics in Japan. Transomic networks were visualized in Graph Modeling Language formats using Python 2.7 and Visualization and Analysis of Networks containing Experimental Data (VANTED) (*119*).

### Statistical Analysis

As described above, we conducted statistical analysis using MATLAB 2021a (The MathWorks Inc.) or R. For identification of responsive molecules, one-way ANOVA tests (ANOVA-like testing of edgeR (51–53) for transcriptome) were conducted in WT and *ob*/*ob* mice respectively. For pathway enrichment analysis, inference of activities of TFs, and inference of activities of protein kinases, one-tailed Fisher’s exact test followed by FDR correction by the Benjamini-Hochberg procedure (96) was conducted. For comparison of *t*-half values of the hub molecules between WT and *ob*/*ob* mice, Wilcoxon rank sum test was conducted. For multi comparison of *t*-half values, Kruskal-Wallis test followed by Dunn-Sidak test was conducted. For identification of responsive molecules or phosphorylation events, *P*-values less than 0.025 were defined as significant. For multiple comparisons except for identification of responsive molecules or phosphorylation events, *Q*-values less than 0.1 was defined as significant. *P*-values less than 0.05 were defined as significant in other cases.

## Acknowledgments

We thank Maki Ohishi and Ayano Ueno (Keio University) for their expertise and assistance with metabolome analysis using CE-MS. We thank Maiko Goto (Kyushu University) for technical assistance with lipidome analysis using SFC-MS/MS and LC-MS. We also thank Kazusa DNA Research Institute for conducting the proteomic and phosphoproteomic measurements. We also thank our laboratory members for critically reading this manuscript and technical assistance with the experiments. The computational analysis of this work was performed in part with support of the supercomputer system of the National Institute of Genetics (NIG), Research Organization of Information and Systems (ROIS). The authors thank Nancy R. Gough (BioSerendipity, LLC) for editorial assistance and critical input. We used ChatGPT, DeepL, and Claude3 for the assistance of manuscript editing and coding.

## Funding and Assistance

This study was supported by the Japan Society for the Promotion of Science (JSPS) KAKENHI grant numbers JP17H06299, JP17H06300, JP18H03979, JP21H04759, JP23H04939, and JP23H04946 to S.K.); CREST, the Japan Science and Technology Agency (JST) (JPMJCR2123 to S.K., Y. Inaba, and T.S.); The Uehara Memorial Foundation (to S.K.); K.M. receives funding from a Grant-in-Aid for Early-Career Scientists (JP21K15342). A.Hatano receives funding from a Grant-in-Aid for Early-Career Scientists (JP22K15034). T.K. receives funding from a Grant-in-Aid for Early-Career Scientists (JP21K16349). T.T. receives funding from a Grant-in-Aid for Early-Career Scientists (JP20K19915). H.O. receives funding from a Grant-in-Aid for Early-Career Scientists (JP22K17992). S.O. receives funding from a Grant-in-Aid for Early-Career Scientists (JP21K14467). Y. Inaba also receives AMED-PRIME (JP23gm6910002). This work was also supported by the Japan Society for the Promotion of Science (JSPS) KAKENHI grant number JP22H04925 (PAGS); in part by the MEXT Cooperative Research Project Program, Medical Research Center Initiative for High Depth Omics, and CURE:JPMXP1323015486 for MIB, Kyushu University; and AMED Grant Number JP21zf0127001 (T.S.), JST, MEXT KAKENHI Grant Number JP23H04946(T.S.) and World Premier International Research Center Initiative (WPI), Human Biology-Microbiome-Quantum Research Center (Bio2Q) (T.S.), MEXT, Japan.; and JST FOREST Program (Grant Number JPMJFR2052, to A. Hirayama).

## Author contributions

K.M., A. Hatano, T.K., H.S, S.O, and S.K. designed the project. K.M., A. Hatano, and M.M. conducted animal experiments and sample preparation for omic measurements. K.M. conducted single-omic analysis, transomic analysis, and network structural and temporal analyses. T.S. and A. Hirayama performed metabolomic analysis using CE-MS and IC-QEMS. Y. Izumi., M.T., and T.B. performed lipidomic analysis using LC-MS and SFC-MS/MS. Y.S. performed transcriptomic analysis using RNA-seq. K.M., T.T., R.E., and H.O. analyzed the RNA-seq data. K.M., T.K., H.S., D.L., and A.T. conducted database for transomic analysis. The writing group consisted of K.M., Y. Inaba, H.I., and S.K. The study was conceived and supervised by K.M. and S.K.

## Conflict of Interest

The authors declare that they have no competing interests.

## Data and code availability

Sequencing data have been deposited in the DNA Data Bank of Japan Sequence Read Archive (DRA) (www.ddbj.nig.ac.jp/) with accession no. DRR571760-DRR571839. Proteome and phosphoproteome data were deposited in jPOSTrepo (*120*) (https://repository.jpostdb.org/) with accession no. JPST003082 (proteome, for reviewers: https://repository.jpostdb.org/preview/56792610366738a30bc36f, Access key is 9357) and JPST003130 (phosphoproteome, for reviewers: https://repository.jpostdb.org/preview/124893828466738a2ae0323, Access key is 4346). All other data are present in the paper or the Supplementary Materials. The code used for the analysis in this paper was deposited in Zenodo (DOI: 10.5281/zenodo.14061381).

## Prior Presentation

A non–peer-reviewed version of this article was submitted to the bioRxiv preprint server (https://www.biorxiv.org/content/10.1101/2024.06.17.599249v1).

## Supplementary Materials

Figs. S1 to S12

Data Files S1 to S10

## SUPPLEMENTARY FIGURE TITLES AND LEGENDS

**fig. S1:** Time courses of responsive molecules during starvation The x-axis represents the starvation time (hours) and the y-axis represents the concentration of the indicated molecule: ng/µg-protein for glycogen, nmol/mg-protein for other metabolites (F6P, Ala, 3-hydroxybutyrate); pmol/mg-protein for lipids (PS, PE, PI) and FFA & acyls (Acyl-CoA 16:0); arbitrary units for proteins (Cpt1a, Pck1, G6pc) and phosphorylation (Ser^775^ of Pfkl). TPM for transcripts (*Cpt1a*, *Pck1*, *G6pc*). Blue lines are the responses to starvation in WT mice, and red lines are those in *ob*/*ob* mice. Solid lines represent that the responses were significant; dotted lines represent no significance. Data are shown as the mean and SEM. Dots represent the data from individual mice. *n* = 5 biological replicates per group. *P* values of one-way ANOVA of the molecules or ratio with significant changes are shown.

**fig. S2:** Overview of each omic data The heatmaps of the time courses of 123 responsive metabolites (liver, **A**), 10 responsive lipids (**B**), 27 responsive FFA & acyls (**C**), 60 responsive metabolites (plasma, **D**), 9,872 responsive transcripts (**E**), 663 responsive proteins (**F**), and 391 responsive phosphorylation events (**G**) of WT and *ob*/*ob* mice during starvation. For heatmap, two time courses for each molecule were divided by the geometric mean of the values of WT and *ob*/*ob* mice before starvation (0 h) and then log2-transformed. The x-axis represents time for starvation, and the y-axis represents the molecules or phosphorylation events. *n* = 5 biological replicates per group.

**fig. S3:** Regulatory relationships in the starvation-responsive transomic network We assumed that the starvation-responsive molecules cooperatively regulate metabolic reactions (see also Fig. 1D). (**A**) Metabolic network. The enzyme_protein (protein abundance of metabolic enzymes) and enzyme_PTM (phosphorylation events on metabolic enzymes) regulate metabolic reactions (identified with KEGG). Metabolites, lipids, and FFA & acyls regulate metabolic reactions as substrates, products (identified with KEGG), and allosteric regulators (identified with BRENDA). (**B**) Transport network identified with TCDB. The transporter_protein (protein abundance of transporters), transporter_PTM (phosphorylation events on transporters), and plasma metabolites regulate transport_activity (activities of transport). Transport_activity regulates metabolites in the liver, thereby regulating metabolic reactions in the liver. (**C**) Gene regulatory network identified with ChIP-Atlas and Ensembl. The TF_mRNA (mRNA abundance of TFs), TF_protein (protein abundance of TFs), and TF_PTM (phosphorylation events on TFs) regulate TF_activity (activities of TFs). TF_activity regulates enzyme_mRNA (mRNA abundance of metabolic enzymes) and transporter_mRNA (mRNA abundance of transporters). Enzyme_mRNA and transporter_mRNA regulate enzyme_protein and transporter_protein, respectively, and were aligned with Ensembl database. (**D**) Phosphorylation network identified with NetPhorest. The kinase_mRNA (mRNA abundance of kinases), kinase_protein (protein abundance of kinase), and kinase_PTM (phosphorylation events on kinases) regulate kinase_activity (activities of kinases). The kinase_activity regulates enzyme_PTM and transporter_PTM. (**E**) The number of metabolic reactions that lipids regulate as substrates, products, and/or allosteric regulators. The colors of the bars represent common responses (green), *ob*/*ob*-specific responses (red), and opposite responses (magenta) to starvation.

**fig. S4:** Metabolic pathways regulated by the hub nodes Heatmaps representing the number of regulated metabolic reactions in each metabolic pathway (x-axis) regulated by the hub nodes (y-axis) in the starvation-responsive metabolic network in WT (**A**) and *ob*/*ob* mice (**B**). Colors of the labels of the hub nodes represent the responsiveness: green indicates common responses both in WT and *ob*/*ob* mice; blue, specific to WT mice; red, specific to *ob*/*ob* mice; magenta, opposite responses between WT and *ob*/*ob* mice. KEGG metabolic pathways (x-axis) in carbohydrate metabolism (olive green), lipid metabolism (orange), nucleotide metabolism (brown), and amino acid metabolism (light blue) are displayed.

**fig. S5:** Maintenance of the number of nodes with relatively large degrees in *ob*/*ob* mice The number of nodes (**A**-**E**) and the percentage of nodes to the number of responsive nodes in each genotype (**F**-**J**), whose log10-transformed degree are as indicated. (**K**) The percentages of nodes whose log10-transformed degree is more than 1.

**fig. S6:** Identification of molecules essential for the negative correlation between degree and *t*-half in WT mice (**A**) Schema for leave-one-out correlations. Log10-transformed degree negatively correlated with *t*-half in WT mice (center, “Rho of full”). The leave-one-out correlation of molecule X (left, “Rho without X”) was defined as correlation coefficient between log10-transformed degree and *t*-half that was calculated for all the molecules except for X. The larger leave-one-out correlation (left, “Rho without X”) represents the larger contribution to the negative correlation between log10-transformed degree and *t*-half. By contrast, the smaller leave-one-out correlation (right, “Rho without Y”) represents the smaller contribution to the negative correlation between log10-transformed degree and *t*-half. (**B**) The cumulative leave-one-out correlation of molecule X (right) was defined by the correlation coefficient between log10-transformed degree and *t*-half that was calculated for the molecules with leave-one-out correlation smaller than X (indicated in light blue). (**C**) Leave-one-out correlations (gray bars), cumulative leave-one-out correlations (black dots), and *P*-values of the cumulative leave-one-out correlations (black solid line) of the indicated molecules. Here, we displayed molecules with *P*-values of the cumulative leave-one-out correlations that were less than 0.05 and with leave-one-out correlations that were less than other molecules. *n* = 291 molecules in total.

**fig. S7:** The starvation-responsive metabolic network for starvation-responsive metabolic reactions in fatty acid degradation (**A**) Time courses of fatty acids as substrates, acyl-CoAs and acyl-carnitines as intermediates, and 3-Hydroxybutyrate (a ketone body) as a product of fatty acid degradation. Gray lines indicate the time courses of molecules in each group in each genotype. Solid lines represent significantly responsive molecules, while dashed lines represent not significantly responsive molecules. Bold and colored lines represent mean and STD of the time courses of the significantly responsive molecules. The total numbers of molecules are shown in parentheses following to the names of the molecule types. The numbers of significantly changed molecules in each genotype are shown in parentheses following to the label for genotype. (**B**) The transomic network for starvation-responsive metabolic reactions in fatty acid degradation, that were manually constructed based on “Fatty acid degradation” (mmu00071), “Propanoate metabolism” (mmu00640), and “Butanoate metabolism” (mmu00650) in the KEGG database (*55*– *57*). Time courses of measured metabolites, lipids, FFA & acyls, enzyme_protein, and enzyme_PTM are shown for corresponding nodes as the mean and SEM. *n* = 5 biological replicates per group. The colors of the frames represent common responses (green), WT-specific responses (blue), *ob*/*ob*-specific responses (red), and opposite responses (magenta) to starvation. The black frames indicate that were not responsive to starvation. From metabolites, lipids, and FFA & acyls to metabolic reactions, only allosteric regulatory connections are colored. (**C**) The *t*-half values of the indicated types of molecules in WT and *ob*/*ob* mice. The x-axis represents molecule types and genotypes. The y-axis represents *t*-half values. Light blue dots represent *t*-half values of each molecule. Red horizontal lines indicate median, boxes the interquartile range (25th to 75th percentile), and whiskers 1.5 times the interquartile range. Red crosses represent outliers. *Q*-value less than 0.1 for the differences of *t*-half values between WT and *ob*/*ob* mice by Wilcoxon rank sum test followed by FDR using Benjamini-Hochberg procedure is shown. The number of molecules in each category is shown in parentheses following to the genotype label. (**D**) The number of responsive molecules of the indicated molecule types in WT and *ob*/*ob* mice. Blue represents WT and red represents *ob*/*ob* mice. The x-axis represents molecule types and genotypes. The y-axis represents the number of responsive molecules. (**E**) The distribution of Pearson’s correlation coefficients between the time courses of enzyme_mRNA and enzyme_protein for each metabolic enzyme involved in fatty acid degradation, whose mRNA and protein were both measured in our transcriptome and proteome data. The x-axis represents genotype and the y-axis represents Pearson’s correlation coefficient. The *P*-value calculated by two-sample *t*-test comparing the correlation coefficients of WT and *ob*/*ob* mice is shown. *n* = 13 commonly measured molecules both in transcriptome and proteome that fatty acid degradation map (fig. S7**B**) includes.

**fig. S8:** Activities of protein translation and degradation in *ob*/*ob* mice (**A**) Phosphorylation ratio of eIF2α in WT and *ob*/*ob* mice before starvation. The x-axis represents genotype and the y-axis represents the ratio of phosphorylated eIF2α to total eIF2α. The *P*-value of a two-sample *t*-test comparing the phosphorylation ratios of eIF2α between WT and *ob*/*ob* mice is shown. (**B**) Images of western blotting for (**A**). (**C**) The time course of phosphorylation ratio of p62 in WT (blue line) and *ob*/*ob* mice (red line). The x-axis represents starvation time (h) and the y-axis represents ratio of phosphorylated p62 to total p62. The solid line represents that the response was significant; the dotted line represents no significance. Data are shown as the mean and SEM. Dots represent the data from individual mice. *P* value of one-way ANOVA of ratio-pp62 with significant changes is shown. (**D**) Images of western blotting for (**C**) at the indicated starvation time (h) of the indicated genotype. IC represents internal control to normalize the signals across membranes. Arrowheads indicate the total and phosphorylated p62 bands. *n* = 5 biological replicates per group.

**fig. S9:** The starvation-responsive metabolic network for starvation-responsive metabolic reactions in glycolysis/gluconeogenesis The starvation-responsive metabolic network for starvation-responsive metabolic reactions in glycolysis/gluconeogenesis, that were manually constructed based on “Glycolysis/gluconeogenesis” (mmu00010) in the KEGG database (*55–57*), in WT (**A**) and *ob*/*ob* (**B**) mice. Time courses of measured metabolites, lipids, FFA & acyls, enzyme_protein, and enzyme_PTM are shown for corresponding nodes as the mean and SEM. *n* = 5 biological replicates per group. The colors of the frames represent common responses (green), WT-specific responses (blue), *ob*/*ob*-specific responses (red), and opposite responses (magenta) to starvation. The black frames indicate that were not responsive to starvation. From metabolites, lipids, and FFA & acyls to metabolic reactions, only allosteric regulatory connections are colored. (**C** and **D**) The degree of the indicated nodes in WT (**C**) and in *ob*/*ob* (**D**) mice. Nodes with degree more than 4 are shown. The x-axis represents degree (the number of edges) and the y-axis represents nodes. It should be noted that the degrees shown in **C** and **D** are not necessarily identical to those shown in **A** and **B.** This is because if a molecule regulates a metabolic reaction both as a substrate or product, and as an allosteric regulator, the edges were counted as a single edge in **C** and **D**, but were shown separately in **A** and **B**. (**E**) Degree distributions of the nodes in the starvation-responsive metabolic network of glycolysis/gluconeogenesis in WT (blue) and *ob*/*ob* (red) mice. The x-axis represents the log10-transformed degree (*k*); the y-axis represents the log10-transformed number of nodes with degree of *k*. Lines show the linear regression of the degree distributions. The slope of the lines (γ), which is defined as a scaling parameter, and the 95% credible interval are shown. *n* = 49 molecules for WT mice and *n* = 37 molecules for *ob*/*ob* mice. (**F** and **G**) Scatter plots for the log10-transformed degree (x-axis) and *t*-half (y-axis) in WT (**F**) and *ob*/*ob* (**G**) mice. Dots represent the molecules (including phosphorylation events) in the starvation-responsive metabolic network of glycolysis/gluconeogenesis. Blue dots represent WT and red dots represent *ob*/*ob* mice. Spearman’s correlation coefficients (Rho) and *P* values are shown. *n* = 49 molecules for WT mice and *n* = 37 molecules for *ob*/*ob* mice. (**H**) Distribution of the pairwise Pearson’s correlation coefficients among time courses of responsive molecules in the starvation-responsive metabolic network of glycolysis/gluconeogenesis in WT and *ob*/*ob* mice. The x-axis represents correlation coefficients; the y-axis represents the number of pairs of molecules examined for correlation. *n* = 49 molecules for WT mice and *n* = 37 molecules for *ob*/*ob* mice.

**fig. S10:** The starvation-responsive metabolic network for starvation-responsive metabolic reactions in urea cycle The starvation-responsive metabolic network for starvation-responsive metabolic reactions in urea cycle, that were manually constructed based on “Arginine biosynthesis” (mmu00220) in the KEGG database (*55–57*) (**A**). Time courses of measured metabolites, lipids, FFA & acyls, enzyme_protein, and enzyme_PTM are shown for corresponding nodes as the mean and SEM. The colors of the frames represent common responses (green), WT-specific responses (blue), *ob*/*ob*-specific responses (red), and opposite responses (magenta) to starvation. The black frames indicate that were not responsive to starvation. From metabolites, lipids, and FFA & acyls to metabolic reactions, only allosteric regulatory connections are colored. (**B** and **C**) The degree of the indicated nodes in WT (**B**) and in *ob*/*ob* (**C**) mice. Nodes with degree more than 2 are shown. The x-axis represents degree (the number of edges) and the y-axis represents nodes. It should be noted that the degrees shown in **B** and **C** are not necessarily identical to those shown in **A.** This is because if a molecule regulates a metabolic reaction both as a substrate or product, and as an allosteric regulator, the edges were counted as a single edge in **B** and **C**, but were shown separately in **A**. (**D**) Degree distributions of the nodes in the starvation-responsive metabolic network of urea cycle in WT (blue) and *ob*/*ob* (red) mice. The x-axis represents the log10-transformed degree (*k*); the y-axis represents the log10-transformed number of nodes with degree of *k*. Lines show the linear regression of the degree distributions. The slope of the lines (γ), which is defined as a scaling parameter, and the 95% credible interval are shown. *N* = 40 molecules for WT mice and *n* = 22 molecules for *ob*/*ob* mice. (**E** and **F**) Scatter plots for the log10-transformed degree (x-axis) and *t*-half (y-axis) in WT (**E**) and *ob*/*ob* (**F**) mice. Dots represent the molecules (including phosphorylation events) of the starvation-responsive metabolic network in urea cycle. Blue dots represent WT and red dots represents *ob*/*ob* mice. Spearman’s correlation coefficients (Rho) and *P* values are shown. *n* = 40 molecules for WT mice and *n* = 22 molecules for *ob*/*ob* mice. (**G**) Distribution of the pairwise Pearson’s correlation coefficients among time courses of responsive molecules in the starvation-responsive metabolic network of urea cycle in WT and *ob*/*ob* mice. The x-axis represents correlation coefficients; the y-axis represents the number of pairs of molecules examined for correlation. *n* = 40 molecules for WT mice and *n* = 22 molecules for *ob*/*ob* mice.

**fig. S11:** Distribution of the pairwise Pearson’s correlation coefficients among time courses of all the measured metabolites, lipids, FFA & acyls, enzyme_protein, and enzyme_PTM The x-axis represents the pairwise Pearson’s correlation coefficients among time courses of all the measured metabolites, lipids, FFA & acyls, enzyme_protein and enzyme_PTM; the y-axis represents the number of pairs of molecules examined for correlation. The data of WT (**A**) and *ob*/*ob* (**B**) mice are shown. *n* = 5,866 molecules in each genotype.

**fig. S12:** The correlation metabolic network during starvation (**A**) Scatter plots showing the degree distributions of correlation metabolic network corresponding to each threshold of absolute values of correlation coefficients (|Rho|) ranging from 0.3 to 0.9. The x-axis represents log_10_(*k*), the log10-transformed degree; the y-axis represents log_10_(*n*(*k*)), the log10-transformed number of nodes with degree of *k*. Dots denote the degree distribution of the indicated networks and lines show the linear-regression of the degree distribution. The scaling parameters (γ) and the 95% credible interval are shown. Data for WT and *ob*/*ob* mice are shown. The number of clusters (**B**), relative size of the largest cluster (**C**) and the average path length (**D**) for each threshold are also shown. The lower threshold than 0.8 for the absolute value of correlation coefficients resulted in formation of a single cluster (**B**). The higher threshold (0.8 or 0.9) for the absolute value of correlation coefficients resulted in more scale-free-like networks. To identify the qualitative differences between WT and *ob*/*ob* mice clearly, we used 0.9 as a threshold for the absolute value of correlation coefficients in Fig. 7D and **E**. Using 0.9 as a threshold for the absolute value of correlation coefficients, the relative size of the largest cluster was smaller (**C**) and the average path length was larger in *ob*/*ob* mice than in WT mice (**D**). For average path length, the numbers of the investigated paths are shown in parentheses following to the genotype label. The error bars show standard deviation.

## TITLES AND LEGENDS FOR SUPPLEMENTARY TABLES

**Data File S1:** All the analyzed data This file includes all the analyzed data (metabolite, lipid, FFA & acyls, protein, phosphorylation, transcript, plasma metabolite, and western blotting), that were quantified in more than half of replicates in both WT and *ob*/*ob* mice at all the time points after food removal. Raw transcriptome data, raw proteome data, and raw phosphoproteome data are uploaded on public repositories (see Data and code availability section). All measured metabolome, lipidome, and FFA & acyls data are included in Data File S9. The columns represent biological samples (S25-S104), whose biological conditions are summarized in ‘Sample_information’ tab. The rows represent molecules or phosphorylation events. The units were nmol/mg-protein for metabolites except for glycogen, ng/μg-protein for glycogen, μM for plasma metabolites, pmol/mg-protein for lipids and FFA & acyls, TPM for transcripts, and arbitrary units for protein phosphorylation, and western blotting.

**Data File S2:** Properties of the analyzed molecules This file includes properties of the time courses (AUC and *t*-half), and results of statistical tests. The columns the names of properties of the time courses (AUC and *t*-half) and the results of statistical test (*P*-values, *Q*-values, and changes). The rows represent molecules. Abbreviations for WT_change and *ob*/*ob*-change columns were as follows: Up, Down, and NS represent increase, decrease and no significant changes.

**Data File S3:** KEGG Pathway Enrichment Analyses This file includes all the results of Fisher’s exact test in KEGG Pathway Enrichment Analyses for transcriptome, proteome, and phosphoproteome data. The Entry, Name, Class, and Subclass are displayed for the information of each pathway. *P*-values, *Q*-values, and odds ratio for Fisher’s exact test, and significantly changed downstream molecules are listed.

**Data File S4:** TF and kinase enrichment analyses This file includes all the results of TF/kinase enrichment analysis for increased or decreased transcripts/phosphorylation events in WT and *ob*/*ob* mice, respectively. For TF, Fisher’s exact tests were conducted for each ChIP-seq experiment. The table contains experimental IDs, name of TFs, organ, and cell for the information of ChIP-experiments. Also, *P*-values, *Q*-values, and odds ratio for Fisher’s exact test are displayed. For protein kinases, names of kinase groups, ENSMUSG IDs of kinases, and names of kinases were displayed. Also, *P*-values, *Q*-values, and odds ratio are listed.

**Data File S5:** The global starvation responsive transomic network This file includes all the information of nodes and edges of the global starvation-responsive transomic network. The node tab includes information of node; IDs, names, omic layers, changes in WT and *ob*/*ob* mice, and comparisons between them (common responses both in WT and *ob*/*ob* mice; specific to WT mice; specific to *ob*/*ob* mice; opposite responses between the two genotypes). The edge tab includes the layers and IDs of the source and target nodes, the type of regulation (Act: activation or Inh: inhibition), and the presence or absence of the regulation in WT and *ob*/*ob* mice (TRUE or FALSE).

**Data File S6:** The starvation responsive metabolic network This file includes all the information of nodes and edges of the starvation-responsive metabolic network. The node tab includes information of node: names, IDs, omic layers, significance of responses to starvation in WT and *ob*/*ob* mice (TRUE or FALSE), AUC of the time courses in WT and *ob*/*ob* mice, *t*-half in WT and *ob*/*ob* mice, and degrees and cluster IDs of the background network (network of the database) and starvation-responsive metabolic networks in WT and *ob*/*ob* mice. The edge tab includes the layers and IDs of the source and target nodes, the type of regulation (Act: activation or Inh: inhibition), and the presence or absence of the regulation in WT and *ob*/*ob* mice (TRUE or FALSE).

**Data File S7:** The correlation metabolic network during starvation This file includes all the pairwise correlation coefficients among time courses of molecules in the starvation-responsive metabolic networks. Each tab represents WT or *ob*/*ob* mice. Both the columns and rows represent responsive molecules in the starvation-responsive metabolic network in WT and *ob*/*ob* mice. The pairwise correlation coefficients are displayed as a matrix in each tab.

**Data File S8:** MSigDB Enrichment Analyses This file includes all the results of Fisher’s exact test in MSigDB Enrichment Analyses for transcriptome and proteome data. *P*-values, *Q*-values, and odds ratio for Fisher’s exact test, and significantly changed downstream molecules are provided for each Hallmark.

**Data File S9:** All the metabolome, lipidome, and FFA & acyls data This file includes all data of metabolome, lipidome (including FFA), acyl-carnitines, and acyl-CoAs. The columns represent biological samples (S25-S104), whose biological conditions are summarized in ‘Sample_information’ tab. The rows represent molecules. The units were nmol/mg-protein for metabolites, pmol/mg-protein for lipids including FFA, acyl-carnitine, and acyl-CoA. Most metabolites were measured by CE-MS, while F1,6P and F2,6P were separately measured by IC-QEMS. F1,6P was quantified both in CE-MS and IC-QEMS. We used values of IC-QEMS in this study.

**Data File S10:** Kinase names and ENSMUSG IDs of all the kinase groups in NetPhorest Kinase names, kinase groups and ENSMUSG IDs of protein kinases were displayed in this table.

## Notes

### Competing Interest Statement

The authors have declared no competing interest.

### Summary of Updates

Experimental validations were conducted (Figs. 5J, K, and L, fig. S8). Changes of specific metabolic pathways were described (figs. S7, S9, and S10). Clinical significance was clarified (Fig. 8).

