## Supplementary material for "Structural robustness and temporal vulnerability of the starvation-responsive metabolic network in liver of healthy and obese mice": Supplemantary figures

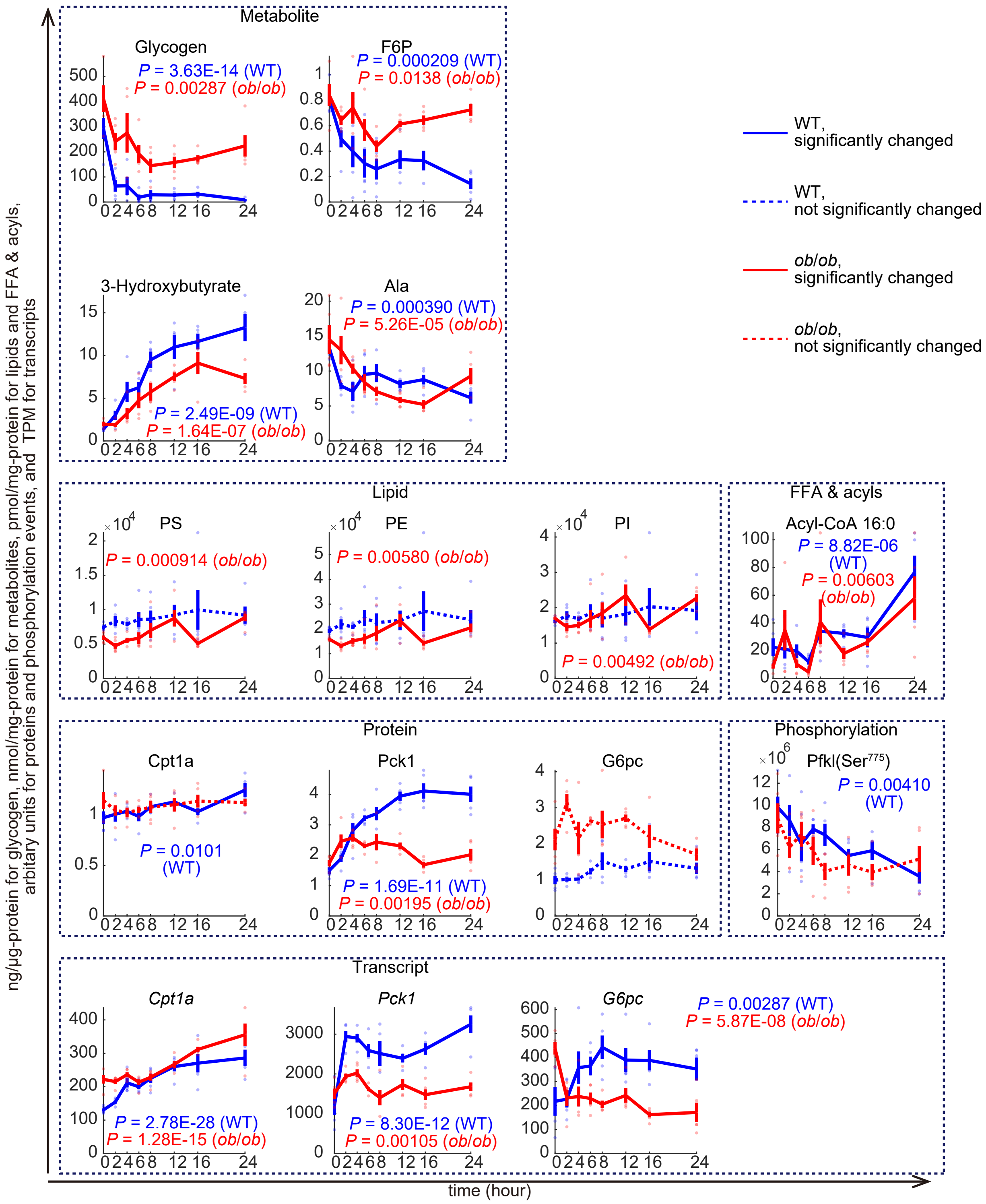


**fig. S1: Time courses of responsive molecules during starvation**
The x-axis represents the starvation time (hours) and the y-axis represents the concentration of the indicated molecule: ng/μg-protein for glycogen, nmol/mg-protein for other metabolites (F6P, Ala, 3-hydroxybutyrate); pmol/mg-protein for lipids (PS, PE, PI) and FFA & acyls (Acyl-CoA 16:0); arbitrary units for proteins (Cpt1a, Pck1, G6pc) and phosphorylation ($Ser^{775}$ of Pfkl). TPM for transcripts (*Cpt1a*, *Pck1*, *G6pc*). Blue lines are the responses to starvation in WT mice, and red lines are those in *ob*/*ob* mice. Solid lines represent that the responses were significant; dotted lines represent no significance. Data are shown as the mean and SEM. Dots represent the data from individual mice. *n* = 5 biological replicates per group. *P* values of one-way ANOVA of the molecules or ratio with significant changes are shown.

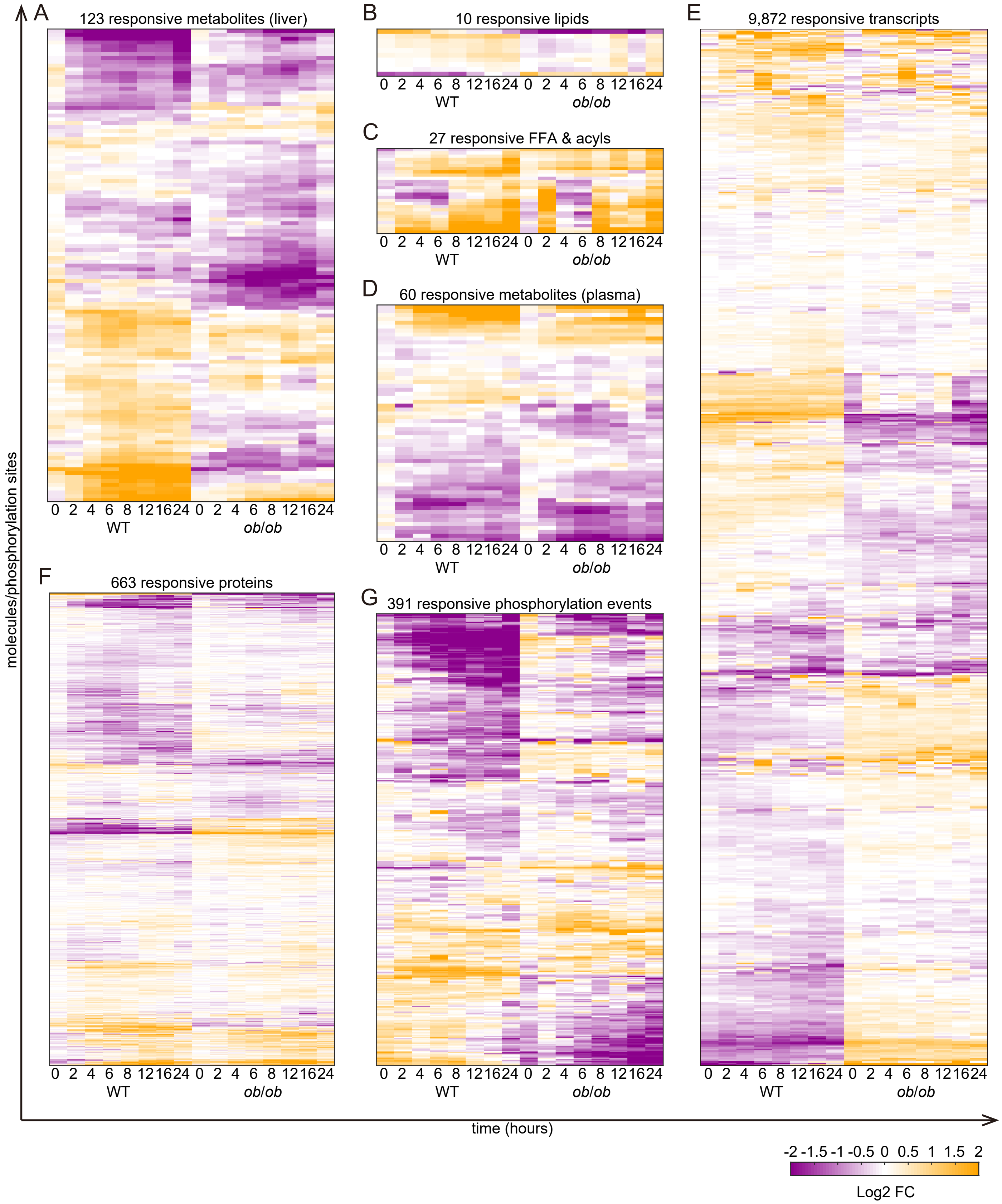


**fig. S2: Overview of each omic data**
The heatmaps of the time courses of 123 responsive metabolites (liver, A), 10 responsive lipids (B), 27 responsive FFA & acyls (C), 60 responsive metabolites (plasma, D), 9,872 responsive transcripts (E), 663 responsive proteins (F), and 391 responsive phosphorylation events (G) of WT and *ob*/*ob* mice during starvation. For heatmap, two time courses for each molecule were divided by the geometric mean of the values of WT and *ob*/*ob* mice before starvation (0 h) and then log2-transformed. The x-axis represents time for starvation, and the y-axis represents the molecules or phosphorylation events. *n* = 5 biological replicates per group.

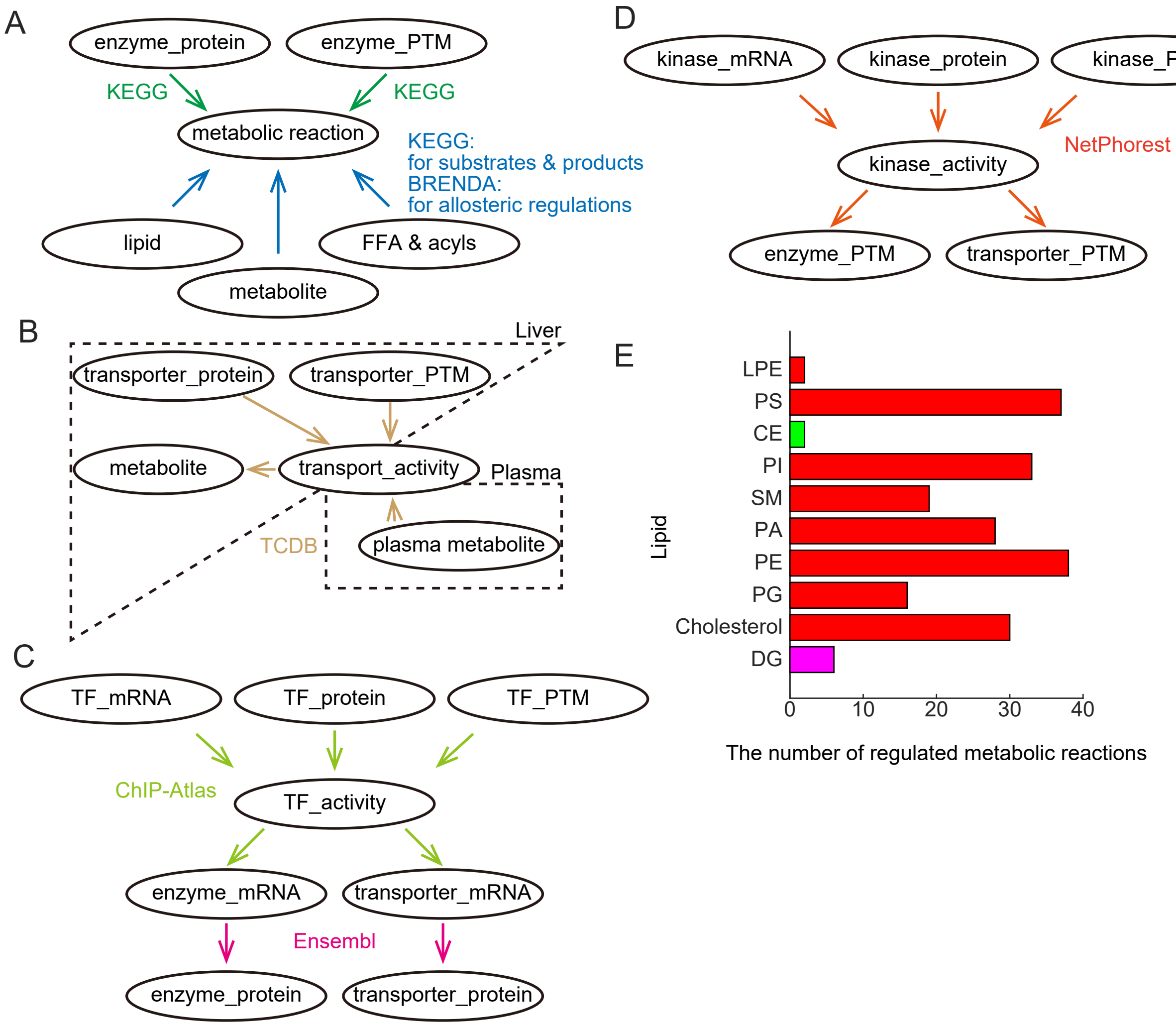


**fig. S3: Regulatory relationships in the starvation-responsive transomic network**

We assumed that the starvation-responsive molecules cooperatively regulate metabolic reactions (see also Fig. 1D).

**(**A) Metabolic network. The enzyme_protein (protein abundance of metabolic enzymes) and enzyme_PTM (phosphorylation events on metabolic enzymes) regulate metabolic reactions (identified with KEGG). Metabolites, lipids, and FFA & acyls regulate metabolic reactions as substrates, products (identified with KEGG), and allosteric regulators (identified with BRENDA). (B) Transport network identified with TCDB. The transporter_protein (protein abundance of transporters), transporter_PTM (phosphorylation events on transporters), and plasma metabolites regulate transport_activity (activities of transport). Transport_activity regulates metabolites in the liver, thereby regulating metabolic reactions in the liver. (C) Gene regulatory network identified with ChIP-Atlas and Ensembl. The TF_mRNA (mRNA abundance of TFs), TF_protein (protein abundance of TFs), and TF_PTM (phosphorylation events on TFs) regulate TF_activity (activities of TFs). TF_activity regulates enzyme_mRNA (mRNA abundance of metabolic enzymes) and transporter_mRNA (mRNA abundance of transporters). Enzyme_mRNA and transporter_mRNA regulate enzyme_protein and transporter_protein, respectively, and were aligned with Ensembl database. (D) Phosphorylation network identified with NetPhorest. The kinase_mRNA (mRNA abundance of kinases), kinase_protein (protein abundance of kinase), and kinase_PTM (phosphorylation events on kinases) regulate kinase_activity (activities of kinases). The kinase_activity regulates enzyme_PTM and transporter_PTM. (E) The number of metabolic reactions that lipids regulate as substrates, products, and/or allosteric regulators. The colors of the bars represent common responses (green), *ob*/*ob*-specific responses (red), and opposite responses (magenta) to starvation.

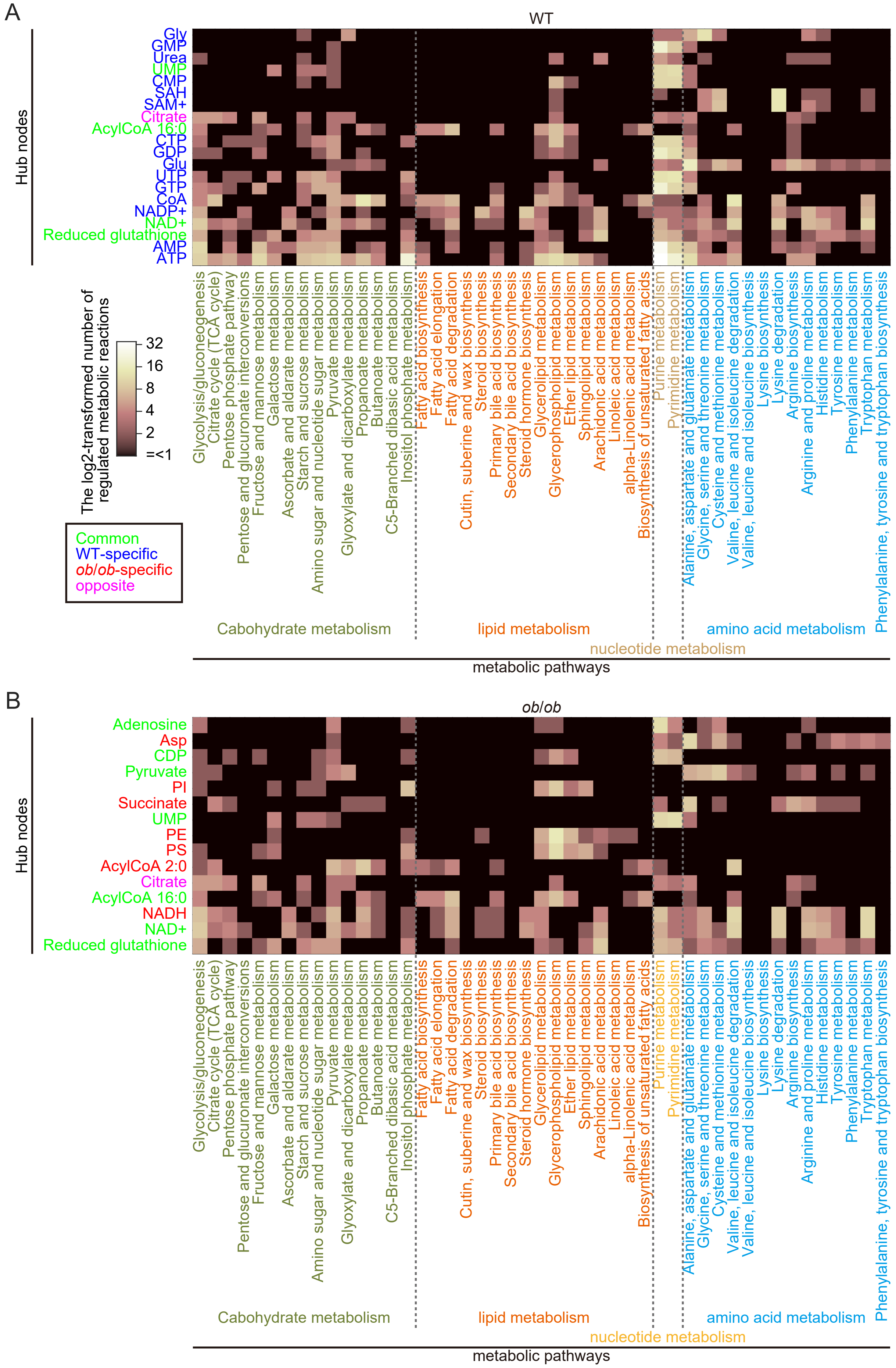


**fig. S4: Metabolic pathways regulated by the hub nodes**

Heatmaps representing the number of regulated metabolic reactions in each metabolic pathway (x-axis) regulated by the hub nodes (y-axis) in the starvation-responsive metabolic network in WT (A) and *ob*/*ob* mice (B). Colors of the labels of the hub nodes represent the responsiveness: green indicates common responses both in WT and *ob*/*ob* mice; blue, specific to WT mice; red, specific to *ob*/*ob* mice; magenta, opposite responses between WT and *ob*/*ob* mice. KEGG metabolic pathways (x-axis) in carbohydrate metabolism (olive green), lipid metabolism (orange), nucleotide metabolism (brown), and amino acid metabolism (light blue) are displayed.

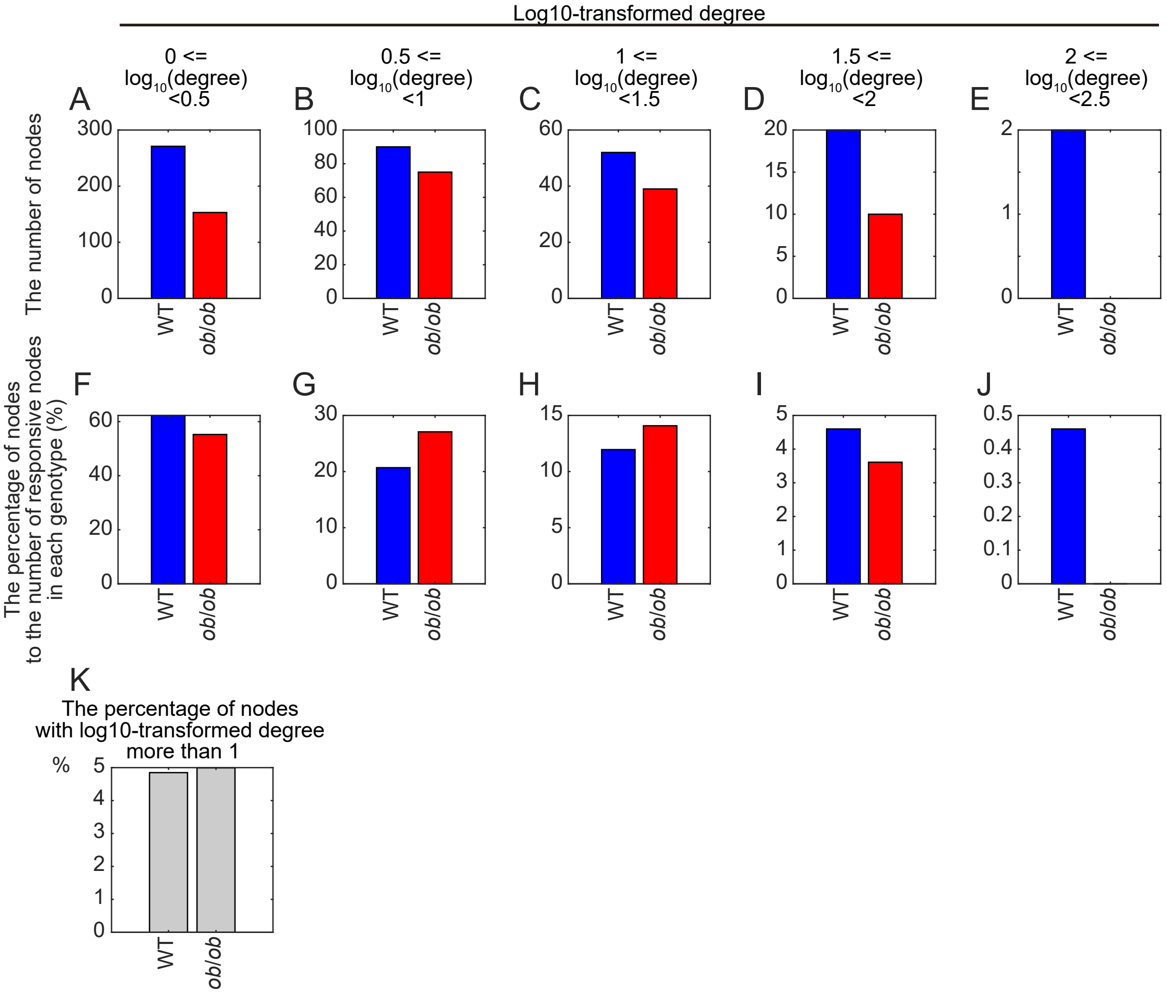


**fig. S5: Maintenance of the number of nodes with relatively large degrees in *ob*/*ob* mice**
The number of nodes (A-E) and the percentage of nodes to the number of responsive nodes in each genotype (F-J), whose log10-transformed degree are as indicated. (K) The percentages of nodes whose log10-transformed degree is more than 1.

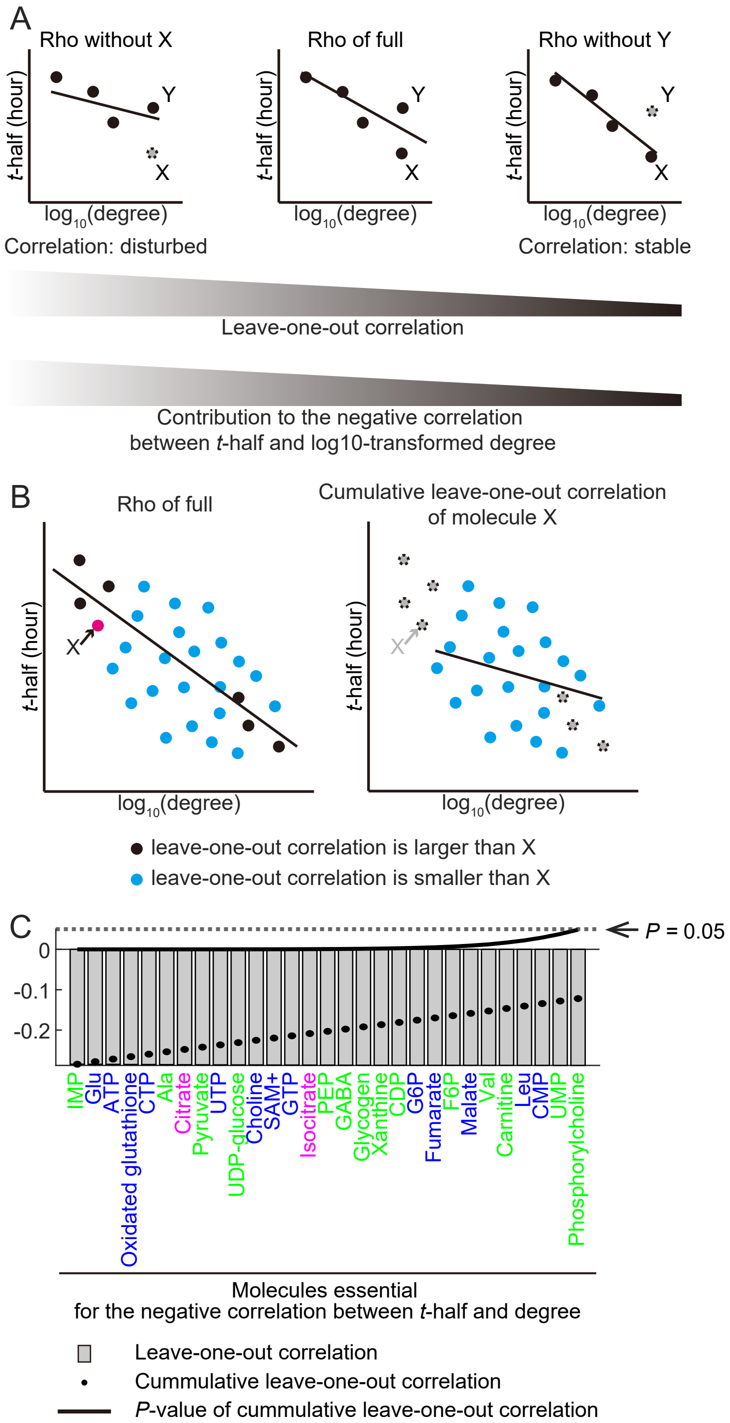


**fig. S6: Identification of molecules essential for the negative correlation between degree and *t*-half in WT mice**

(A) Schema for leave-one-out correlations. Log10-transformed degree negatively correlated with *t*-half in WT mice (center, “Rho of full”). The leave-one-out correlation of molecule X (left, “Rho without X”) was defined as correlation coefficient between log10-transformed degree and *t*-half that was calculated for all the molecules except for X. The larger leave-one-out correlation (left, “Rho without X”) represents the larger contribution to the negative correlation between log10-transformed degree and *t*-half. By contrast, the smaller leave-one-out correlation (right, “Rho without Y”) represents the smaller contribution to the negative correlation between log10-transformed degree and *t*-half. (B) The cumulative leave-one-out correlation of molecule X (right) was defined by the correlation coefficient between log10-transformed degree and *t*-half that was calculated for the molecules with leave-one-out correlation smaller than X (indicated in light blue). (C) Leave-one-out correlations (gray bars), cumulative leave-one-out correlations (black dots), and *P*-values of the cumulative leave-one-out correlations (black solid line) of the indicated molecules. Here, we displayed molecules with *P*-values of the cumulative leave-one-out correlations that were less than 0.05 and with leave-one-out correlations that were less than other molecules. *n* = 291 molecules in total.

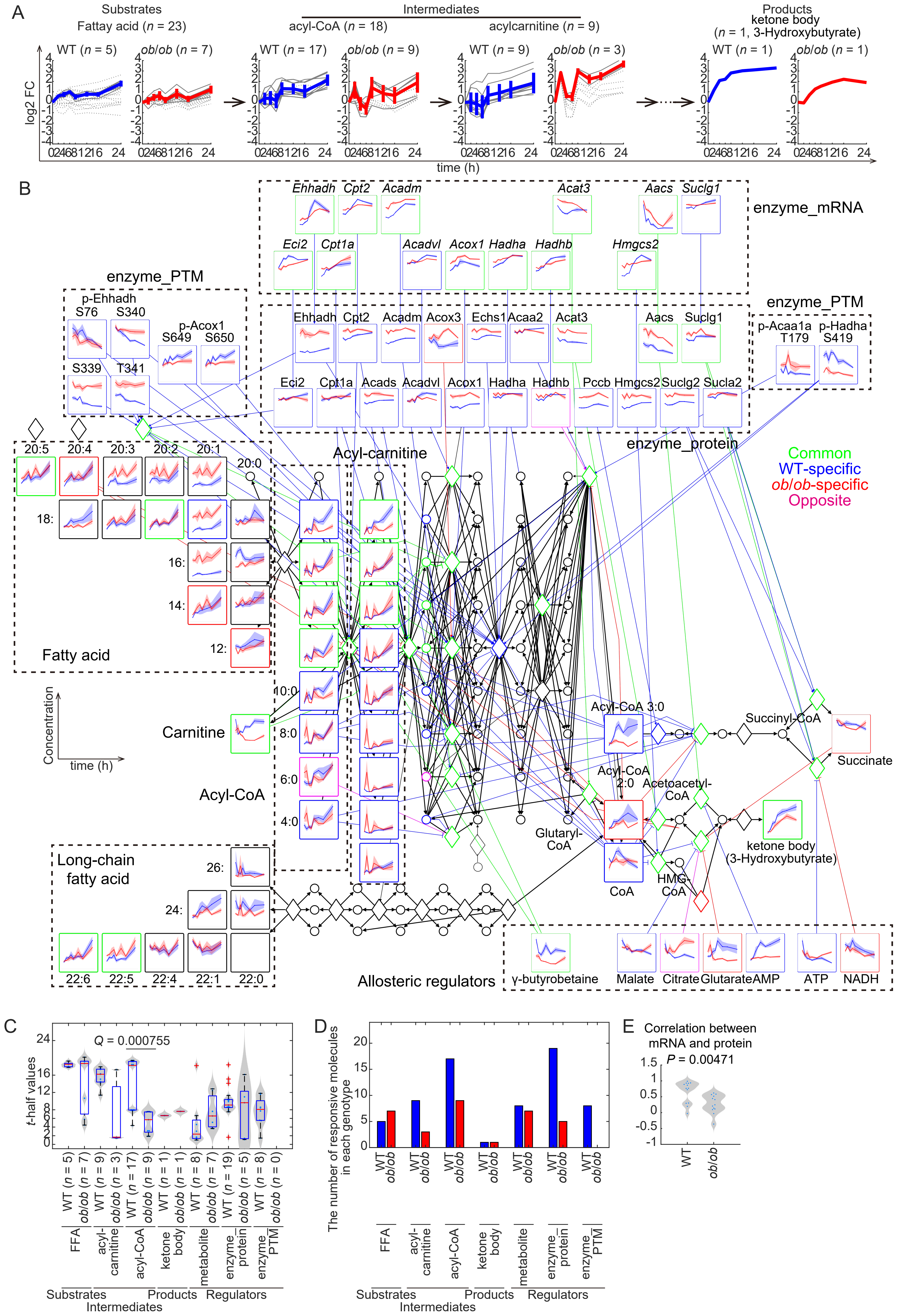

A
Substrates
Fattay acid (n = 23)
Intermediates
acyl-CoA (n = 18)
acylcarnitine (n = 9)
Products
ketone body
(n = 1, 3-Hydroxybutyrate)
WT (n = 5)
ob/ob (n = 7)
WT (n = 17)
ob/ob (n = 9)
WT (n = 9)
ob/ob (n = 3)
WT (n = 1)
ob/ob (n = 1)
log2 FC
time (h)
B
enzyme_mRNA
enzyme_PTM
enzyme_protein
Common
WT-specific
ob/ob-specific
Opposite
Acyl-carnitine
Fatty acid
Carnitine
Acyl-CoA
Long-chain fatty acid
Concentration
time (h)
Acyl-CoA 3:0
Succinyl-CoA
Succinate
Acyl-CoA 2:0
Acetoacetyl-CoA
Glutaryl-CoA
ketone body (3-Hydroxybutyrate)
CoA
HMG-CoA
Allosteric regulators
γ-butyrobetaine
Malate
Citrate
Glutarate
AMP
ATP
NADH
C
Q = 0.000755
t-half values
FFA
acyl-carnitine
acyl-CoA
ketone body
metabolite
enzyme_protein
enzyme_PTM
Substrates
Intermediates
Products
Regulators
D
The number of responsive molecules in each genotype
E
Correlation between mRNA and protein
P = 0.00471
WT
ob/ob

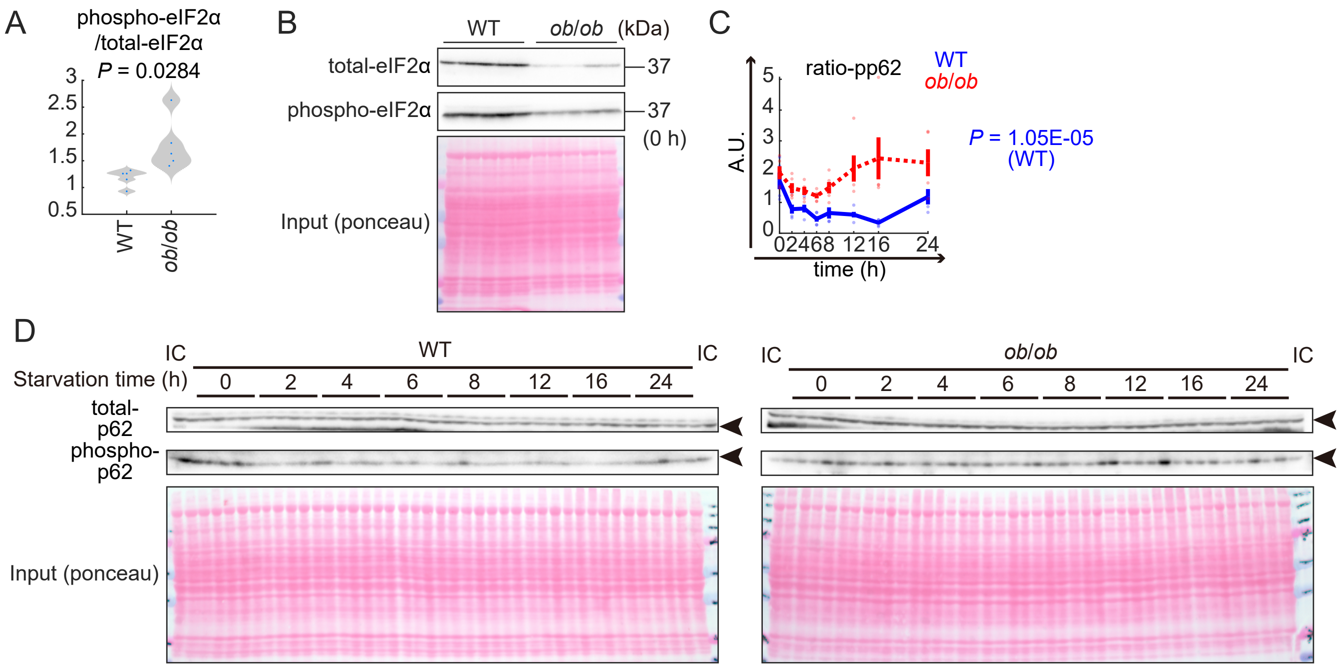


**fig. S8: Activities of protein translation and degradation in *ob*/*ob* mice**
(A) Phosphorylation ratio of eIF2α in WT and *ob*/*ob* mice before starvation. The x-axis represents genotype and the y-axis represents the ratio of phosphorylated eIF2α to total eIF2α. The *P*-value of a two-sample *t*-test comparing the phosphorylation ratios of eIF2α between WT and *ob*/*ob* mice is shown. (B) Images of western blotting for (A). (C) The time course of phosphorylation ratio of p62 in WT (blue line) and *ob*/*ob* mice (red line). The x-axis represents starvation time (h) and the y-axis represents ratio of phosphorylated p62 to total p62. The solid line represents that the response was significant; the dotted line represents no significance. Data are shown as the mean and SEM. Dots represent the data from individual mice. *P* value of one-way ANOVA of ratio-pp62 with significant changes is shown. (D) Images of western blotting for (C) at the indicated starvation time (h) of the indicated genotype. IC represents internal control to normalize the signals across membranes. Arrowheads indicate the total and phosphorylated p62 bands. *n* = 5 biological replicates per group.

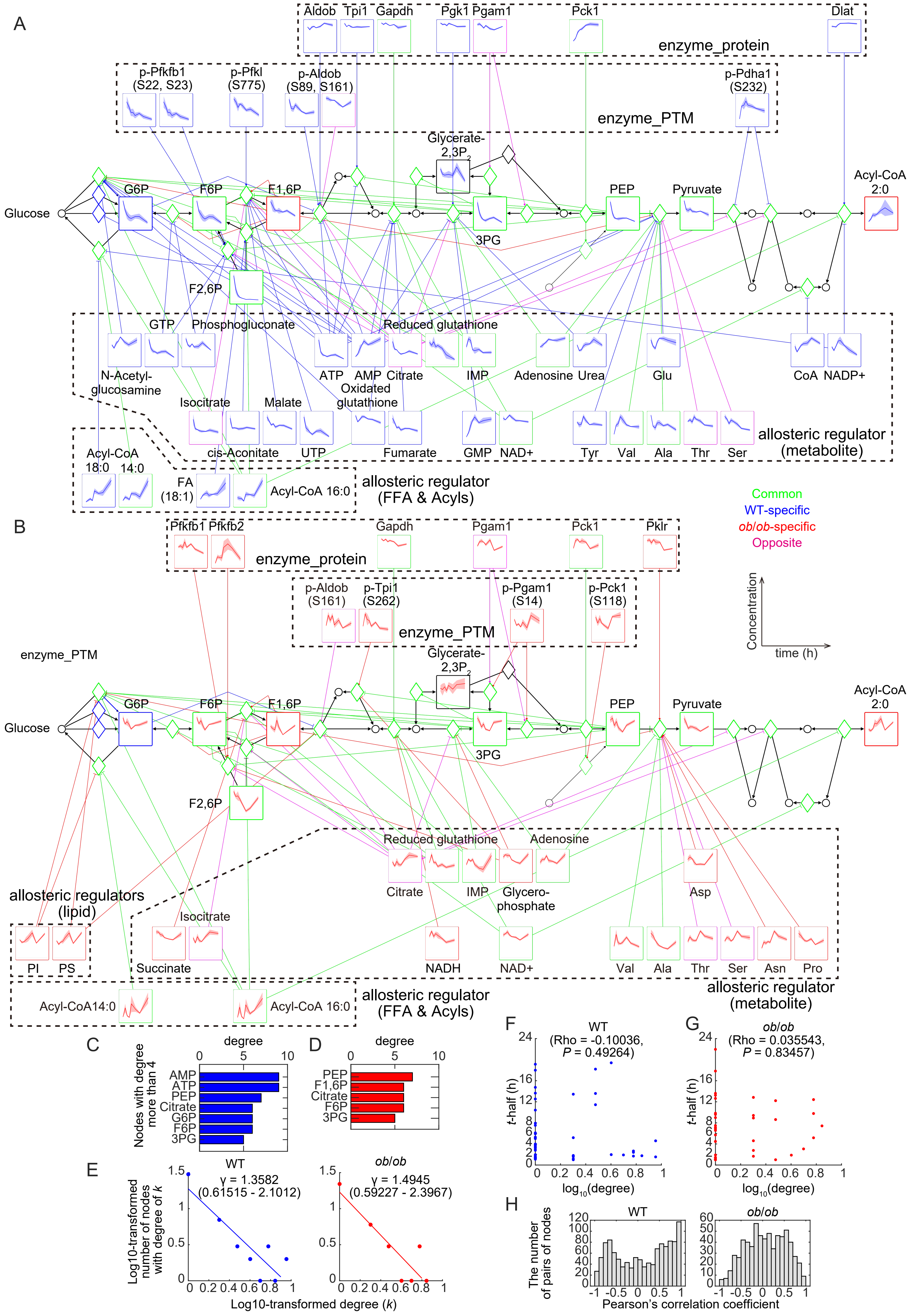

A
enzyme_protein
Aldob Tpi1 Gapdh Pgk1 Pgam1 Pck1 Dlat
enzyme_PTM
p-Pfkfb1 (S22, S23)
p-Pfkl (S775)
p-Aldob (S89, S161)
p-Pdha1 (S232)
Glycerate-2,3P2
Glucose
G6P
F6P
F1,6P
F2,6P
3PG
PEP
Pyruvate
Acyl-CoA 2:0
GTP
Phosphogluconate
Reduced glutathione
N-Acetyl-glucosamine
ATP
AMP
Citrate
Oxidated glutathione
IMP
Adenosine
Urea
Glu
CoA
NADP+
Isocitrate
Malate
cis-Aconitate
UTP
Fumarate
GMP
NAD+
Tyr
Val
Ala
Thr
Ser
allosteric regulator (metabolite)
Acyl-CoA 18:0 14:0
FA (18:1)
Acyl-CoA 16:0
allosteric regulator (FFA & Acyls)
Common
WT-specific
ob/ob-specific
Opposite
B
enzyme_protein
Pfkfb1 Pfkfb2 Gapdh Pgam1 Pck1 Pklr
enzyme_PTM
p-Aldob (S161)
p-Tpi1 (S262)
p-Pgam1 (S14)
p-Pck1 (S118)
Concentration
time (h)
enzyme_PTM
Glycerate-2,3P2
Glucose
G6P
F6P
F1,6P
F2,6P
3PG
PEP
Pyruvate
Acyl-CoA 2:0
Reduced glutathione
Adenosine
Citrate
IMP
Glycero-phosphate
Asp
allosteric regulators (lipid)
PI
PS
Isocitrate
Succinate
NADH
NAD+
Val
Ala
Thr
Ser
Asn
Pro
allosteric regulator (metabolite)
Acyl-CoA14:0
Acyl-CoA 16:0
allosteric regulator (FFA & Acyls)
C
degree
0 5 10
Nodes with degree more than 4
AMP
ATP
PEP
Citrate
G6P
F6P
3PG
D
degree
0 5 10
PEP
F1,6P
Citrate
F6P
3PG
E
WT
γ = 1.3582
(0.61515 - 2.1012)
ob/ob
γ = 1.4945
(0.59227 - 2.3967)
Log10-transformed number of nodes with degree of k
Log10-transformed degree (k)
0 0.5 1 1.5
0 0.2 0.4 0.6 0.8 1
F
WT
(Rho = -0.10036, P = 0.49264)
t-half (h)
log10(degree)
0 2 4 6 8 12 16 24
0 0.2 0.4 0.6 0.8 1
G
ob/ob
(Rho = 0.035543, P = 0.83457)
t-half (h)
log10(degree)
H
WT
ob/ob
The number of pairs of nodes
Pearson's correlation coefficient
0 20 40 60 80 100 120
0 10 20 30 40 50 60
-1 -0.5 0 0.5 1

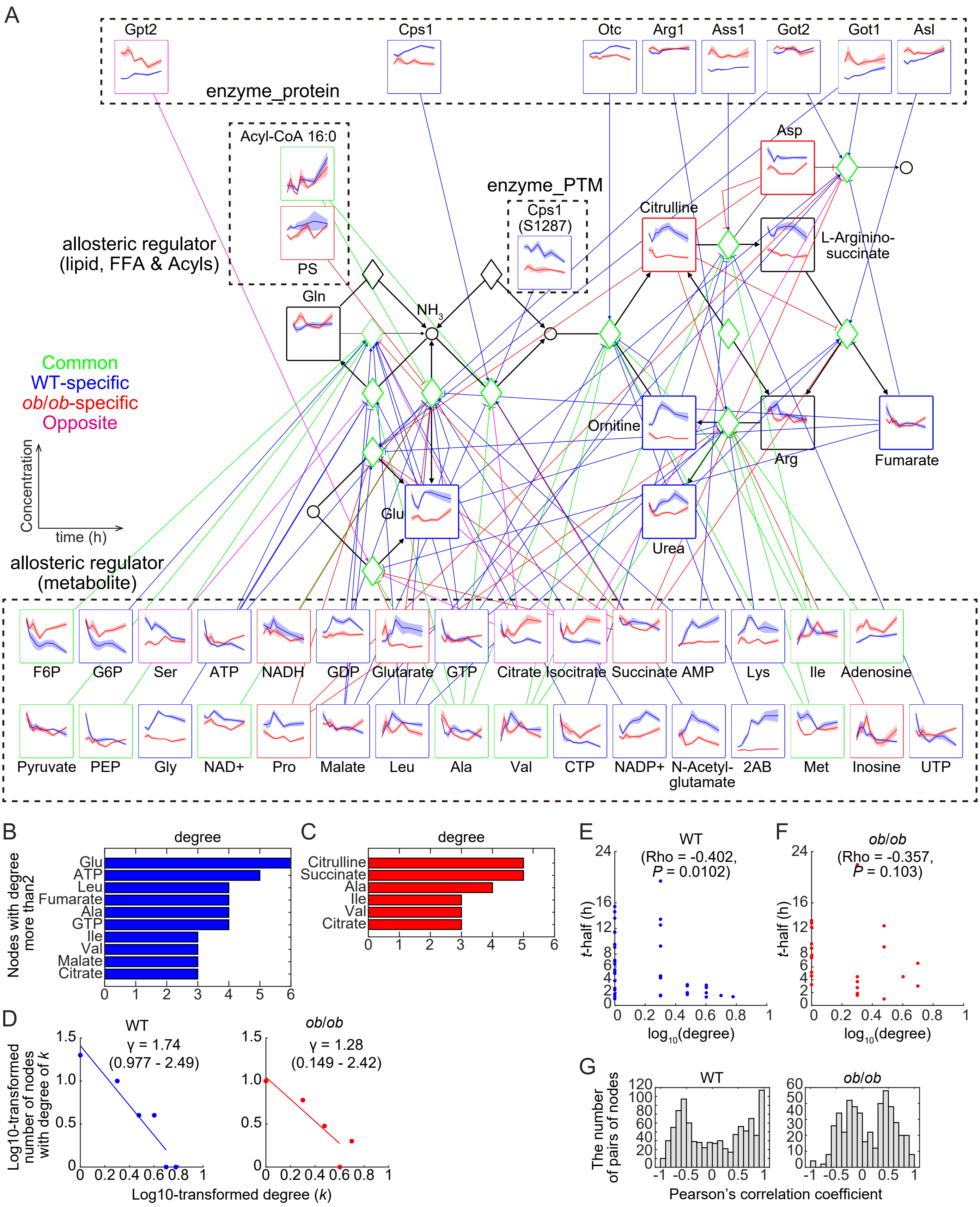
A
enzyme_protein
Gpt2
Cps1
Otc
Arg1
Ass1
Got2
Got1
Asl
Acyl-CoA 16:0
PS
allosteric regulator
(lipid, FFA & Acyls)
enzyme_PTM
Cps1
(S1287)
Asp
Citrulline
L-Arginino-
succinate
Gln
NH3
Ornitine
Arg
Fumarate
Glu
Urea
Common
WT-specific
ob/ob-specific
Opposite
Concentration
time (h)
allosteric regulator
(metabolite)
F6P
G6P
Ser
ATP
NADH
GDP
Glutarate
GTP
Citrate
Isocitrate
Succinate
AMP
Lys
Ile
Adenosine
Pyruvate
PEP
Gly
NAD+
Pro
Malate
Leu
Ala
Val
CTP
NADP+
N-Acetyl-
glutamate
2AB
Met
Inosine
UTP
B
degree
Nodes with degree
more than2
Glu
ATP
Leu
Fumarate
Ala
GTP
Ile
Val
Malate
Citrate
0 1 2 3 4 5 6
C
degree
Citrulline
Succinate
Ala
Ile
Val
Citrate
0 1 2 3 4 5 6
D
WT
γ = 1.74
(0.977 - 2.49)
ob/ob
γ = 1.28
(0.149 - 2.42)
Log10-transformed
number of nodes
with degree of k
Log10-transformed degree (k)
E
WT
(Rho = -0.402,
P = 0.0102)
t-half (h)
log10(degree)
F
ob/ob
(Rho = -0.357,
P = 0.103)
t-half (h)
log10(degree)
G
WT
ob/ob
The number
of pairs of nodes
Pearson's correlation coefficient

**fig. S10: The starvation-responsive metabolic network for starvation-responsive metabolic reactions in urea cycle**

The starvation-responsive metabolic network for starvation-responsive metabolic reactions in urea cycle, that were manually constructed based on "Arginine biosynthesis" (mmu00220) in the KEGG database (*55*–*57*) (**A**). Time courses of measured metabolites, lipids, FFA & acyls, enzyme_protein, and enzyme_PTM are shown for corresponding nodes as the mean and SEM. The colors of the frames represent common responses (green), WT-specific responses (blue), *ob*/*ob*-specific responses (red), and opposite responses (magenta) to starvation. The black frames indicate that were not responsive to starvation. From metabolites, lipids, and FFA & acyls to metabolic reactions, only allosteric regulatory connections are colored. (**B** and **C**) The degree of the indicated nodes in WT (**B**) and in *ob*/*ob* (**C**) mice. Nodes with degree more than 2 are shown. The x-axis represents degree (the number of edges) and the y-axis represents nodes. It should be noted that the degrees shown in **B** and **C** are not necessarily identical to those shown in **A.** This is because if a molecule regulates a metabolic reaction both as a substrate or product, and as an allosteric regulator, the edges were counted as a single edge in **B** and **C**, but were shown separately in **A**. (**D**) Degree distributions of the nodes in the starvation-responsive metabolic network of urea cycle in WT (blue) and *ob*/*ob* (red) mice. The x-axis represents the log10-transformed degree ($k$); the y-axis represents the log10-transformed number of nodes with degree of $k$. Lines show the linear regression of the degree distributions. The slope of the lines ($\gamma$), which is defined as a scaling parameter, and the 95% credible interval are shown. $n$ = 40 molecules for WT mice and $n$ = 22 molecules for *ob*/*ob* mice. (**E** and **F**) Scatter plots for the log10-transformed degree (x-axis) and *t*-half (y-axis) in WT (**E**) and *ob*/*ob*

(**F**) mice. Dots represent the molecules (including phosphorylation events) of the starvation-responsive metabolic network in urea cycle. Blue dots represent WT and red dots represents *ob*/*ob* mice. Spearman's correlation coefficients (Rho) and *P* values are shown. $n$ = 40 molecules for WT mice and $n$ = 22 molecules for *ob*/*ob* mice. (**G**) Distribution of the pairwise Pearson's correlation coefficients among time courses of responsive molecules in the starvation-responsive metabolic network of urea cycle in WT and *ob*/*ob* mice. The x-axis represents correlation coefficients; the y-axis represents the number of pairs of molecules examined for correlation. $n$ = 40 molecules for WT mice and $n$ = 22 molecules for *ob*/*ob* mice.

A

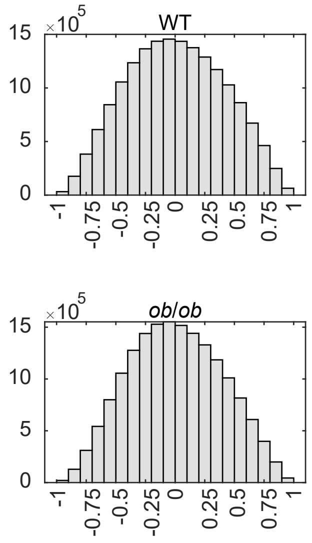


**fig. S11: Distribution of the pairwise Pearson' s correlation coefficients among time courses of all the measured metabolites, lipids, FFA & acyls, enzyme_protein, and enzyme_PTM**

The x-axis represents the pairwise Pearson' s correlation coefficients among time courses of all the measured metabolites, lipids, FFA & acyls, enzyme_protein and enzyme_PTM; the y-axis represents the number of pairs of molecules examined for correlation. The data of WT (A) and *ob*/*ob* (B) mice are shown. *n* = 5,866 molecules in each genotype.

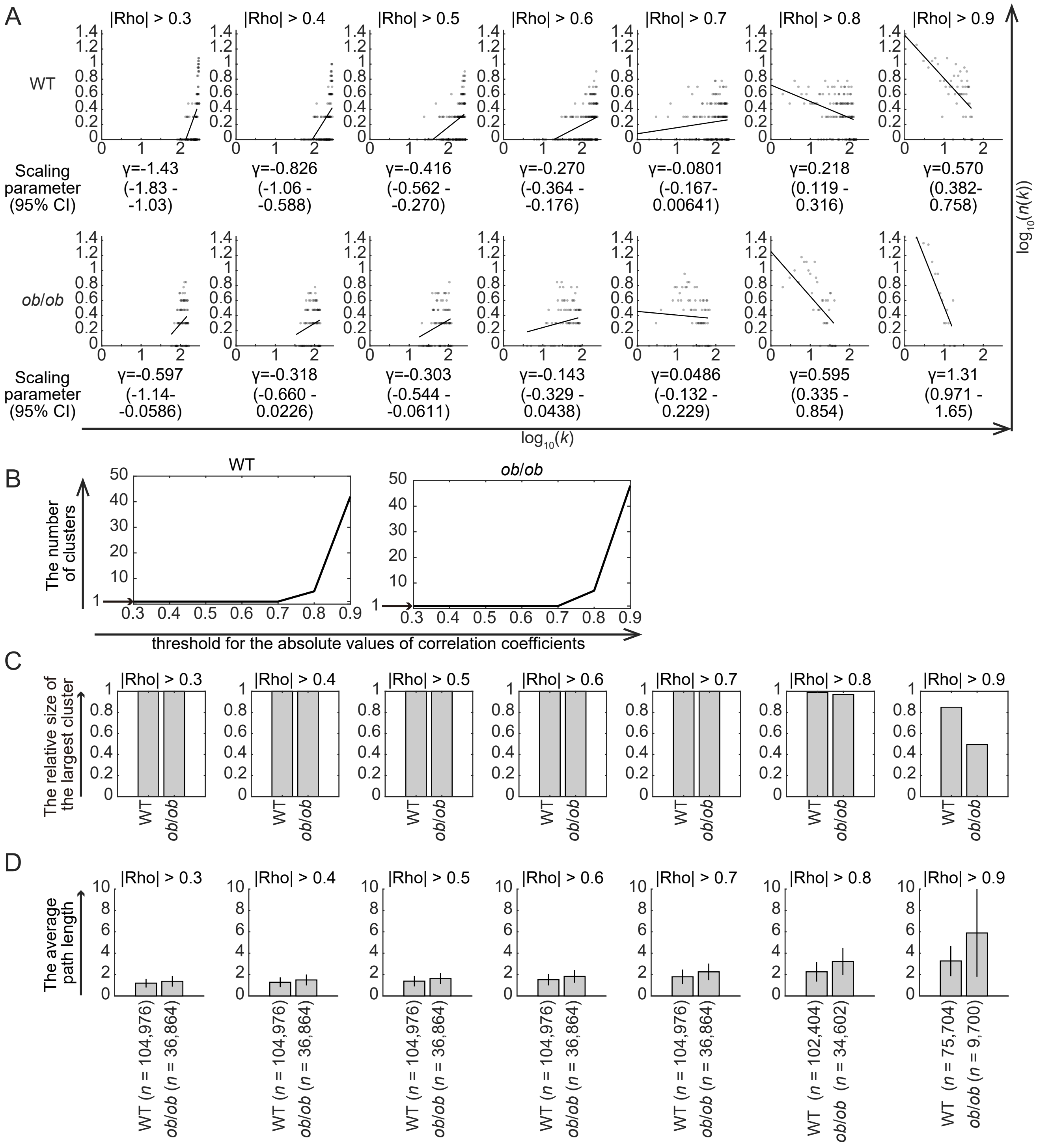


**Fig. S12: The correlation metabolic network during starvation**
(A) Scatter plots showing the degree distributions of correlation metabolic network corresponding to each threshold of absolute values of correlation coefficients (|Rho|) ranging from 0.3 to 0.9. The x-axis represents $\log_{10}(k)$, the log10-transformed degree; the y-axis represents $\log_{10}(n(k))$, the log10-transformed number of nodes with degree of *k*. Dots denote the degree distribution of the indicated networks and lines show the linear-regression of the degree distribution. The scaling parameters ($\gamma$) and the 95% credible intervals are shown. Data for WT and *ob*/*ob* mice are shown. The number of clusters (B), relative size of the largest cluster (C) and the average path length (D) for each threshold are also shown. The lower threshold than 0.8 for the absolute value of correlation coefficients resulted in formation of a single cluster (B). The higher threshold (0.8 or 0.9) for the absolute value of correlation coefficients resulted in more scale-free-like networks. To identify the qualitative differences between WT and *ob*/*ob* mice clearly, we used 0.9 as a threshold for the absolute value of correlation coefficients in Fig. 7**D** and **E**. Using 0.9 as a threshold for the absolute value of correlation coefficients, the relative size of the largest cluster was smaller (C) and the average path length was larger in *ob*/*ob* mice than in WT mice (D). For average path length, the numbers of the investigated paths are shown in the parentheses following to the genotype label. The error bars show standard deviation.
